# Nanopore sequencing of antibiotic-resistant *Klebsiella pneumoniae* JRCGR1 isolate from Pakistan and country scale pan-genomics

**DOI:** 10.1101/2023.03.05.531167

**Authors:** Zarrin Basharat, Zaib un Nisa, Muhammad Irfan, Rao Muhammad Abid Khan, Asad Karim, Muhammad Aurongzeb, Syed Shah Hassan

**Author notes:** Corresponding author email: Z.B.; S.S.H.

## Abstract

*Klebsiella pneumoniae* is a gram-negative, encapsulated, non-motile bacterium that can cause damage to human lungs. Antibiotic resistance of this strain requires constant monitoring and for this purpose, Oxford nanopore mediated whole genome sequencing was done for a strain acquired from the pleural fluid of a diabetic patient suffering from chronic kidney disease. A genome of 5.98 MB was obtained, with 8,869 CDSs and 110 RNAs. Around 96 proteins were seen as involved in virulence, disease or defense, while 25 antibiotic resistance genes were identified. Pan-genome was inferred as open, with 12,857 and 1,303 CDSs forming respective accessory and core genome of antibiotic resistant Pakistani strains (n=168) of this species. Apart from notable antimicrobial resistance genes like beta-lactamases, a gene conferring resistance to colistin was also mined. To the best of our knowledge, this is first report of nanopore mediated long read sequencing of a hypermucoviscous virulent *K. pneumoniae* from Pakistan.

## 1. Introduction

*Klebsiella pneumoniae* mostly affects immunocompromised or the elderly people [1,2]. The patient populace is assumed to have a compromised respiratory defense system, including people having chronic obstructive pulmonary diseases, diabetes, kidney failure, liver disease, etc [3]. Most infections are attained when a person is hospitalized for some other disease (a nosocomial infection) [4]. Along with pneumonia, *Klebsiella* can also lead to biliary tract, urinary tract and infections at the surgical wound sites [5]. The additional array of clinical ailments comprise meningitis, cholecystitis, diarrhea, pneumonia, bacteremia, thrombophlebitis, upper respiratory tract infection, wound infection, osteomyelitis and sepsis [6].

The most frequent illness instigated by *Klebsiella* outside the hospital is pneumonia, normally manifested as bronchopneumonia or bronchitis [7]. Such patients have an amplified risk of lung abscess development, empyema, cavitation, or pleural adhesion formation. The death rate in such cases is approximately 50%, even upon administration of antimicrobial therapy[8]. Treatment of *K. pneumoniae* includes antibiotics (like cephalosporins and aminoglycosides etc) [9]. The choice of treatment medicine, as well as regimen, is conditional to the person’s medical history, health state and disease severity. Undue antibiotic usage habits can influence and lead to a heightened risk of *Klebsiella* spp. nosocomial infections. *K. pneumoniae* strains usually carry beta-lactamases, which make it ampicillin resistant [10]. It has been noted that several strains have acquired an extended-spectrum beta-lactamase and plasmid mediated quinolone resistance[11]. Further resistance to antibiotics like amoxicillin, carbenicillin, and ceftazidime has also been noted [12,13]. It is worrisome that colistin-resistant strains of *K. pneumoniae* have also been reported in the literature [14,15]. Isolates of *K. pneumoniae* carrying the New Delhi Metallo beta-lactamase (NDM-1) gene, which confers resistance to intravenous antibiotic carbapenem, were reported in 2009 in the neighboring countries of India and Pakistan [16,17].

In this study, we set to study the whole genome sequence-based antibiotic resistance profile of a clinical isolate of the *K. pneumoniae* using state-of-the-art long read technology i.e. Oxford Nanopore sequencing. Detailed information regarding resistant genes was inferred and comparsion to other *Klebsiella pneumoniae* strain data from Pakistan was made using pooled genome data, to comprehensively study the pan-genome.

## 2. Material and methods

### 2.1. Sample collection and DNA extraction

To study the drug-resistant *K. pneumoniae* genome, the strain was obtained from the Civil Hospital laboratory in Karachi, Pakistan. Written informed consent was obtained from the patient and study was approved by the ethics committee of the ICCBS, University of Karachi (Reference no: ICCBS/IEC-070-HUS-2021/Protocol/1.0). The strain was isolated from the pleural fluid of a diabetic patient with chronic kidney disease. Ring testing confirmed that it was a hypermucoid strain.

### 2.2. Minimum inhibitory concentration (MIC)

MIC was also studied according to the Performance Standards for Antimicrobial Susceptibility Testing, Clinical and Laboratory Standards Institute (CLSI), Supplement 100[18]. Briefly, resistance was measured for the antibiotics (mentioned in Table 1) by two-fold broth microdilution. For this purpose, JRCGR1 strain was cultured overnight and diluted in sterilized phosphate buffered saline to ~105 CFU/mL. Each antibiotic was prepared at varying concentrations via two-fold serial dilutions in Mueller-Hinton broth. Aliquots of 5 μL of JRCGR1 suspension were then conveyed to 96-well microtiter plate with 100 μL of prepared antibiotic solution of different concentration. These plates were incubated at 37°C and the MIC values were recorded after 16 hrs.

**Table 1.** MICs obtained by broth dilution method.

| **Serial no.** | **Antibiotic** | **MIC (mg/L)** |
| --- | --- | --- |
| 1. | Ceftriaxone | > 4 |
| 2. | Ciprofloxacin | > 1 |
| 3. | Amikacin | >64 |
| 4. | Tazobactam | 16 |
| 5. | Piperacillin | 32 |
| 6. | Sulbactam | 64 |
| 7. | Sulphamethoxazole | 0.05 |
| 8. | Trimethoprim | >0.05 |
| 9. | Nitrofurantoin | >64 |
| 10. | Fosfomycin | >128 |
| 11. | Gentamycin | >8 |

### 2.3. DNA extraction

For DNA extraction, the isolate was initially cultured on the nutrient agar plate and the colonies were then picked and mixed with nuclease free water to form suspension. Centrifugation (3,000 g for 15 min) was done and supernatant was removed without disturbing the sediment. DNA was extracted by using commercial DNA extraction kit (QIAamp DNA-Mini Kit; (Qiagen^®^). The Qubit^™^ dsDNA BR Assay kit (Invitrogen) was used to calculate the amount of DNA with the Qubit^®^2.0 Fluorometer (Invitrogen), according to the supplier’s instructions.

### 2.4. Library preparation and genome sequencing

Genomic DNA Ligation sequencing method was followed for library preparation (using kit# SQK-LSK109, according to the instructions at: https://community.nanoporetech.com/protocols/gDNA-sqk-lsk109; <u>accessed 25 Feb, 2022</u>). Prepared library was primed and loaded on flow cell. Sequencing Buffer (SQB), Loading Beads (LB), Flush Tether (FLT) and one tube of Flush Buffer (FB) were thawed at room temperature and mixed. MinION Mk1B lid was removed and the flow cell slid under the clip. 30 μl of thawed and mixed Flush Tether (FLT) was added directly to the tube of thawed and mixed Flush Buffer (FB) and mixed by vortexing at room temperature. 800 μl of the priming mix was loaded into the flow cell via the priming port, evading air bubbles formation. This was left for 5 minutes and in the course of this time, the library was prepared for loading, by thoroughly mixing the contents of the Loading Beads (LB) by pipetting. 37.5 μl Sequencing Buffer, 25.5 μl Loading Beads, and 12 μl DNA library. 200 μl of the priming mix was then put into the flow cell, through the priming port. Then, 75 μl of the sample was added through the SpotON sample port in a dropwise fashion. Ports were closed and sequencing started by clicking the sequencing option in MinKNOW software. After initial data generation, base calling was done.

### 2.5. Genome assembly and annotation

Whole genome assembly was done using unicycler with the parameters: single_end_libs, Reads: combined.fastq.gz, platform nanopore, min_contig_len: 300, pilon iteration:2, minimum contig coverage of 5, and racon iteration:2 [19]. Genome annotation was done using RAST server [20]. Plasmid sequence was inferred using PlasmidFinder-2.0 (https://cge.cbs.dtu.dk/services/PlasmidFinder/; <u>accessed 28 February, 2022</u>). Detailed annotation of plasmids was done using Plannotate (retrieved from http://plannotate.barricklab.org/ on 7 Dec, 2022). This tool queries the submitted sequence against numerous databases (genoLIB, FPBase, Rfam), with a match cut-off of 95%, using methods like BLAST, DIAMOND, and Infernal. For matching Swiss-Prot, the score is ≥50% match. Multi-Locus Sequence Typing (MLST) was done using the Center for Genomic Epidemiology website (https://cge.cbs.dtu.dk/; <u>accessed on 28 Feburary, 2022</u>). Some parameters were determined from Pathogenwatch (https://pathogen.watch/). Phage regions were mapped using PHASTER (https://phaster.ca/) and CRISPR-Cas sequences were matched using CRISPRCasFinder (https://crisprcas.i2bc.paris-saclay.fr/CrisprCasFinder/). Virulence genes were predicted using BLAST against the VFDB (http://www.mgc.ac.cn/VFs/). Antimicrobial resistance genes were predicted using the Comprehensive Antibiotic Resistance Database (CARD) (https://card.mcmaster.ca/home).

### 2.6. Pan-genome and resistome analysis

For comparative analysis with other *K. pneumoniae* Pakistani isolates, whole gneome data of *K. pneumoniae* isolates (n=168) was obtained from PATRIC database on 12 March, 2022 (currently https://bv-brc.org). Pan-genome analysis was done using the BPGA software [21]. The clustering of homologous genes was done using USEARCH clustering algorithm, with a 70% cut-off value. The .fasta files were taken as input. Genomes were aligned using MUSCLE tool. Phylogeny was built using the UPGMA method and functional annotation was done using Cluster of Orthologous Group (COG). This was to catalog and arrange the homologous genes. Comprehensive Antibiotic Resistance Database (CARD) was utilized for the resistome mapping of the genome fractions (pan, core, and unique) [22]. The selection standard for the mined homologs was kept as perfect and strict hits only.

## 3. Results and Discussion

### 3.1. Characteristics of JRCGR1

*K. pneumoniae* JRCGR1 was characterized as a hypermucoviscous virulent strain, with phenotypic resistance to several antibiotics. In comparison to typical opportunistic *K. pneumoniae* strains, hypermucoviscous/hypervirulent strains lead to severe infections, which may be life-threatening even for the formerly healthy people of the community. Numerous research outcomes have reported that gene cross-over of multidrug resistant and hypermucoviscous strains may lead to the development of a highly contagious and drug-resistant bacterium [23]. MIC profile showed that it was resistant to several antibiotics (Table 1).

### 3.2. Whole genome analysis

Since WGS is being progressively applied in analyzing bacteria clones, studying antimicrobial resistance and tracking pathogen population structure, we utilized long-read sequencing technology Oxford nanopore for the sequence determination of our isolate. Nanopore sequencing is a cost and time-effective approach that is portable and can sequence without doing PCR amplification[24,25]. Since it displays results in real time and has the potential to be used in mobile testing, it has the potential to be embedded in national surveillance and testing frameworks in case of outbreaks of pathogens. Additionally, it provides long reads which facilitate better assemblies of the pathogenic genomes[26,27].

For strain JRCGR1, with coverage of 19x, a genome of 5.7 MB was assembled. The final assembly consisted of 6 contigs. Two contigs were ≥ 25000 bp while 3 contigs were ≥ 50,000 bp. The largest contig was 5.4 MB. Average long read coverage was 114.6 while N50 was 5410184 bp. GC (%) was 56.68. Around 8,869 CDSs and 110 RNAs were obtained. MLST validated the sequence as *K. pneumoniae* (Table 2) via alignment and coverage of alleles. It is an unequivocal technique for typification of bacterial isolates at a species scale, using the internal fragment sequences of seven housekeeping genes. No allele hit was obtained for mdh locus, while partial match for gapA locus was obtained, thus the nearest sequence type (ST) for isolate *K. pneumoniae* JRCGR1 was identified as 2096 or 3610, using MLST server. Pasteur institute scheme was used for validation and 2096 was obtained from its server as ST of *K. pneumoniae* JRCGR1. This ST has previously been identified in the hypervirulent *K. pneumoniae* from India [28] but not reported from Pakistan. Capsule K matched KL2 at Pathogenwatch but it did not have a good confidence score. Virulence analysis from Kleborate showed a score of 3, and included aerobactin (*iuc1*) and/or salmochelin without yersiniabactin/colibactin). Russo et al. have reported that aerobactin, a citry1-hydroxamate siderophore arbitrates virulence in hypervirulent *K. pneumoniae* and leads to a boost in the production of siderophore under iron-restrictive conditions[29].

**Table 2.** MLST statistics of the sequenced genome.

| Locus | Identity | Coverage | Alignment<br>Length | Allele<br>Length | Gaps | Allele |
| --- | --- | --- | --- | --- | --- | --- |
| gapA | 99.77778 | 99.77778 | 450 | 450 | 1 | gapA_1 |
| infB | 100 | 100 | 318 | 318 | 0 | infB_6 |
| mdh | 99.79036 | 99.79036 | 477 | 477 | 1 | mdh_1 |
| pgi | 100 | 100 | 432 | 432 | 0 | pgi_1 |
| phoE | 100 | 100 | 420 | 420 | 0 | phoE_384 |
| rpoB | 100 | 100 | 501 | 501 | 0 | rpoB_46 |
| tonB | 100 | 100 | 414 | 414 | 0 | tonB_1 |

In total, four plasmid fragments were present (Table 3) and only contigs 1 and 3 referred completely to chromosomal DNA. Due to low coverage of the whole genome, we could not capture complete circular plasmid genome sequences. However, we describe the characteristics of the mentioned plasmids as the fragments mean that these genes would be hosted by the strain JRCGR1 as well, even though not captured by 19x nanopore genome. blaOXA-232 carbapenemase-encoding ColKP3 plasmid has been identified previously in the multidrug-resistant strains of ST16/ST231 *K. pneumoniae* and *Escherichia coli* from Thailand [30]. This plasmid carries mobilization proteins, a replication protein, and a class D beta-lactamase blaOXA-181. ColRNAI has previously been noted as a mobilizable plasmid in *K. pneumoniae*.

**Table 3.** Plasmid information for *K. pneumoniae* JRCGR1.

| Plasmid | Identity | Query /<br>Template<br>length | Contig | Position in<br>contig | Accession of<br>matching<br>plasmid |
| --- | --- | --- | --- | --- | --- |
| ColKP3 | 100 | 280 / 280 | 5 | 8751..9030 | JN205800 |
| ColKP3 | 100 | 280 / 280 | 5 | 2624..2903 | JN205800 |
| ColRNAI | 96.95 | 131 / 130 | 6 | 8217..8347 | DQ298019 |
| IncFIB(pNDM-Mar) | 99.54 | 439 / 439 | 2 | 101585..102023 | JN420336 |
| IncHI1B(pNDM-MAR) | 100 | 570 / 570 | 2 | 138477..139046 | JN420336 |
| IncHI1B(pNDM-MAR) | 100 | 570 / 570 | 2 | 86..655 | JN420336 |

It usually carries a lysis precursor unit rom, immunity gene similar to that of cloacin, exclusion and colicin-like protein-encoding genes. The unit rom is known to negatively impact transcription while immunity genes encode bacteriocins, which confer protection.Mob proteins are known for replication relaxation and mob also carries VirD4 region, part of the type IV secretory pathway. IncFIB(pNDM-Mar) and IncHI1B(pNDM-MAR) encodes insertion sequences, transposases, resolvases, beta-lactamase OXA-1, aminoglycoside N(6’)-acetyltransferase, Metallo beta-lactamase NDM-1, bleomycin resistance gene bleMBL, tellurite resistance protein, mercury resistance operon, chloramphenicol acetyltransferase, oxidoreductases, plasmid partition, transfer, stabilization, replication and pilin protein apparatus. IncFIB(pNDM-Mar) and IncHI1B(pNDM-MAR) plasmids present in our strain have also previously been reported in *K. pneumoniae* [31,32].

We validated our findings by doing a BLAST search (results retrieved on 7 December 2022) for plasmid sequences, and contig 5 and 6 were validated as plasmid sequences (Supplementary Fig. 1A and 1B). Top match for contig 5 was *K. pneumoniae* strain FDAARGOS_440 plasmid (with copies of erythromycin esterase, *MobC*, relaxase apparatus and beta-lactamase *OXA-232* gene), while for contig 6 was the strain *K. pneumoniae* Kpn-14 plasmid (coding for *TonB* receptor, type II toxin-antitoxin system *RelE*/*ParE* family toxin, DNA-binding beta-propeller fold protein *YncE* and plasmid stabilization protein). Hypothetical protein sequences were also present. Plannotate annotation (Supplementary Fig. 2A and 2B) showed that the contig 5 resembling *K. pneumoniae* strain FDAARGOS_440 plasmid consisted of *mbeA*, *mbeC*, and *ereA* gene with 59, 55 and 87% sequence identity, respectively. *mbeA* codes for relaxase that has a role in plasmid transfer through conjugation. *mbeC* helps *mbeA* through DNA bending while *ereA* confers resistance to the antibiotic erythromycin. Contig 6 resembling *K. pneumoniae* strain Kpn-14 plasmid consisted of the *CloDF13* (CDF) origin of replication (98% sequence identity), *rop* gene responsible for regulating the plasmid DNA replication through initiation of primer RNA precursor transcription and its modulation (61% sequence identity) and *mob* gene was detected that has a role in the plasmid transfer (62% sequence identity). Negligible identity was seen for the element *cos* which permits packaging into λ phage particles and *Csy4* CRISPR-associated endoribonuclease responsible for pre-crRNA processing. These were considered false-positives as the tool can mine false-positives at low sequence identity.

A total of 13 prophage regions were identified, of which 7 were intact (Fig. 1), 4 were incomplete, and 2 regions were questionable (Supplementary Table 1; Supplementary Fig. 3). Three CRISPR and 22 spacer sequences were also determined (Supplementary Table 2).

**Fig. 1.**
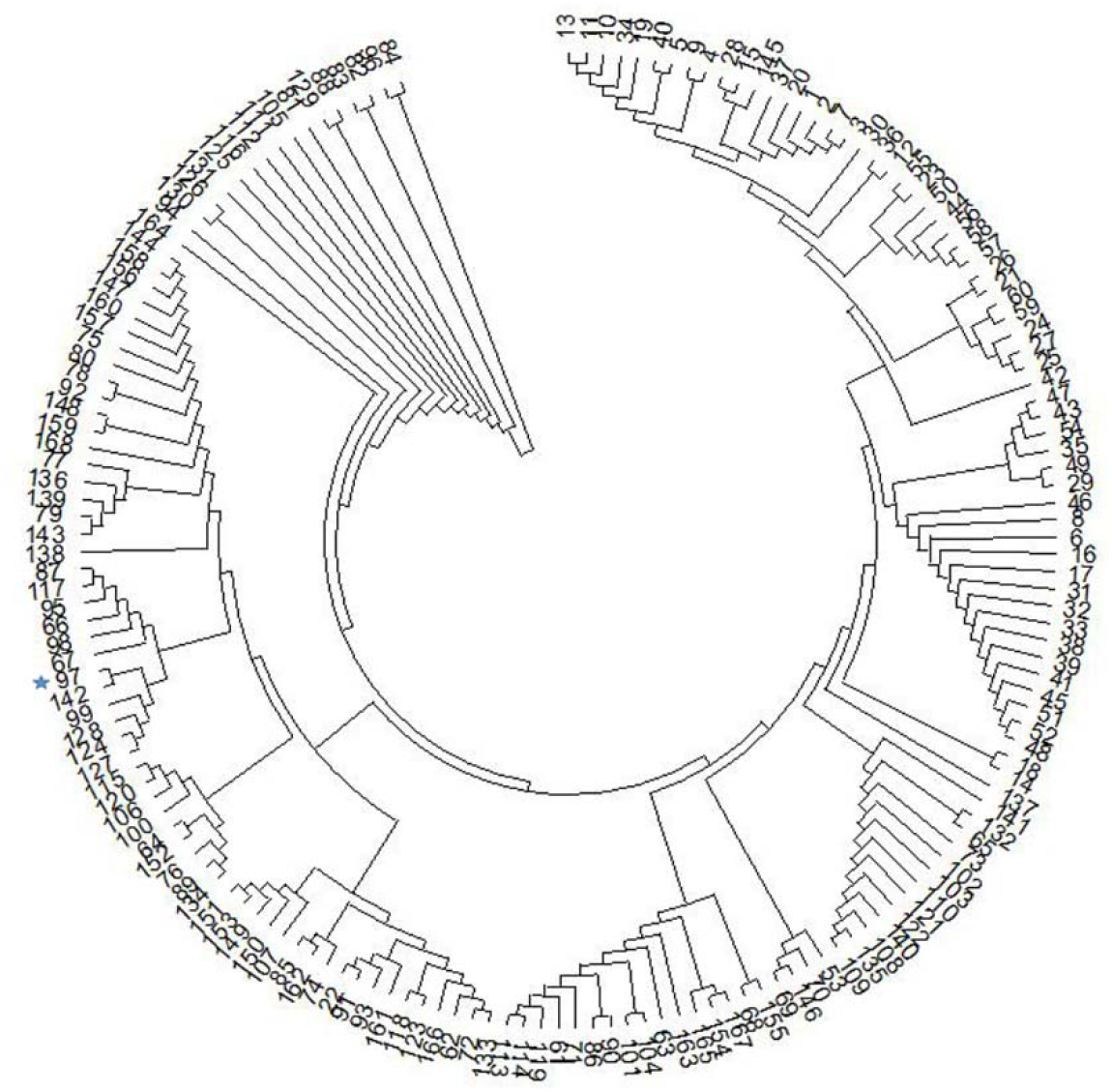
Phylogenetic tree from core genome and the position of *K. pneumoniae* JRCGR1 is marked by blue star (genome serial no. 97).

### 3.3. Virulence and antimicrobial resistance mapping

According to the RAST subsystem, 96 proteins involved in virulence, disease, or defense were identified. When JRCGR1 was subjected to virulence analysis using VFDB, 197 virulence factors were identified (Supplementary Table 3). Key virulence factors include outer membrane proteins, porins, transporter EfpA, fimbrial biogenesis proteins, fimbriae anchoring protein, multidrug efflux pump, multidrug resistance proteins etc. Apart from these, Type 6 Secretion System (T6SS) component TssC (ImpC/VipB), toxin delivery protein VgrG, nitric oxide regulator, carbon storage regulator, iron receptor, and siderophore producing proteins were also mined as virulence factors. T6SS is an anti-eukaryotic or anti-bacterial apparatus, with cell contact and puncture mode of action. It consists of cell effector and immunity proteins, where the effectors are toxic to the other cells while immunity proteins sheath from self-injury [33]. This system is regulated by PhoPQ two-component system and through ROS generation, the VgrG protein domain (DUF2345) kills bacteria and yeast in *K. pneumoniae* [34]. Since expression of T6SS is regulated by many factors like temperature, pH, iron and salt, so presence of regulator proteins also play a role in virulence [34].

CARD identified 25 antibiotic resistance genes (Table 4). Mechanisms mostly involved efflux, target alteration and antibiotic inactivation. For some virulence and antibiotic resistance genes, more than one copy of the gene was found.

**Table 4.**
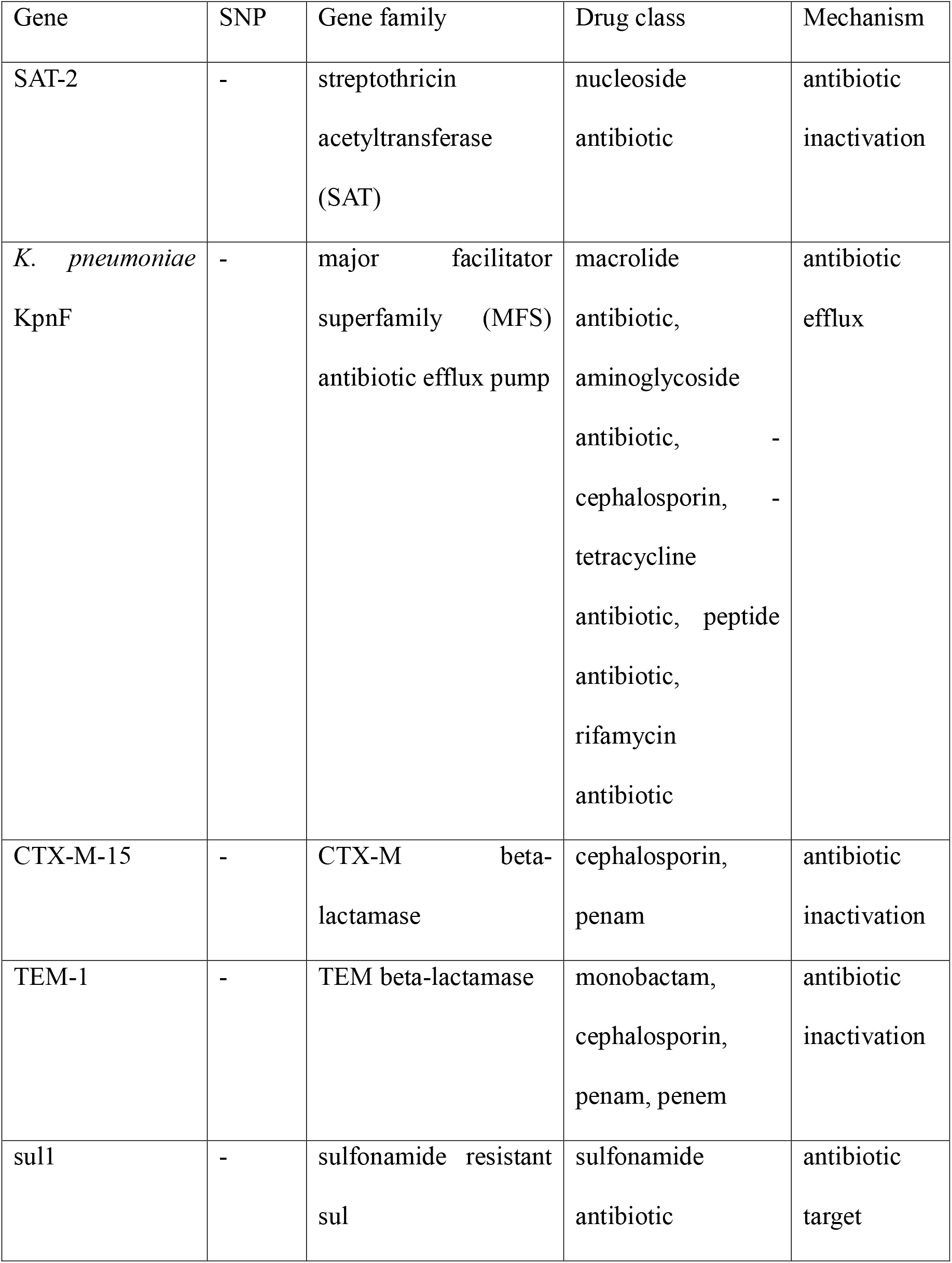
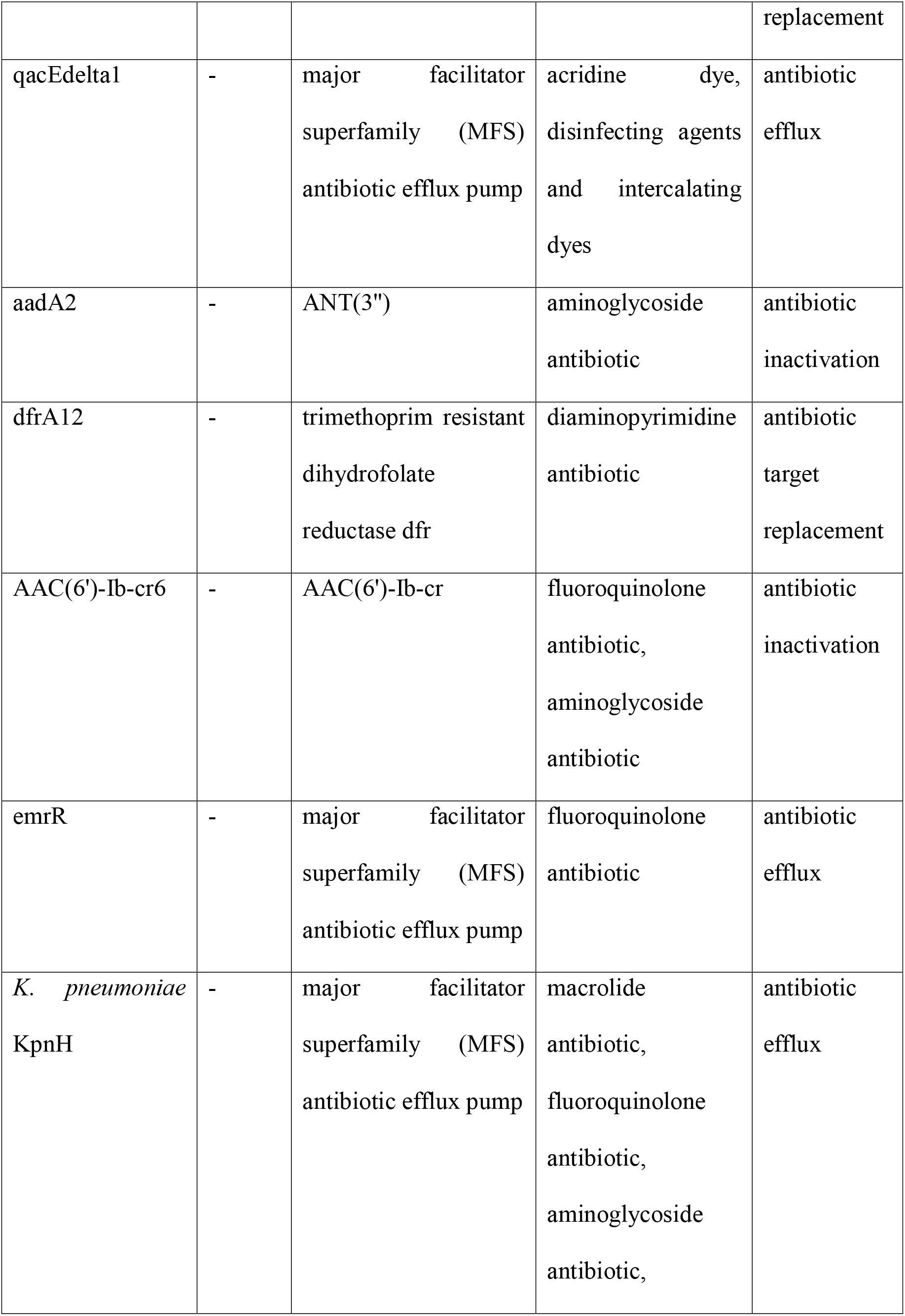
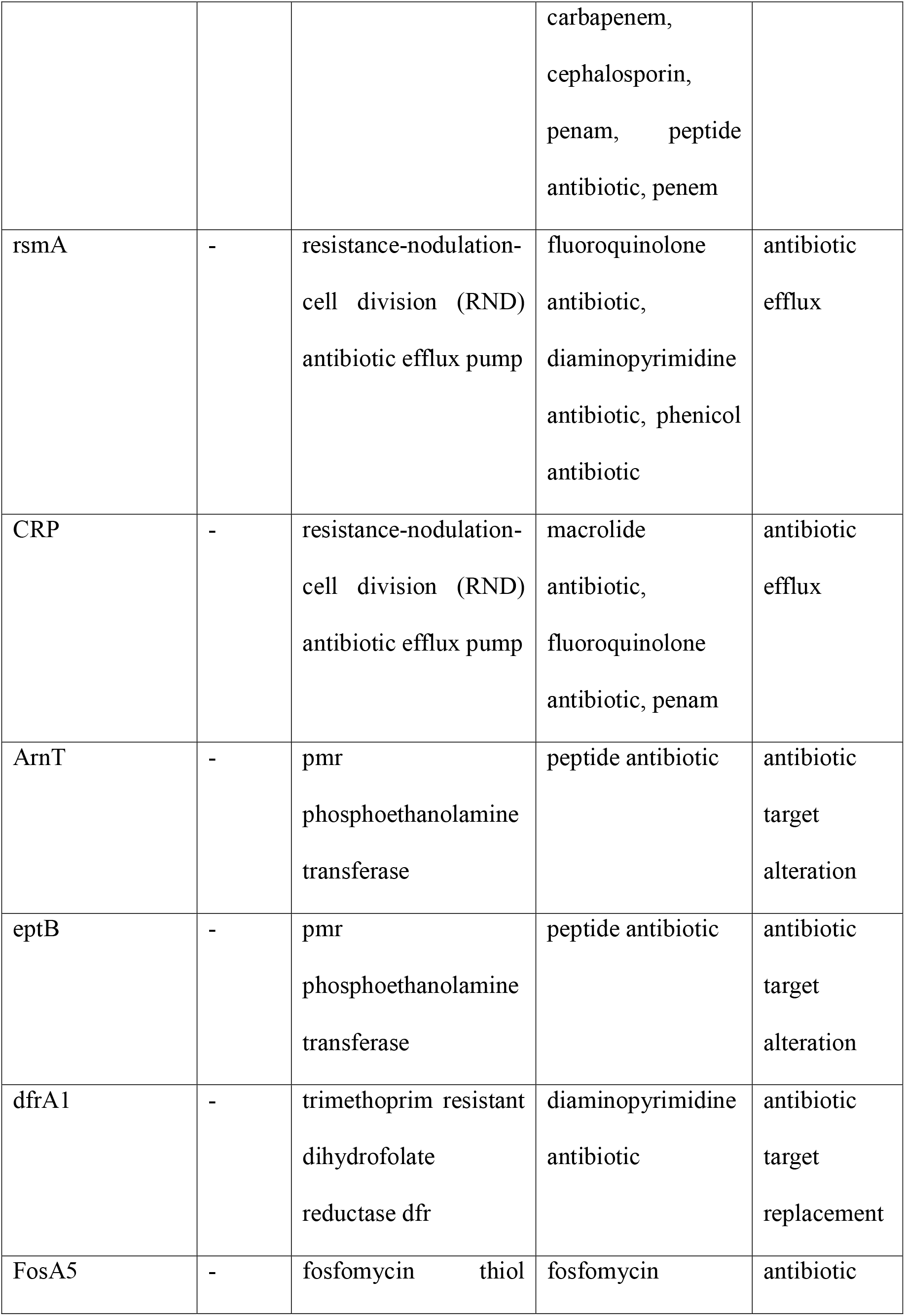
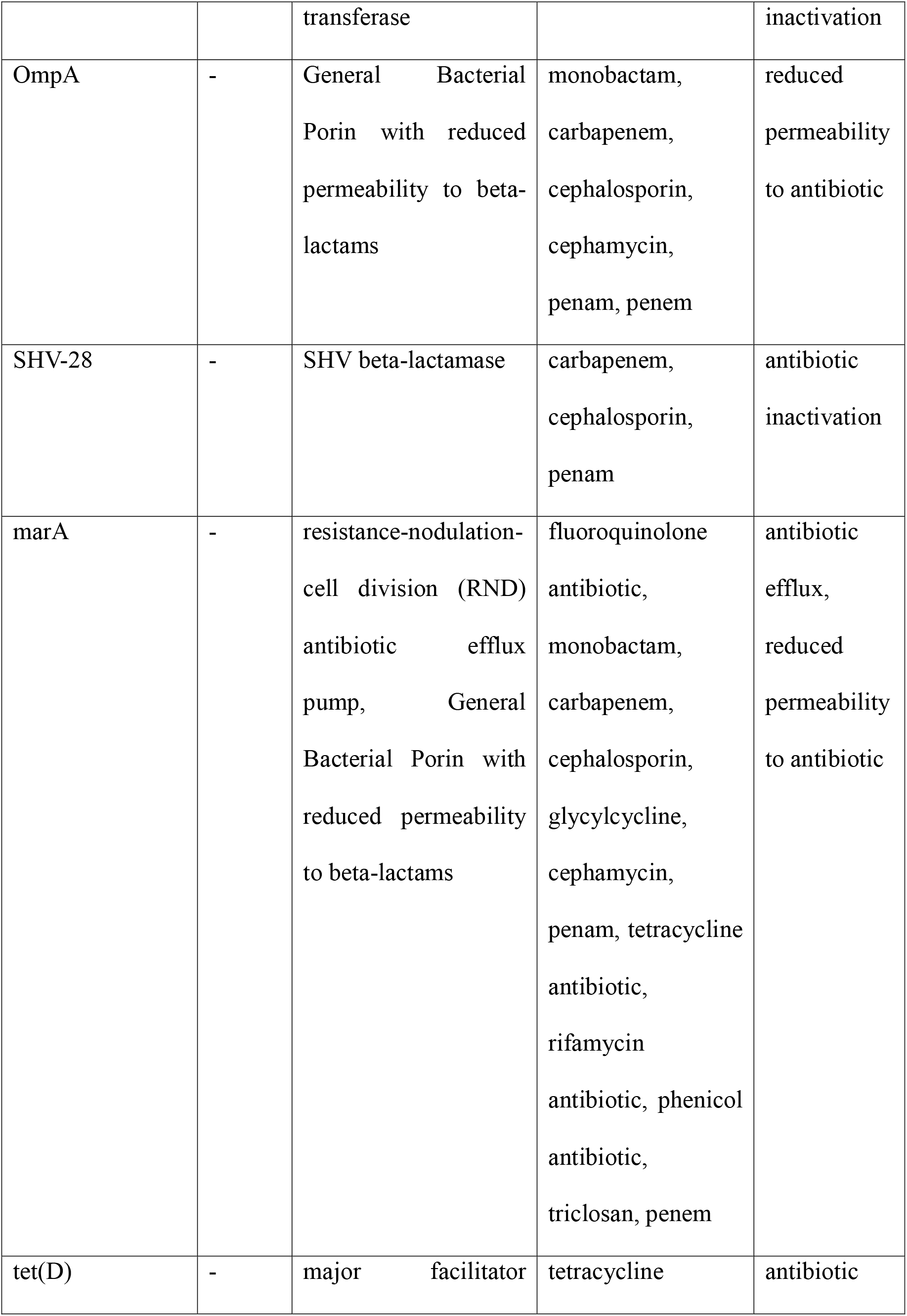
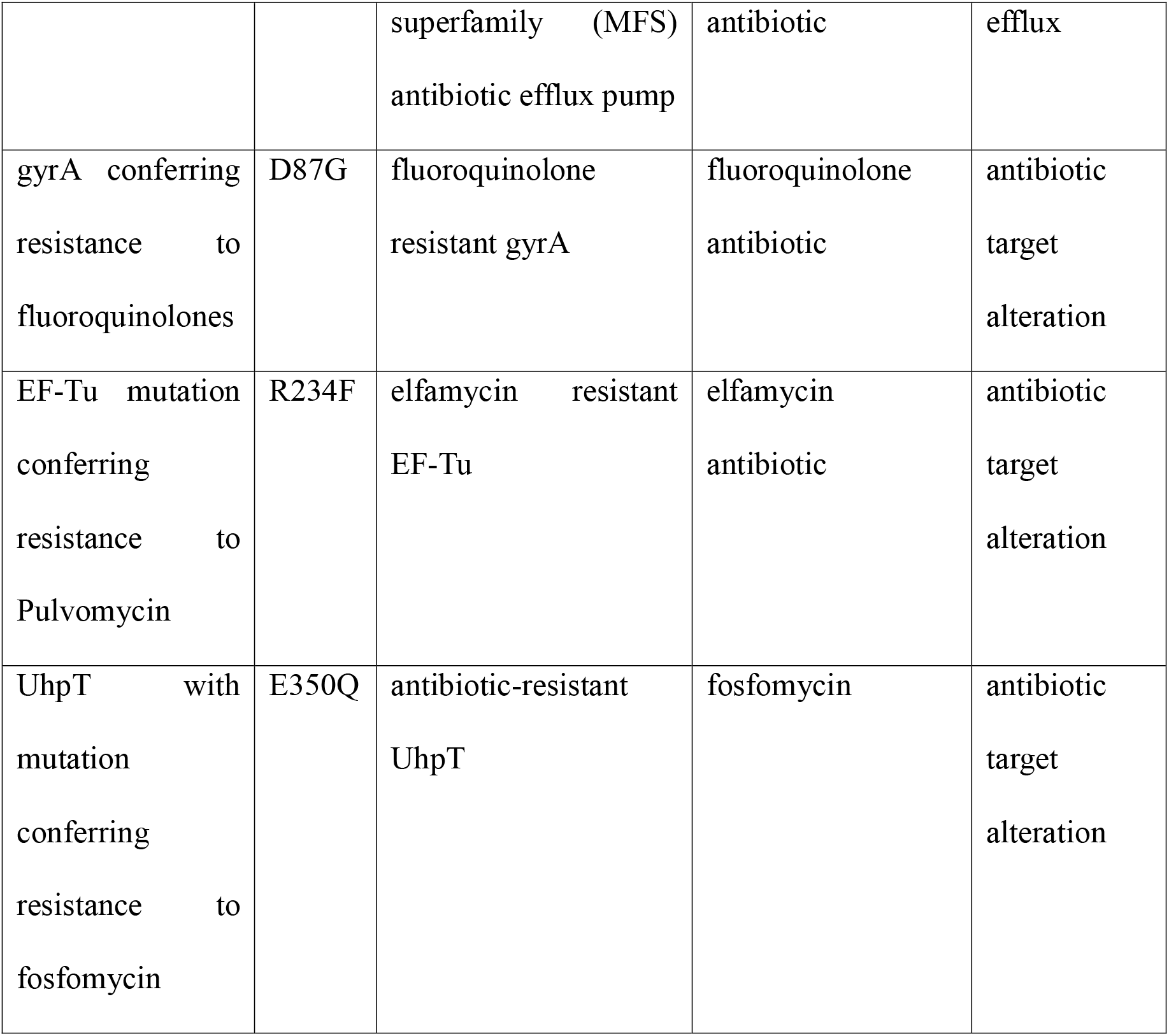
Antibiotic resistance genes identified in *K. pneumoniae* JRCGR1.

### 3.4. Pan-genomics

A list of Pakistani isolates used for pan-genome analysis is shown in Supplementary Table 4, along with statistics of CDSs, which formed the respective core and accessory genome of antibiotic-resistant strains (n=168) of this species, including *K. pneumoniae* JRCGR1. Core genome consisted of 1,303 genes. Core gene based phylogeny was inferred and a tree representation was made (Fig. 1), which showed several clades and sub-clades. Sum of branch length was 0.13, compared to 2.30 in pan-genome tree. JRCGR1 showed least distance to MMGX21 on core tree compared to colistin-resistant 10718 (isolated from soft tissue infection patient) and MMGA109 (isolated from tracheal secretion of repirtaory tract infection) on pan-genome tree. MMGX21 strain had been isolated from an indwelling device of a patient with post surgery infection and was NDM producing carbapenem resistant.

Pan-genome was inferred as open (b=0.42). Open pan-genomes usually exist in bacteria since horizontal gene transfer causes the addition of new genes at all times. JRCGR1 had 4,390 accessory genes, while 1,095 genes were unique and 31 genes were absent. Strain K184 had the highest number of unique genes i.e. 1,607, followed by JRCGR1. Strain 3189STDY5864759, CFSAN059643, KP_093, KP_116, and KP_136 did not carry any unique gene. PH10 had the largest number of missing genes i.e. 1,841 and only 638 genes were shared with the accessory genome. The highest number of accessory genes (n=4,731) were present in strain 3189STDY5864874. It also showed a large number of unique genes (n=184).

Total of 27 perfect and 38 strict hits were obtained for accessory fraction antibiotic resistance gene mapping. SHV-28, rsmA, aadA2, AAC(6’)-Ib-cr6, eptB and gyrA were unique to JRCGR1 compared to the studied genomes, while CRP, eptB, EF-Tu and UhpT were shared with core genome. However CTX-M-15 and SHV-28 beta-lactamases, present in JRCGR1 (which render resistance to cephalosporins and work by antibiotic inactivation), have already been reported in some areas of Pakistan via PCR tests [35,36] or genome [37] not listed in the PATRIC database (from where we picked our data). S83I mutation in gyrA gene conferring resistance to ciprofloxacin has been previously detected in Pakistani isolates [38], which is common to global lineage. gyrA works by antibiotic target alteration and also impacts aminoglycoside antibiotics. In our isolate, D87G mutation was observed, which confers resistance to fluoroquinolone and is near to the Texas lineage [39]. Nahid et al. [40] have reported the presence of aadA2 and AAC(6’)-Ib-cr in ST147 *K. pneumoniae* isolate. Variant AAC(6’)-Ib-cr6 was found in our isolate. These two genes are usually present on plasmids and work by antibiotic inactivation. Apart from mediating resistance to aminoglocosides, AAC(6’)-Ib-cr also confers resistance to fluoroquinolone. CRP is a resistance-nodulation-cell division (RND) antibiotic efflux pump and mediates resistance to macrolide antibiotics, fluoroquinolones and penams. It has not been well characterized in Pakistani *K. pneumoniae* isolates via PCR or other laboratory methods but its presence through *in silico* characterization indicates the resistance capability of *Klebsiella* sp. from Pakistan.

Ef-Tu (resistance to elfamycin with same mutation R234F) and UhpT (resistance to fosfomycin with mutation E350Q) were present in all isolates and have been reported from neighboring country Iran [41] but no PCR studies on this gene for *Klebsiella* sp. have been reported from Pakistan. An XDR *K. pneumonia* ST147 from India has also shown the presence of both these genes [42]. The presence of colistin resistance-conferring eptB gene in the core genome is of grave concern as colistin is the last resort cephalosporin antibiotic and recently, its presence has been reported through PCR-assay in *Klebsiella* sp. from Pakistan [43]. Urooj et al. [43] chose strains that already carried *mcr* genes. Nucleotide mutation G507A was present in both whole genome sequenced isolates but additionally, one carried C747T and the other A642T, C651T and C933T mutation. However, no amino acid mutation was inferred. Our strain lacked *mcr* gene but it has been noted that even with strains lacking this gene, the presence of eptB, as well as several lipid-A genes, lead to colistin resistance when mutated in gram-negative pathogens [44–46]. MarA gene fragment, causing resistance to colistin was identified based on sequence similarity.

To the best of our knowledge, this is the first national-scale pan-genome and pan-resistome report of *K. pneumoniae* from Pakistan. Although multidrug-resistant strains continue to pose a threat to the health of patients and the community, *K. pneumoniae* is not extensively studied. Constant monitoring and studies on antimicrobial resistance of *K. pneumoniae* need to be undertaken. Here, we have only presented the draft genome and associated plasmids but our group is further working on the epigenetics of this strain as well as comparison with global lineages.

## 4. Conclusion

*K. pneumoniae* JRCGR1 was isolated from an immunocompromised patient and shows that it has evolved to evade the pressure of antibiotics by acquiring mutations or by horizontal gene transfer. It can, therefore, cause hospital-acquired pneumonia outbreaks in immunocompromised patients. In addition, the antibiotic resistance profile shows that it can also affect the healthy community. It seems to have a T6SS for damaging neighbouring bacteria, yeast, or eukaryotes. Additionally, important regulators for utilization of iron production, carbon, nitrogen and nutrients seemingly can impact its virulence and need to be studied further. Apart from beta-lactamases and other AMR genes of concern, colistin resistance gene eptB was determined in the core fraction of the genome. The pan-genome was inferred to be open and further information from downstream analysis might facilitate the development of drugs and vaccines against this species. The information from this genome may as well as be useful to researchers and clinicians with an interest in antibiotic resistance profiling, monitoring of pathogens, and screening drugs against resistant isolates of *K. pneumoniae*.

## Data availability

The genome has been submitted to NCBI and given the accession no. JAMKOI000000000.

## Ethical approval

Ethical approval was obtained from Ethical committee of the International Center for Chemical and Biological Sciences, Dr. Panjwani Center for Molecular Medicine and & Drug Research, University of Karachi, Ref. ICCBS/IEC-070-HUS-2021/Protocol/1.0.

## Acknowledgement

The project was funded by ICCBS, UoK (project no: 3702-2020).

## Author contribution

ZB conceived and designed the experiment. MI, ZN, MA, SSH, RMAK and ZB performed experiments. ZB and RMAK provided resources. ZB and AK analyzed data. ZB validated the findings. ZB wrote the manuscript. All authors read and approved the paper.

## References

1. Bengoechea, J.A.; Sa Pessoa, J. *Klebsiella pneumoniae* infection biology: living to counteract host defences. FEMS Microbiol Rev 2019, 43, 123–144, doi: 10.1093/femsre/fuy043.

2. Paczosa, M.K.; Mecsas, J. *Klebsiella pneumoniae:* Going on the Offense with a Strong Defense. Microbiol Mol Biol Rev 2016, 80, 629–661, doi:10.1128/MMBR.00078-15.

3. Yu, W.L.; Chuang, Y.C. Clinical features, diagnosis, and treatment of *Klebsiella pneumoniae* infection. CalderWood 2015.

4. Liu, C.; Du, P.; Xiao, N.; Ji, F.; Russo, T.A.; Guo, J. Hypervirulent *Klebsiella pneumoniae* is emerging as an increasingly prevalent *K. pneumoniae* pathotype responsible for nosocomial and healthcare-associated infections in Beijing, China. Virulence 2020, 11, 1215–1224, doi:10.1080/21505594.2020.1809322.

5. Ali, S.; Alam, M.; Hasan, G.M.; Hassan, M.I. Potential therapeutic targets of *Klebsiella pneumoniae:* a multi-omics review perspective. Brief Funct Genom 2021, doi:10.1093/bfgp/elab038.

6. Al-Nakeeb, N.K.; Radi, J.; Hamdan, K.; Fouad, Z. Clinical and immunological effects of experimental infection with *Klebsiella pneumoniae* in lambs in Iraq. Al-Qadisiyah J Vet Med Sci, 2018, 17, 44–48.

7. Chang, E.K.; Miller, M.; Shahin, K.; Batac, F.; Field, C.L.; Duignan, P.; Struve, C.; Byrne, B.A.; Murray, M.J.; Greenwald, K.; et al. Genetics and pathology associated with *Klebsiella pneumoniae* and *Klebsiella* spp. isolates from North American Pacific coastal marine mammals.. Vet Microbiol 2022, 265, 109307.

8. Bassetti, M.; Righi, E.; Carnelutti, A.; Graziano, E.; Russo, A. Multidrug-resistant *Klebsiella pneumoniae*: challenges for treatment, prevention and infection control. Expert Rev Anti-infective Therap 2018, 16, 749–761.

9. Al-Zalabani, A.; AlThobyane, O.A.; Alshehri, A.H.; Alrehaili, A.O.; Namankani, M.O.; Aljafri, O.H. Prevalence of *Klebsiella pneumoniae* antibiotic resistance in Medina, Saudi Arabia, 2014-2018. Cureus 2020, 12.

10. Musawy, W.K.A.; Hashimy, A.B. Molecular study and antibiotic susceptibility patterns of some extended spectrum beta-lactamase genes (ESBL) Of *Klebsiella Pneumoniae* from urinary tract infections. European J Mol Clin Med, 2020, 7, 796–804.

11. Akinbami, O.R.; Olofinsae, S.; Ayeni, F.A. Prevalence of extended spectrum beta lactamase and plasmid mediated quinolone resistant genes in strains of *Klebsiella pneumonia*, *Morganella morganii*, *Leclercia adecarboxylata* and *Citrobacter freundii* isolated from poultry in South Western Nigeria. PeerJ 2018, 6, e5053, doi:10.7717/peerj.5053.

12. Devipalanisamy, D.; Olaganathan, S.; Marimuthu, M. Detection of extended spectrum beta lactamases (ESBLs) producing Enterobacteriaceae family from urinary tract infection (UTI) patients. Int J Pharm Invest 2021, 11, 113–117.

13. King, T.L.; Schmidt, S.; Essack, S.Y. Antibiotic resistant *Klebsiella* spp. from a hospital, hospital effluents and wastewater treatment plants in the uMgungundlovu District, KwaZulu-Natal, South Africa. Sci Total Environ 2020, 712, 135550.

14. Choi, M.J.; Ko, K.S. Loss of hypermucoviscosity and increased fitness cost in colistin-resistant *Klebsiella pneumoniae* sequence type 23 strains. Antimicrob Agents Chemother 2015, 59, 6763–6773, doi:10.1128/AAC.00952-15.

15. Ontong, J.C.; Ozioma, N.F.; Voravuthikunchai, S.P.; Chusri, S. Synergistic antibacterial effects of colistin in combination with aminoglycoside, carbapenems, cephalosporins, fluoroquinolones, tetracyclines, fosfomycin, and piperacillin on multidrug resistant *Klebsiella pneumoniae* isolates. PLoS One 2021, 16, e0244673, doi:10.1371/journal.pone.0244673.

16. Heinz, E.; Ejaz, H.; Bartholdson Scott, J.; Wang, N.; Gujaran, S.; Pickard, D.; Wilksch, J.; Cao, H.; Haq, I.U.; Dougan, G.; et al. Resistance mechanisms and population structure of highly drug resistant *Klebsiella* in Pakistan during the introduction of the carbapenemase NDM-1. Sci Rep 2019, 9, 2392, doi:10.1038/s41598-019-38943-7.

17. Khan, A.U.; Maryam, L.; Zarrilli, R. Structure, genetics and worldwide spread of New Delhi Metallo-beta-lactamase (NDM): a threat to public health. BMC Microbiol 2017, 17, 101, doi:10.1186/s12866-017-1012-8.

18. Lewis II, J.S. Performance Standards for Antimicrobial Susceptibility Testing, 32nd ed.; 2022.

19. Wick, R.R.; Judd, L.M.; Gorrie, C.L.; Holt, K.E. Unicycler: Resolving bacterial genome assemblies from short and long sequencing reads. PLoS Comput Biol 2017, 13, e1005595, doi:10.1371/journal.pcbi.1005595.

20. Brettin, T.; Davis, J.J.; Disz, T.; Edwards, R.A.; Gerdes, S.; Olsen, G.J.; Olson, R.; Overbeek, R.; Parrello, B.; Pusch, G.D.; et al. RASTtk: a modular and extensible implementation of the RAST algorithm for building custom annotation pipelines and annotating batches of genomes. Sci Rep 2015, 5, 8365, doi:10.1038/srep08365.

21. Chaudhari, N.M.; Gupta, V.K.; Dutta, C. BPGA-an ultra-fast pan-genome analysis pipeline. Sci Rep 2016, 6, 24373, doi:10.1038/srep24373.

22. Alcock, B.P.; Raphenya, A.R.; Lau, T.T.Y.; Tsang, K.K.; Bouchard, M.; Edalatmand, A.; Huynh, W.; Nguyen, A.V.; Cheng, A.A.; Liu, S.; et al. CARD 2020: antibiotic resistome surveillance with the comprehensive antibiotic resistance database. Nucleic Acids Res 2020, 48, D517–D525, doi:10.1093/nar/gkz935.

23. Bailey, D.C.; Alexander, E.; Rice, M.R.; Drake, E.J.; Mydy, L.S.; Aldrich, C.C.; Gulick, A.M. Structural and functional delineation of aerobactin biosynthesis in hypervirulent Klebsiella pneumoniae. J Biol Chem 2018, 293, 7841–7852, doi: 10.1074/jbc.RA118.002798.

24. Batista, F.M.; Stapleton, T.; Lowther, J.A.; Fonseca, V.G.; Shaw, R.; Pond, C.; Walker, D.I.; van Aerle, R.; Martinez-Urtaza, J. Whole genome sequencing of Hepatitis A virus using a PCR-free single-molecule nanopore sequencing approach. Front Microbiol 2020, 11, 874, doi:10.3389/fmicb.2020.00874.

25. Schmidt, J.; Blessing, F.; Fimpler, L.; Wenzel, F. Nanopore sequencing in a clinical routine laboratory: Challenges and opportunities. Clin Lab 2020, 66, doi:10.7754/Clin.Lab.2019.191114.

26. Lu, H.; Giordano, F.; Ning, Z. Oxford nanopore MinION sequencing and genome assembly. Genom Proteom Bioinform 2016, 14, 265–279, doi:10.1016/j.gpb.2016.05.004.

27. van Dijk, E.L.; Jaszczyszyn, Y.; Naquin, D.; Thermes, C. The Third revolution in sequencing technology. Trends Genet 2018, 34, 666–681, doi:10.1016/j.tig.2018.05.008.

28. Azam, M.; Gaind, R.; Yadav, G.; Sharma, A.; Upmanyu, K.; Jain, M.; Singh, R. Colistin resistance among multiple sequence types of *Klebsiella pneumoniae* is associated with diverse resistance mechanisms: A report from India. Front Microbiol 2021, 12, 609840, doi:10.3389/fmicb.2021.609840.

29. Russo, T.A.; Olson, R.; Macdonald, U.; Metzger, D.; Maltese, L.M.; Drake, E.J.; Gulick, A.M. Aerobactin mediates virulence and accounts for increased siderophore production under iron-limiting conditions by hypervirulent (hypermucoviscous) Klebsiella pneumoniae. Infect Immun 2014, 82, 2356–2367, doi:10.1128/IAI.01667-13.

30. Boonyasiri, A.; Jauneikaite, E.; Brinkac, L.M.; Greco, C.; Lerdlamyong, K.; Tangkoskul, T.; Nguyen, K.; Thamlikitkul, V.; Fouts, D.E. Genomic and clinical characterisation of multidrug-resistant carbapenemase-producing ST231 and ST16 *Klebsiella pneumoniae* isolates colonising patients at Siriraj hospital, Bangkok, Thailand from 2015 to 2017. BMC Infect Dis 2021, 21, 142, doi:10.1186/s12879-021-05790-9.

31. Musicha, P.; Msefula, C.L.; Mather, A.E.; Chaguza, C.; Cain, A.K.; Peno, C.; Kallonen, T.; Khonga, M.; Denis, B.; Gray, K.J.; et al. Genomic analysis of *Klebsiella pneumoniae* isolates from Malawi reveals acquisition of multiple ESBL determinants across diverse lineages. J Antimicrob Chemother 2019, 74, 1223–1232, doi:10.1093/jac/dkz032.

32. Saavedra, S.Y.; Bernal, J.F.; Montilla-Escudero, E.; Arevalo, S.A.; Prada, D.A.; Valencia, M.F.; Moreno, J.; Hidalgo, A.M.; Garcia-Vega, A.S.; Abrudan, M.; et al. Complexity of genomic epidemiology of carbapenem-resistant *Klebsiella pneumoniae* isolates in colombia urges the reinforcement of whole genome sequencing-based surveillance programs. Clin Infect Dis 2021, 73, S290–S299, doi:10.1093/cid/ciab777.

33. Liu, L.; Ye, M.; Li, X.; Li, J.; Deng, Z.; Yao, Y.F.; Ou, H.Y. Identification and characterization of an antibacterial type vi secretion system in the carbapenem-resistant strain *Klebsiella pneumoniae* HS11286. Front Cell Infect Microbiol 2017, 7, 442, doi:10.3389/fcimb.2017.00442.

34. Storey, D.; McNally, A.; Astrand, M.; Sa-Pessoa Graca Santos, J.; Rodriguez-Escudero, I.; Elmore, B.; Palacios, L.; Marshall, H.; Hobley, L.; Molina, M.; et al. *Klebsiella pneumoniae* type VI secretion system-mediated microbial competition is PhoPQ controlled and reactive oxygen species dependent. PLoS Pathog 2020, 16, e1007969, doi:10.1371/journal.ppat.1007969.

35. Hadjadj, L.; Syed, M.A.; Abbasi, S.A.; Rolain, J.M.; Jamil, B. Diversity of carbapenem resistance mechanisms in clinical gram-negative bacteria in Pakistan. Microb Drug Resist 2021, 27, 760–767, doi:10.1089/mdr.2019.0387.

36. Muzaheed; Sattar Shaikh, N.; Sattar Shaikh, S.; Acharya, S.; Sarwar Moosa, S.; Habeeb Shaikh, M.; F, M.A.; Ibrahim Alomar, A. Molecular epidemiological surveillance of ctx-m-15-producing *Klebsiella pneumoniae* from the patients of a teaching hospital in Sindh, Pakistan. F1000Res 2021, 10, 444, doi:10.12688/f1000research.53221.3.

37. Rahmat Ullah, S.; Majid, M.; Andleeb, S. Draft genome sequence of an extensively drug-resistant neonatal *Klebsiella pneumoniae* isolate harbouring multiple plasmids contributing to antibiotic resistance. J Glob Antimicrob Resist 2020, 23, 100–101, doi:10.1016/j.jgar.2020.08.008.

38. Alvi, R.F.; Aslam, B.; Shahzad, N.; Rasool, M.H.; Shafique, M. Molecular basis of quinolone resistance in clinical isolates of *Klebsiella pneumoniae* from Pakistan. Pak J Pharm Sci 2018, 31, 1591–1596.

39. Peirano, G.; Chen, L.; Kreiswirth, B.N.; Pitout, J.D.D. Emerging antimicrobial-resistant high-risk *Klebsiella pneumoniae* clones ST307 and ST147. Antimicrob Agents Chemother 2020, 64, doi:10.1128/AAC.01148-20.

40. Nahid, F.; Zahra, R.; Sandegren, L. A blaOXA-181-harbouring multi-resistant ST147 *Klebsiella pneumoniae* isolate from Pakistan that represent an intermediate stage towards pan-drug resistance. PLoS One 2017, 12, e0189438, doi:10.1371/journal.pone.0189438.

41. Bolourchi, N.; Shahcheraghi, F.; Giske, C.G.; Nematzadeh, S.; Noori Goodarzi, N.; Solgi, H.; Badmasti, F. Comparative genome analysis of colistin-resistant OXA-48-producing *Klebsiella pneumoniae* clinical strains isolated from two Iranian hospitals. Ann Clin Microbiol Antimicrob 2021, 20, 74, doi:10.1186/s12941-021-00479-y.

42. Dey, S.; Gaur, M.; Sahoo, R.K.; Das, A.; Jain, B.; Pati, S.; Subudhi, E. Genomic characterization of XDR *Klebsiella pneumoniae* ST147 co-resistant to carbapenem and colistin - The first report in India. J Glob Antimicrob Resist 2020, 22, 54–56, doi:10.1016/j.jgar.2020.05.005.

43. Urooj, M.; Ullah, R.; Ali, S.; Mohyuddin, A.; Mirza, H.M.; Faryal, R. Elucidation of molecular mechanism for colistin resistance among gram-negative isolates from tertiary care hospitals. J Infect Chemother 2022, 28, 602–609, doi:10.1016/j.jiac.2022.01.002.

44. Mathur, P.; Veeraraghavan, B.; Devanga Ragupathi, N.K.; Inbanathan, F.Y.; Khurana, S.; Bhardwaj, N.; Kumar, S.; Sagar, S.; Gupta, A. Multiple mutations in lipid-A modification pathway & novel fosA variants in colistin-resistant *Klebsiella pneumoniae*. Future Sci OA 2018, 4, FSO319, doi:10.4155/fsoa-2018-0011.

45. Pragasam, A.K.; Shankar, C.; Veeraraghavan, B.; Biswas, I.; Nabarro, L.E.; Inbanathan, F.Y.; George, B.; Verghese, S. Molecular mechanisms of colistin resistance in *Klebsiella pneumoniae* causing bacteremia from India-a first report. Front Microbiol 2016, 7, 2135, doi:10.3389/fmicb.2016.02135.

46. Zhang, H.; Srinivas, S.; Xu, Y.; Wei, W.; Feng, Y. Genetic and biochemical mechanisms for bacterial lipid a modifiers associated with polymyxin resistance. Trends Biochem Sci 2019, 44, 973–988, doi:10.1016/j.tibs.2019.06.002.

