## Supplementary Data for "Nanopore sequencing of antibiotic-resistant *Klebsiella pneumoniae* JRCGR1 isolate from Pakistan and country scale pan-genomics"

Supplementary Table 1. Prophage regions in *K. pneumoniae* JRCGR1.

| **Region** | **Region Length** | **Completeness** | **Score** | **# Total Proteins** | **Region Position** | **Most Common Phage** | **GC %** |
| --- | --- | --- | --- | --- | --- | --- | --- |
| contig_1,length,5410184 |  |  |  |  |  |  |  |
| 1 | 60.4Kb | intact | 150 | 94 | [364103-424568](https://phaster.ca/submissions/ZZ_5d7ac7b91d#region_dna0) | PHAGE_Edward_GF_2_NC_026611(16) | 53.95% |
| 2 | 46.8Kb | intact | 150 | 68 | [524214-571020](https://phaster.ca/submissions/ZZ_5d7ac7b91d#region_dna1) | PHAGE_Erwini_vB_EhrS_59_NC_048198(14) | 52.84% |
| 3 | 51.5Kb | intact | 150 | 93 | [1260946-1312520](https://phaster.ca/submissions/ZZ_5d7ac7b91d#region_dna2) | PHAGE_Escher_500465_1_NC_049342(11) | 51.48% |
| 4 | 3.9Kb | questionable | 70 | 6 | [2225540-2229496](https://phaster.ca/submissions/ZZ_5d7ac7b91d#region_dna3) | PHAGE_Stx2_c_Stx2a_F451_NC_049924(3) | 50.80% |
| 5 | 10.6Kb | incomplete | 50 | 17 | [2529978-2540626](https://phaster.ca/submissions/ZZ_5d7ac7b91d#region_dna4) | PHAGE_Bacill_G_NC_023719(2) | 49.98% |
| 6 | 13.9Kb | incomplete | 60 | 23 | [4291184-4305139](https://phaster.ca/submissions/ZZ_5d7ac7b91d#region_dna5) | PHAGE_Cronob_vB_CsaM_GAP32_NC_019401(2) | 58.23% |
| 7 | 57.9Kb | intact | 114 | 57 | [4594186-4652178](https://phaster.ca/submissions/ZZ_5d7ac7b91d#region_dna6) | PHAGE_Klebsi_ST16_OXA48phi5.4_NC_049450(35) | 53.84% |
| contig_2,length,400087 |  |  |  |  |  |  |  |
| 8 | 6.3Kb | incomplete | 40 | 8 | [20406-26727](https://phaster.ca/submissions/ZZ_5d7ac7b91d#region_dna7) | PHAGE_Escher_RCS47_NC_042128(2) | 47.28% |
| 9 | 16.2Kb | incomplete | 20 | 33 | [79251-95451](https://phaster.ca/submissions/ZZ_5d7ac7b91d#region_dna8) | PHAGE_Klebsi_Matisse_NC_028750(2) | 44.53% |
| 10 | 26.2Kb | intact | 100 | 44 | [108126-134411](https://phaster.ca/submissions/ZZ_5d7ac7b91d#region_dna9) | PHAGE_Salmon_SJ46_NC_031129(4) | 47.81% |
| 11 | 5.5Kb | questionable | 90 | 10 | [218053-223620](https://phaster.ca/submissions/ZZ_5d7ac7b91d#region_dna10) | PHAGE_Stx2_c_Stx2a_F451_NC_049924(3) | 54.96% |
| 12 | 25.8Kb | intact | 100 | 36 | [333195-359023](https://phaster.ca/submissions/ZZ_5d7ac7b91d#region_dna11) | PHAGE_Stx2_c_Stx2a_F451_NC_049924(3) | 49.84% |
| [contig_3,length,40771](https://phaster.ca/submissions/ZZ_5d7ac7b91d#region_dna12) |  |  |  |  |  |  |  |
| 13 | 40.2Kb | intact | 150 | 78 | [3-40246](https://phaster.ca/submissions/ZZ_5d7ac7b91d#region_dna12) | PHAGE_Entero_mEp235_NC_019708(8) | 50.09% |

Supplementary Table 2. CRISPR/Cas elements in *K. pneumoniae* JRCGR1.

| **Element** | **CRISPR Id / Cas Type** | **Start** | **End** | **Spacer / Gene** | **Repeat consensus / cas genes** | **Direction** | **Evidence Level** |
| --- | --- | --- | --- | --- | --- | --- | --- |
| CRISPR | [contig_1_length_5410184](https://crisprcas.i2bc.paris-saclay.fr/CrisprCasFinder/Viewing/637888973493882004) | 147562 | 148327 | 12 | GGAAACACCCCCACGTGCGTGGGGAAGAC | ND | 4 |
| Cas cluster | [CAS-TypeIE](https://crisprcas.i2bc.paris-saclay.fr/CrisprCasFinder/Viewing/637888973493882004) | 148375 | 156648 | 6 | cas2_TypeIE, cas5_TypeIE, cas6_TypeIE, cse1_TypeIE, cas3_TypeI, cas3a_TypeI | | |
| CRISPR | contig_1_length_5410184 | 157060 | 157637 | 9 | AGAAACACCCCCACGCGTGTGGGGAAGAC | ND | 4 |
| CRISPR | [contig_1_length_5410184](https://crisprcas.i2bc.paris-saclay.fr/CrisprCasFinder/Viewing/637888973493882004) | 3571210 | 3571338 | 1 | GTTTTGTAGGCCGGGTAAGGCGCCAGCCGCCACCCGGCA | - | 1 |

### Supplementary Table 3. Virulence factors.

| Serial no. | Name | Sequence |
| --- | --- | --- |
| 1. | Protein_ID\|573.43724.peg.122_Carbonic anhydrase, beta class (EC 4.2.1.1) | MQHIIEGFLNFQKEIFPQRKELFRSLASSQNPKALFISCSDSRLVPELVTQQEPGQLFVI  RNAGNIVPSFGPEPGGVSATIEYAVVALGVTDIVICGHSNCGAMKAIATCQCLEPMPAVS  HWLRYADAAKAVVEKKTWASETDKVNGMVQENVIAQLNNIKTHPSVAVGLRDHTLRLHGW  FYDIETGDIQALDKNTKSFVSLSENPDVFFE |
| 2. | Protein_ID\|573.43724.peg.157_Transposase | MSDNLFNGRRFRALTVVDNFSRECVAIHAGKSLKGEDVVGVMESLRVLGKRLPVRIQTDN GSEFISKSLDKWAYEHGVTMDFSRPGKPTDNSFIESFNGSLRDECLNIHWFLSLEDAQDK LDNWRREYNHERTHSSLNDMTPAEFIRSLRKDEDL |
| 3. | Protein_ID\|573.43724.peg.227_Cytochrome c heme lyase subunit CcmF | MPLDAVARVLAVMGMIAFGFLLFILFTSNPFSRGLPQYPIDGRDLNPLLQDIGMIFHPPI  LYMGYVGFSVAFAFAIASLLAGRLDTAWARWSRPWTQAAWMFLTLGIVLGSAWAYYELGW  GGWWFWDPVENASFMPWLVGTALLHSLAVTEKRGSFRAWTVLLAIAAFSLCLLGTFLVRS  GVLVSVHAFASDPSRGLFILVLLIVAIGGSLLLYALKGGRVRARVEHTLWSRESFLLGNN  ILLMAAMLVVLLGTLLPLVHKELGLGSISIGEPFFNTMFTALMAPFSLLLGLGPLIRWRR  DDVARQIKRLIIALLVTLSLSLALPWLLQDRITAMAVIGLMMALWVLIFALMEVHERATH  RHGFWRGLRTLTRSQWGMVLGHVGVAVTVIGITFSQNYSVERDVRMRPGDSIDIHRYHFV  FNGVRNIVGPNWTGGEGIIAVTRNGRPEATLYAEKRFYTASRMMMTEAAISGGLTRDLYA  ALGEELSDGSWAVRLYYKPFVRWIWYGGVLMALGGLCCMLDPRYRMRKKLQEAS |
| 4. | Protein_ID\|573.43724.peg.229_Cytochrome c-type biogenesis protein CcmE, heme chaperone | MGGYVQPGSLQRDPQTLDVRFKLYDARGVVDVSYKGILPDLFREGQGVVAQGVLDGERHI TAQQVLAKHDENYTPPEVKNAMTPEKTGTQP |
| 5. | Protein_ID\|573.43724.peg.231_Cytochrome c-type biogenesis protein CcmC, putative heme lyase for CcmE | MVDGAVPAMAVAAFVGVVWQIKMADLAIAALAPVGAVCTLVALVSGAAWGKPMWGTWWIW DARLTSELVLLFLYAGVIALWHAFDDRRLAGRAAGILVLVGVVNLPIIHYSVYWWNTLHQ GSTNLQQTIDPSMRLPAADLHLCLPHAVGHPDPDAPA |
| 6. | Protein_ID\|573.43724.peg.374_Transcriptional regulator KPN_02146, AcrR family | MAHFSRYGYEKTTVTDLAKAIGFSKAYIYKFFDSKQAIGEAICASRLEKIMVAVSEAIAD APSASEKLRRLFRALTEAGRNCFSRIANCTTSPPSRRAINGLQRNSMPVICSS |
| 7. | Protein_ID\|573.43724.peg.402_2-keto-3-deoxy-D-arabino-heptulosonate-7-phosphate synthase I alpha (EC 2.5.1.54) | MPCSCNKQGLVYRDAHHFPAFSPRKKQIFETPMNKTDELRTARIESLVTPAELAQRHPVT ADVAAHVSASRRRIEKILNGEDRRLLVIIGPCSIHDTDAALEYARRLQGMRERYQPQLEI VMRTYFEKPRTVVGWKGLISDPDLNGSYRVNHGIELARRLLLQVNELGVPTATEFLDMVT GQFIADLISWGPSGRAPQKARSTVRWPQRSPARSVLKTAPTAIRASRWTPSAPPAPAICS SLRTSRGR |
| 8. | Protein_ID\|573.43724.peg.403_2-keto-3-deoxy-D-arabino-heptulosonate-7-phosphate synthase I alpha (EC 2.5.1.54) | MFLSPDKQGQMTIYQTSGNPYGHIIMRGGKRPNYHAEDIAAAGEALREFDLPEQLVVDFS HGNCQKQHRRQLEVCADICQQIRAGSTAIAGIMAESFLQEGTQKVVPGQPLTWGQSITDP CLSWEDSERLLSELAAATATRL |
| 9. | Protein_ID\|573.43724.peg.448_hypothetical protein | MFSISPVFFWANIYVPADFEDYCVNTLKKTALLSVLALYIPVSQAAAKEYSLDPQHTSVV ISWNHFGFSNPTAYISDVSGKLAFDKENPEKSSVNVTLPVKTIDAHVKALTDEFLGKEYF DVKTFPDATFQSTKVESKGDNKYDVEGNLTIKGITKPVVLHAVLNKQDMHPMVKKEAIGF DATGVIKRSDFKLDKYVPAVSDNVTITLSTEAYAK |
| 10. | Protein_ID\|573.43724.peg.483_Respiratory nitrate reductase alpha chain (EC 1.7.99.4) | MKPEEVEWRDNGLDGKLDLVVTLDFRLSSTCLYSDIVLPTATWYEKDDMNTSDMHPFIHP LSAAVDPAWESKSDWEIYKGIAKKFSEVCVGHLGKETDVVTLPIQHDSAAEMAQPLDVKDWKKGECDLIPGKTAPHIIPVERDYPATYERFTSIGPLLETIGNGGKGIAWNTQSEMDLLR  KLNYTKAEGPAKGQPKLETAIDAAEMILTLAPETNGQVAVKAWQALSEITGREHTHLALN  KEDEKIRFRDIQAQPRKIISSPTWSGLEDEHVSYNAGYTNVHELIPWRTLTGRQSLYQDH  QWMRDFGESLLVYRPPIDTRSVKAVMGEKSNGNPEKALNFLTPHQKWGIHSTYSDNLLML  TLSRGGPIVWMSEADAKDLGIEDNDWIEVFNANGALTARAVVSQRVPAGMTMMYHAQERI  VNLPGSEITGQRGGSITPSPVLRRNRPT |
| 11. | Protein_ID\|573.43724.peg.485Respiratory nitrate reductase alpha chain (EC 1.7.99.4) | AEIAKLCDLWLAPKQGTDAAMALAMGHVMLREFHLDKPSQYFTDYVRRYTDMPMLVMLEE  RDGYYAAGRTLRASDLVDSLGQENNPEWKTVAFDEKGDMTVPNGSLGFRWGDKGKWNLEQ  RDGKTGEEIELRLSLLGSHDEVANVGFPYFGGEGSEHFNKVDLENILLHKLPAKRLQLAD  GSTALVTTVYDLTMANYGLERGLNDDNCAAGYDEVKAYTPAWAEKITGVSRAHIIRTARE  FADNADKTHGRSMIIVGAGLNHWFHLDMNYRGLINMLIFCGCVGQSGGGWAHYVGQEKLR  PQTGWQPLAFALDWQRPARHMNSTSYFYNHSSQWRYETVTAQELLSPMADKSRYSGHLID  FNVRAERMGWLPSAPQLGVNPLRIADEAKKPA |
| 12. | Protein_ID\|573.43724.peg.523_Glutamyl-tRNA reductase (EC 1.2.1.70) | MLLVGAGETIELVARHLREHHVRKMVIANRTRERAQALAEEVGAEVIALSDIDERLKEAD IIISSTASPLPIIGKGMVERALKARRNQPMLLVDIAVPRDVEPEVGKLANAYLYSVDDLQ NIIQHNLAQRKAAAVQAESIVEQETSEFMAWLRAQSASETIREYRSQSEQVREELTAKAL AALEQGGDAQEIMQDLARKLTNRLIHAPTKSLQQAARDGDDERLHILRNSLGLE |
| 13. | Protein_ID\|573.43724.peg.524_Glutamyl-tRNA reductase (EC 1.2.1.70) | MTLLALGINHKTAPVALRERVTFSPETLDKALESLLAQPMVQGGVVLSTCNRTELYLSVE EQDNLQEALIRWLCNYHGLNEEDLRKSLYWHQDNDAVSHLMRVASGLDSLVLGEPQILGQ VKKPSPTPAAAILTSASWSGCSRNPFQWLSACVPKPISAPARSLWLSPPVPWRGKSLNRS PASPCYWSAPVKPLSW |
| 14. | Protein_ID\|573.43724.peg.525_Outer membrane lipoprotein component of lipoprotein transport system LolB | MNRLFRLLPLASLVLTACSLHTPQGPGKSPDSPQWRQHQQAVRSLNQFQTRGAFAYLSDE QKVYARFFWQQTGQDRYRLLLTNPLGSTELSLTAQPGSVQLIDNKGQTYTATDAEEMIGR LTGMPIPLNSLRQWIIGLPGDATDYSLDDRYRLRELNYTQNGKTWHVTYGGYTSDTQPAL PSNVELNNGAQRIKLKMDNWIVK |
| 15. | Protein_ID\|573.43724.peg.743_D-amino acid dehydrogenase (EC 1.4.99.6) | MEPALAEVSHKLTGGLRLPNDETGDCQLFTTRLAAMAEQAGVTFRFNTAVDALLHEGDRI AGVKCGDEIIKGDAYVMAFGSYSTAMLKGLVDIPVYPLKGYSLTIPIAQEDGAPVSTILD ETYKIAITRFDQRIRVGGMAEIVGFNKALLQPRRETLEMVVRDLFPRGGHVEQATFWTGL RPMTPDGTPVVGRTAYKNLWLNTGHGTLGWTMACGSGQLISDLISGRTPAIPYDDLAVARYSPGFTPARPQHLHGAHN |
| 16. | Protein_ID\|573.43724.peg.784_c-di-GMP phosphodiesterase (EC 3.1.4.52)/PdeD | MGKPCPDIHLTLRKMAASLQTIRSVALVTSGIVYCSSIFGPRQADLHRLQPALPAPRPLL LFSNDSSLLKGSPVLIQWYPAAENGLDGVMLVVNIELLGTLILNEKSALISDVSLQVGDR YFSSRHGLLEKAHVPQGTVIYRQRSTEFPFTVNINGPGASAIALEELPGELPLALIFSLL MTGIAWLATAGRMSFSREISLGISAREFALWCQPLQDARSGRCCGVEILLRWNNPRRGTI SPEVFIPIAEGNNLIIPLTRYVIAETARRLDAFPRDRHFHIAINVAARHFANGLLLRDLH NYWFSVDPVQQLVVELTERDVLQDGDQHMAEHLHFKGVQLAIDDFGTGNTSLSWLEKLRP DVLKIDRSFTSSVGIDSVNATVTDIIIALANRLHIVTVAEGVETLEQENYLRSHGVDVLQ GFYYARPMPVEAFPAWLASREESEDESETTE |
| 17. | Protein_ID\|573.43724.peg.804_Uncharacterized transporter YebQ, major facilitator superfamily (MFS) | MLNSSFKKPFHLRLIFTMDKNSSDGVPLPQRYGAILTIVLGLTMAVLDGAIANVALPTIA SDLNASPAASIWIVNAYQIAIVIALLPLSFLGDMVGYRRIYKIGLVVFIFTSLACALSRS LEMLTFARVAQGLGGAALMSVNTALIRLIYPQRFLGRGMGINSFVVAVSSAAGPTIAAAI LSLASWQWLFLINVPLGIVAFVLAMRFLPPNSARSKIIRFDLPSTIMNALTFGLLITALS GFAQGSPLSWCWRRSPLCWWWGSSSFAASSRCPSPCCRSTCCVSHSSLSLSALPSAPSAR RCWRWSPALFSAVDDGAQRSGDRSAANALAVSDHGDGAAGRLFDREMSCRAAGGYRFADY GLRPVRPGAAALVALRSGYHLAHGAVRRRLRAVSVAEQPYHRRLGSEPS |
| 18. | Protein_ID\|573.43724.peg.823_VirK | MLRTALFYNATRAMLEALSARDDFNQLLAAQATLPGKVHRQYLTRDLNAWQRAVAVINHY RYIDTLRGSRLAHAMTAVSEVPLLTLNGKEDRRFTLYASSAGKAEREGETTLWLRDSDHT LLASATFSVTRDHDAWQLVIGGLQGPRRHVSHEVIKQATRACYGLFPKRLLLEFIWQLAA RSQIAAIYGVSDNGHVFRALRYRLSKGRHFHASYDEFWQSIDGQPESPGAGGYRYVWKEN RWRVSPVKTG |
| 19. | Protein_ID\|573.43724.peg.850_Lipid A biosynthesis myristoyltransferase (EC 2.3.1.243) | MYASAPQAMVMMAELGLRDPQKILARVDWQGKAIIDEMQRNNEKVIFLVPHAWGVDIPAM LMASGGQKMAAMFHNQGNPVFDYVWNTVRRRFGGRMHARNDGIKPFIQSVRQGYWGYYLP DQDHGAEHSEFVDFFATYKATLPAIGRLMKVCRARVVPLFPVYDGKTHRLTVLVRPPMDD LLDADDTTIARRMNEEVEVFVKPHTEQYTWILKLLKTRKPGEIEPYKRKELFPKRNKKAS VRRLFYLYRAYSTVRIRVVLVLPMVLITSPWVTTTRSPETR |
| 20. | Protein_ID\|573.43724.peg.868_Isochorismatase family protein YecD | MVRVGWSADFAEALKQPVDAQAGAHTLPENWWTYPATLGKQESDIEVTKRQWGAFYGTDL ELQLRRRGIDTIILCGISTNIGVESTARNAWELGFNLVIAEDACSAASAEQHQGSMTHIF PRIGRVRSTEEILTAL |
| 21. | Protein_ID\|573.43724.peg.960_Alpha,alpha-trehalose-phosphate synthase[UDP-forming] (EC 2.4.1.15) | MLWPAFHYRLDLVSFQREAWEGYLRVNAMLADKLLPLIEPDDTLWIHDYHLLPFASELRK RGVNNRIGFFLHIPFPTPEIFNALPPHAELLEQLCDYDLLGFQTESDRTAFLDSIAMQTR LSDLGDKRYQAWGKAFSTEVYPIGIDPDEITRNAKGPLPPKLAQLKNELKNVKNIFSVER LDYSKGLPERFLAYETLLEKYPQHHGKIRYTQIAPTSRGDVQAYQDIRHQLETAAGRING QFGQLGCTPLYYLNQHFDRKLLMKVFRYSDVGLVTPLRDGMNLVAKEYVAAQDPDNPGVL VLSQFAGAAQELTSALIVNPYDRDEVAAALDRALSMPLAERIARHSAMLDVIRENDIHNWQARFVEDLQHISPRSEESRLRGR |
| 22. | Protein_ID\|573.43724.peg.1023_Outer membrane porin, OmpS | MKRKVLALMVPALLMASAANAAEIYNKNGNKLDLYGKVDGLHYFSDDASEDGDQTYVRFG LKGETQITSELTGYGQWEYNIQANTSEKEGANSWTRLGFAGLKFADCGSLDYGRNYGVVY DIESWTDMLPEFGGDTYTQTDVYMTGRTNGVATYRNSDFFGLVDGLHFALQYQGNNENAG SGEGTNNGGKRKLARENGDGFGISSYYDLDMGISFGAAYSSSDRTHNQLAAARSSQRYAN GDKADAWTVGAKYDANNIYLAAMYAETRNMTFYGNDSFGGIANKTQNFEVVAQYQFDDFN LPLRPSVAYLQSKGKDLYAYSRYGDKDLVKYVDVGMTYYFNKNMSTYVDYKINLLDEDDRFYKNSGIATDDIVALGLVYQF |
| 23. | Protein_ID\|573.43724.peg.1057_iron aquisition yersiniabactin synthesis enzyme(Irp2) @ Siderophore biosynthesis non-ribosomal peptide synthetase modules | MISGAPSQDSLLPDNRHAADYQQLRERLIQELNLTPQQLHEESNLIQAGLDSIRLMRWLH WFRKNGYRLTLRELYAAPTLAAWNQLMLSRSPENAEEETPPDESSWPNMTESTPFPLTPV QHAYLTGRMPGQKLGGVGCHLYQEFEGHCLTASQLEQAITTLLQRHPMLHIAFRPDGQQV WLPQPYWNGVTVHDLRHNDAESRQAYLDALRQRLSHRLLRVEIGETFDFQLTLLPDNRHR LHVNIDLLIMDASSFTLFFDELNALLAGESLPAIDTRYDFRSYLLHQQKINQPLRDDARA YWLAKASTLPPRPSCRWPANPPRYVKSVIPDAA |
| 24. | Protein_ID\|573.43724.peg.1062_iron aquisition yersiniabactin synthesis enzyme (Irp2) @ Siderophore biosynthesis non-ribosomal peptide synthetase modules | MREKLAHQVLNPEVWPVFDLQVGYVDGMPARLWLCLDNLLLDGLSMQILLAELEHGYRYP QQLLPPLPVTFRDYLQQPSLQSPNPDSLAWWQAQLDDIPPAPALPLRCLPQEVETPRFAR LNGALDSTRWHRLKKRAADAHLTPSAVLLSVWSTVLSAWSAQPEFTLNLTLFDRRPLHPQ INQILGDFTSLMLLSWHPGESWLHSAQSLQQRLSQNLNHRDVSAIRVMRQLAQQQNVPAV PMPVVFTSALGFEQDNFLARRNLLKPVWGISQTPQVWLDHQIYESEGELRFNWDFVAALF PAGQVERQFEQYCALLNRMAEDESGWQLPLAALVPPVKHAGQCAERSPRVCPEQSQPHIA ADESTVSLICDAFREVVGESVTPAENFFEAGATSLNLVQLHVLLQRHEFSTLTLLDLFTH PSPAALADYLAGVATVEKTKRPRPVRRRQRRI |
| 25. | Protein_ID\|573.43724.peg.1067_iron aquisition yersiniabactin synthesis enzyme (Irp1,polyketide synthetase) @ Siderophore biosynthesis non-ribosomal peptide synthetase modules | MPADQWPLFELVVSEIDDCHYRLHMNLDLLQFDVQSFKVMMDDLAQVWRGETLAPLAITF RDYVMAEQARRQTSAWHEAWDYWQEKLPQLPLAPELPVVETPPETPHFTTFNSTIGKTEW QAVKQRWQQQGVTPSAALLTLFAATLERWSRTTTFTLNLTFFNRQPIHPQINQLIGDFTS VTLVDFNFSAPVTLQEQMQQTQQRLWQNMAHSEMNGVEVIRELGRLRGSQRQPLMPVVFT SMLGMTLEGMTIDQAMSHLFGEPCYVFTQTPQVWLDHQVMESDGELMFSWYCMDNVLEPG AAEAMFNDYCAILQAVIAAPESLKTLASGIAGHIPRRRWPLNAQADYDLRDIEQATLEYPGIRQARAEITEQGALTLDIVMADDPSPSAAMPDEHELTQLALPLPEQAQLDELEATWRWL  EARALQGIAATLNRHGLFTTPEIAHRFSAIVQALSAQASHQRLLRQWLQCLTEREWLIRE  GESWRCRIPLSEIPEPQEACPQSQWSQALAQYLETCIARHDALFSGRCSPLELLFNEQHR  VTDALYRDNPASACLNRYTAQIAALCSAERILEVGAGTAATTAPVLKATRNTRQSYHFTD  VSAQFLNDARSRFHDESQVSYALFDINQPLDFTAHPEAGYDLIVAVNVLHDASHVVQTLR  RLKLLLKAGGRLLIVEATERNSVFQLASVGFIEGLSGYRDFRRRDEKPMLTRSAWQEVLV  QAGFANELAWPAQESSPLRQHLLVARSPGVNRPDKKP |
| 26. | Protein_ID\|573.43724.peg.1138_diguanylate cyclase/phosphodiesterase | MRGAVPPTVFIPVAEKIGLINALGEWVLKTACAEAASWATPLKVSVNVSPIQLMNTSLTD  TIIEVLHQTGLDPRRLDLEITESDVFNENTRSLEILSQLRELGIQISIDDFGTGYSSLSR  LSYFPFDKIKIDRSFVINIPEQKDDLDIVRLIISMGKSLHMRIVAEGVETEEQLASLQAL  GCDLVQGYLIGKPSPLR |
| 27. | Protein_ID\|573.43724.peg.1198_UDP-glucose 6-dehydrogenase (EC 1.1.1.22) | MGVYRLIMKSGSDNFRASSIQGIMKRIKAKGIPVIIYEPVMQEDEFFNSRVVRDLDAFKQ  EADVIISNRMAEELADVADKVYTRDLFGND |
| 28. | Protein_ID\|573.43724.peg.1201_Mannose-1-phosphate guanylyltransferase (EC 2.7.7.13) / Mannose-6-phosphate isomerase (EC 5.3.1.8) | MVVQTPRFNVNRITVKPGGAFSMQMHHHRAEHWVILAGTGQVTVNGKQFLLTENQSTFIP  IGAEHSLENPGRIPLEVLEIQSGSYLGEDDIIRIKDQYGRC |
| 29. | Protein_ID\|573.43724.peg.1202_Mannose-1-phosphate guanylyltransferase (EC 2.7.7.13) / Mannose-6-phosphate isomerase (EC 5.3.1.8) | MLLPVIMAGGTGSRLWPMSRELYPKQFLRLFGQNSMLQETITRLSGLEIHEPMVICNEEH  RFLVAEQLRQLNKLSNNIILEPVGRNTAPAIALAALQATRHGDDPLMLVLAADHIINNQP  VFHDAIRVAEQYADEGHLVTFGIVPNAPETGYGYIQRGVALTDSAHTPYQVARFVEKPDR  ERAEAYLASGEYYWNSGMFMFRAKNTSPSWPNSARISSKPARLR |
| 30. | Protein_ID\|573.43724.peg.1219_Low molecular weight protein-tyrosine-phosphatase (EC 3.1.3.48)/etp | MAGHLGRQFTSKLSKEYELILVMEKNHIEQISNIAPEARGKTMLFGHWLEQRDIPDPYRK  SEEAFASVFKLIEQSALLWAEKLKA |
| 31. | Protein_ID\|573.43724.peg.1248_Multidrug efflux system MdtABC-TolC, membrane fusion component MdtA | MKGSNIRRWGAALAVVIIAGAAYWFWHDSGTSGSGAPAAGQGPQGPGGARHGRFGAALAP  VQAATATEEAVPRYLTGLGTVTAANTVTVRSRVDGQLLSLHFQEGQQVKAGDLLAQIDPS  QFKVALAQAQGQLAKDQATLANARRDLARYQQLVKTNLVSRQELDTQQSLVVESAGTVKA  DEAAVASAQLQLDWTRITAPIEGRVGLKQVDIGNQISSGDTTGIVVLTQTHPIDVVFTLP  ESSIATVVQAQKAGKALSVEAWDRTNKQKISVGELLSLDNQIDATTGTIKLKARFSNLDD  ALFPNQFVNARLLVDTQQNAVVIPAAALQMGNEGHFVWVLNDENKVSKHSVTPGIQDSQK  VVISAGLSAGDRVVTDGIDRLTEGAKVEVVTASSGEQAQPDPRQSGKHGARS |
| 32. | Protein_ID\|573.43724.peg.1250_Multidrug efflux system MdtABC-TolC,inner-membrane proton/drug antiporter MdtB (RND type) | MTLAGGQRPAVRVKLNAQAIAALGLTSETVRTAITSANVNSAKGSLDGPARAVTLSANDQ  MQSAEDYRRLIIAYQNGAPIRLGDVASVEQGAENSWLGAWANQQRAIVMNVQRQPGANII  DTADSIRQMLPQLTESLPKSVKVQVLSDRTTNIRASVRDTQFELMLAIALVVMIIYLFLR  NVPATIIPGVAVPLSLVGTFAVMVFLDFSINNLTLMALTIATGFVVDDAIVVIENISRYI  EKGEKPLAAALKGAGRSASPSSPSPSR |
| 33. | Protein_ID\|573.43724.peg.1255_Multidrug efflux system MdtABC-TolC,inner-membrane proton/drug antiporter MdtC (RND type) | MKFFALFIYRPVATILISLAITLCGILGFRLLPVAPLPQVDFPVIMVSASLPGASPETMA  SSVTTPLERSLGRIAGVNEMTSSSSLGSTRIILEFNFDRDINGAARDVQAAINAAQSLLP  SGMPSRPTYRKANPSDAPIMILTLTSDTYSQGELYDFASTQLAQTIAQIDGVGDVDVGGS  SLPAVRVDLNPQALFNQGVSLDAVRTAISDANVRKPQGALEDSAHRWQVQTNDELKTAAD  YQPLIVHYQNGAAVRLGDVATVSDSVQDVRNAGMTNAKPAILLMIRKLPEANIIQTVDSI  RARLPELQQTIPAAIDLQIAQDRSPTIRASLEEVEQTLVISVALVILVVFLFLRSGRATL  IPAVAVPVSLIGTFAAMYLCGFSLNNLSLMALTIATGFVVDDAIVVLENISRHLEAGMKP  LQAALQGSREVGFTVLSMSLSLVAVFLPLLLMGGLPGRLLREFAVTLSVAIGISLAVSLT  LTPMMCGWLLKSGKPHQPTRNRGFGRLLVAVQGGYGKSLKWVLKHSRLTGLVVLGTIALS  VWLYISIPKTFFPEQDTGVLMGGIQADQSISFQAMRGKLQDFMKIIREDPAVDNVTGFTG  GSRVNSGMMFITLKPRDQRHETAQQVIDRLRKKLANEPGANLFLMAVQDIRVGGRQSNAS  YQYTLLSDDLSALREWEPKIRKALAALPELADVNSDQQDNGAEMDLVYDRDTMSRLGISV  QDANNLLNNAFGQRQISTIYQPLNQYKVVMEVDPAYTQDVSAWIRCSSLTATASRSRWPT  SPNGSRPMPRCR |
| 34. | Protein_ID\|573.43724.peg.1337_Putative multidrug resistance outer membrane protein MdtQ | MDLPAAVKNGWPQTDWWKDYHDPQLNNLIQRALANAPDMQIAEQRIRLAEAQARMSQANL  GPEMDFSADIERQRMSAEGLMGPFATDTDGNTGPWYTNGTFGLTAGWDLDLWGKNRALVK  ARIGELKAQVAEQAQTRELLSGSVARLYWQWQTEAAIKAVLQQVKNEQNNIVTVDKALYQ  RGITNSAEGAENDINVSKTDQQLADVTGTMKEIEARLMALTNSQSQSLNLKPASLPTVSA  QLPDTLGYELLARRPDLQVAHWYIEASLSEMDAAKAAFYPDINLMAFLQQDALHLSDLFR  HSAQQMGVTAGLTLPIFDSGRLNANLDIASAQNSLSIAQYNKAVVDAVNQVAKTASQVET  LMAKSQQQQQVEKDAQRVVDLAQARMAAGILPGSRVSMAKLPALQERITALRLHGQWIDA  SIQLTSALGGGYHQTVK |
| 35. | Protein_ID\|573.43724.peg.1452_DNA-binding capsular synthesis response regulator RcsB | MLRLFAEGFLVTEIAKKLNRSIKTISSQKKSAMMKLGVENDIALLNYLSSVSLSATDKE |
| 36. | Protein_ID\|573.43724.peg.1484_Uncharacterized protein STM3508 | MAFRLMRYAIAAMQRHLDKGHTQLPLVIPLLFYHGRVSPWPYPMCWLAGFADPDIARRIY  GEDFPLIDITSTPDDEIMRHRRVAMLELLQKHIRQRDLMDLHEQLVRLLALGYTSRRQLK  TLLHYLLQAGNAADPVAFCAILRKTFPGGRIRRR |
| 37. | Protein_ID\|573.43724.peg.1626_c-di-GMP phosphodiesterase (EC 3.1.4.52)/PdeA | MRCGDSVIMPDRFIPLIVQFNLSQRFDMLVLETLFSMLHKHPGQRFSVNLLPATLMQKNS  AGQIIALFQRYSVSPELITIEVTEEQAFSNADTSRQNLEALRAFGCAIAIDDFGTGYANY  ERLKHLQADIIKIDGCFVRDILTDPLDAIMVKSIVEMARAKQMSVVAEYVESEPQKARLL  ELGVNYLQGYLIGKPQPLGE |
| 38. | Protein_ID\|573.43724.peg.1717_Aminoglycosides efflux system AcrAD-TolC,inner-membrane proton/drug antiporter AcrD (RND type) | MRKTGDTNILTLAFVSTDGSMDKQDIADYVASNIQDPLSRVNGVGDIDAYGSQYSMRIWL  DPAKLNSYQMTTKDVTDAISSQNAQIAVGQLGGTPSVDKQALNATINSQSLLQTPEQFRN  ITLRVNQDGSEVTLGDVATVEMGAEKYDYLSRYNRQPASGLGIKLASGANEMATAERVIN  RLNELAQFFPHGLEYKVAYETTSFVKASITDVVKTLLEAILLVFLVMYLFLQNFRATLIP  TIAVPVVLMGTFAVLYACGYSINTLTMFAMVLAIGLLVDDAIVVVENVEHIMSEEGLSPR  EATRKSMGQIQGALVGIAMVLSAVFVPMAFFGGTTGAIYRQFSITIVAAMVLSVLVAMIL TPALCATLLKPVKPGESHERTGFFGWFNRTFNRSASRYETFVGKILHRSLRWMLIYVLLL  GGMVFLFLHLPTSFLPLEDRGMFTTSVQLPSGSTQQQTLKVVQKAEDYFLNNEKQNVESV |
| 39. | Protein_ID\|573.43724.peg.1737_hypothetical protein | MKKPTSTPHDAVFKTYLSHPDTARDFLQLYLPETLLKVCDLRTLHLESGHFVEDDLRPFY  ADILYSLKTTAGDGYIYALIEHQSTPDRHMAFRLMRYAIAAMQRHLDAGHDRLPLVIPVL  FYHGLVSPYPFSLRWLDEFIAPELAGHLYHGAFPLADITVIPDDEIATHQRMATLELLQK  HIRQRDLAQLLDKLSELLLTGLATESQVQALMHYLVQAGNTREPIKFIRELALRTPQHKE  TLMTIAEYLEQQGLERGLAQGRQEGRQEGRKEEAQRIAVAMLHSGLPRDLVARLTGLTEQ  ELTPWRTKA |
| 40. | Protein_ID\|573.43724.peg.1749_c-di-GMP phosphodiesterase (EC 3.1.4.52)/PdeF | MGGDPGIIIGLLLTLAHGMMLEQSIGVLFHFLIPCVLCWGGYRIFVPQRQQVSHGNVRLM  PHRLFWQMLLPSVIFLILSQIAEYLGLHPRTTEMTGVTPFSLRSLITFQALMVGCLTGMP  LCYFVLRVIRNPFHLRGFISQVRLQIDPKIKKIEIICWATVLILLLWLLLMPLNDSSTIF  STNYTLSLLMPVMLWGAMRFGYRFISLIWTPVLIAVIHFHYRYLPIYPSYNTQLAITSSS  YLVFSFIVAYTAMLATQQRLIYARVRQMAFLDPVVHMPNLRALSRALNGTSWSTLCFLRI  PELELLGRHYGVLLRIQYKQMLANHLRTLLQPNEAVYHLAGHDLVFRLNSEGHQARIHLI  DRSLRQFRFHWDGVPLQPRIGMSYCNVRSPVKHLYLLLGELNTIADMSLASGHPENLQRR  GAGHVQQDLKDKVVMMNRILKALEHDHFVLMAQPIQGIRGDRYHEVLVRMEGESGELTGP  NEFLPVAHEFGLSTRVDQWVIEHTLAFMDDNRRALPGLRLAINLSPVSLSRSQFPQEVEA  LLQAYNIEPWQIIFELTENYALSNPELVCQTLEHLRALGCRVAIDDFGTGYASYARLKTM  NVDILKIDGSFIRNLLASSLDYQVVDSICRLARMKNMQVVAEYVESPEIRQAVITLGIDY  MQGYDIGVPVPLTQLAEEMTGGTAKSRA |
| 41. | Protein_ID\|573.43724.peg.1875_Pyridoxine 5'-phosphate synthase (EC 2.6.99.2) | MAELLLGVNIDHIATLRNARGTAYPDPVQAAFIAEQAGADGITVHLREDRRHITDRDVRI  LRQTLHTRMNLEMAVTEEMLTIACETKPHFCCLVPEKRQEVTTEGGLDVAGQRDKMRDAC  QRLADAGILVSLFIDADEAQIKAAADVGAPYIEIHTGCYADAKTDAEQARGAGTYCQSGH  LRRQPRPEGECRPRSDLPQRESDCCAAGNA |
| 42. | Protein_ID\|573.43724.peg.1916_2-keto-3-deoxy-D-arabino-heptulosonate-7-phosphate synthase I alpha (EC 2.5.1.54) | MVSLRHALGGNPYRKRATQDRNHAKDALNNVHITDEHVLMTPEQLKAEFPLSVEQEAQIA  HARQTISDIIAGRDPRLLVVCGPCSIHDPEAAIEYARRFKALAAEVSDSLYLVMRVYFEK  PRTTVGWKGLINDPHMDGSFDVEGGLKIARRLLVELVNMGLPLATEALDPNSPQYLGDLF  SWSAIGARTTESQTHREMASGLSMPVGFKNGTDGSLATAINAMRAAAMPHRFVGINQAGQ  VCLLQTQGNPNGHVILRGGKAPNYGPEDVAKCEKEMAQAGLKPSLMVDCSHGNSNKDFRR  QPAVAESVVAQIKDGNRSIIGLMIESNIHEGNQSSEQPREAMKYGVSVTDACISWETTEA  LLRELDKDLRGHLAARLV |
| 43. | Protein_ID\|573.43724.peg.2021_RND efflux system, membrane fusion protein | MLFTIDDRTYRAALEQAQAALARAKTQASLAQSEANRTDKLVHTNLVSREEWEQRRSAAV  QAQADIRAAQAAVDAAQLNLDFTKVTAPIDGRASRALITSGNLVTAGDTASVLTTLVSQK  TVYVYFDVDESTYLHYQNLARRGQGASSDNQALPVEIGLVGEEGYPHQGKVDFLDNQLTP  STGTIRMRALLDNSQRLFTPGLFARVRLPGSAEFKATLIDDKAVLTDQDRKYVYIVDKDG  KAQRRDITPGRLADGLRIVQKGLNPGDSVIVDGLQKVFMPGMPVNAKTVAMTSSATLN |
| 44. | Protein_ID\|573.43724.peg.2022_RND efflux system, inner membrane transporter | MDFSRFFIDRPIFAAVLSILIFITGLIAIPLLPVSEYPDVVPPSVQVRAEYPGANPKVIA  ETVATPLEEAINGVENMMYMKSVAGSDGVLVTTVTFRPGTDPDQAQVQVQNRVAQAEARL  PEDVRRLGITTQKQSPTLTLVVHLFSPNGKYDSLYMRNYATLKVKDELARLPGVGQIQIF  GSGEYAMRVWLDPNKVAARGLTASDVVTAMQEQNVRCLPDSLAPSRCRRRAIS |
| 45. | Protein_ID\|573.43724.peg.2023_RND efflux system, inner membrane transporter | MGSGSYALRSQLNNKDAVGIGIFQSPGANAIDLSNAVRAKMAELATRFPEDMQWAAPYDP  TVFVRDSIRAVVQTLLEAVVLVVLVVILFLQTWRASIIPLIAVPVSVVGTFSILYLLGFS  LNTLSLFGLVLAIGIVVDDAIVVVENVERNIEEGLAPLAAAHQAMREVSGPIIAIALVLC  AVFVPMAFLSGVTGQFYKQFAVTIAISTVISAINSLTLSPALAALLLKPHGAKKDLPTRL  IDRLFGWIFRPFNRFFLRSSNGYQGLVSKTLGRRGAVFAVYLLLLCAAGVMFKVVPGGFI  PTQDKLYLIGGVKMPEGSSLARTDAVIRKMSEIGMNTEGVDYAVAFPGLNACSSPTRRIPGRSFWPETVRPAQTHGGGN |
| 46. | Protein_ID\|573.43724.peg.2024_RND efflux system, inner membrane transporter | MSGAIMQTPGMHFPISTYQANVPQLDVQVDRDKAKAQGVSLTELFGTLQTYLGSSYVNDF  NQFGRTWRVMAQADGPFRESVEDIANLRTRNNQGEMVPIGSMVNISTTYGPDPVIRYNGY  PAADLIGDADPRVLSSSQAMTHLEELSKQILPNGMNIEWTDLSFQQATQGNTALIVFPVA  VLLAFLVLAALYESWTLPLAVILIVPMTMLSALFGVWLTGGDNNVFVQVGLVVLMGLACK NAILIVEFARELEIQGKASWKRRWRRAACVCARS |
| 47. | Protein_ID\|573.43724.peg.2054_fimbrial biogenesis outer membrane usher protein | MDDSKLRYFASAWLISITLPADSAERYNAQFVNGIDPLAFNQFVASDGDVMPGTYDVNIY INDLLVDSRPVRFSEDSAHGGLAPCLSAAEYIRYGVKIDDDHQPCFALSQTIRQAEQQLD IANHRLIIHIPQQYIEHYPRDYVSPMRFDEGINAAFVNYSYSTDANNGDGGSHQYQYLSL  NSGINIASWRLRNNAYWNKFSGQADKWQSIASWAETNIIPWRSRLVVGQTSTDNSVFDSV QFRGVQLGTDAEMRPSSQTGFAPVIRGVANSNARVEVRQNNYLIYSENVPAGPFELNDISAVNRSGDFYVTVIEADGSQTTFTVAYTTLPQLVRAGQWNYQLSAGKYHDGADGYAPALMQ  SSLSYGLNNTFTLYGGALAAENYRAGAFGVGSNLGEIGALSADYTLAGTTLANGQRKQEG  ACVFCMLNPFYRARPTFKSRATVTQRQVIIASAMRLTSVDAGTMACTKMITGPLTKTKAG  RQALPNTITRPGFIIKSIALIFLPGKLSVRTALSSLTSASKITGIAQEVISLCRRVLTAL  SIT |
| 48. | Protein_ID\|573.43724.peg.2055_fimbrial biogenesis outer membrane usher protein | MNYGLYYQNTRSHFTHDDNSITLRVSIPFTLQENRRINTAFTLAHSKSSGTSGQAGVNGT  LLDDGRLSWAVTSAYDDTSHSTNSASLGYLGQYGNLYTGYAYSKSHRQASLNLSGGVVAH  RGGVTLSQPLGSTFALVEAKDAQGVGIENQTGVRIDPFGYAVVPQSVPYRVNSVALNPQD  FDAFLDVPNAVADTVPTRGAITRVRFDTFRGYSVLIHTTLADGSYPPLGAELYRASGISN  GLVGPGGDVYVSGVDSGEKLQIKWGETHQQSCEITLPELRQEPQQATAWRELSLICTVTP  RGNYAPT |
| 49. | Protein_ID\|573.43724.peg.2063_type 1 fimbriae anchoring protein FimD | MNVQSRGYVDPSRWDDGVPAAFVDYYFSGAQIKNADEGESSRSNYLNLRSGLNLGAWRLR  NISSMQYDQQRRHWDTQSIWLQRDVRSLKSLLRIGDTYTTGDVFDSIQFRGVQLMSDDEM  LPDSQRGFAPTIRGVAHSNAKVTVSQHGYVIYETFVSPGAFAISDLYPTSQSGDLEVKVT  ESNGAVRTFTQPYSAVPYMLREGRGKFSLSAGRYHSGGVGALAGISAGHFVLRFDGGIHP  LRWYAAGPGLSGVGAGPGPRFWRVRIAGRGCHPGRHPYAVGQALYGPFAARSVSEKFCQL  GHCVQPRQLSLLVKRLLRFCRSQRP |
| 50. | Protein_ID\|573.43724.peg.2073_sigma-54-dependent transcriptional regulator | MAGNPFLLAPEVNANPLLSDSWSRCQRYGLDPATEDFPRLGAGELADRLASHRGLQQLAQ  PVVEALSRQVADLQSVVILSDPDGLVLHTLGDTQALLQKAQRVALAPGNLWSESGRGTNA  IGTALAIDDGCEIDGRQHFLTRNQNLYCAAMPLQRPDGSIAGVLDISGPANFPHQHTFGW  VKAAAKQIEYLWVKQSLHPEQWLMSLHRQVDKLDSVEELLLVFSDNVLTAGNRLAMREFG  LSAAQFGQLTFASLFPTLTQTAVSVPLPLTTPQGRYHYRLRAPTRRRVAVSAPPAMHLPF  TSPREGEKLLRLLNAGIALCIEGETGSGKEYVSRTLHRHSRWRSGKFVAINCAAIPESLI  ESELFGYQPGAFTGASKNGYIGKIREADGGVLFLDEIGDMPLALQTRLLRVLQEKEVAPL  GASRSVPVNFALICATHRNLTQRVSAGEFREDLLWRLREYALALPPLREWPALETFIATL  WHDLGGASRRVTLSNALLVHLSQLPWPGNVRQLQSVLKVMLALADEGDTLTPDALPEAYR  AAPAPLPRGDCRPMMSS |
| 51. | Protein_ID\|573.43724.peg.2078_Rhodanese-domain-containing inner membrane protein YgaP | MTIATLTPAEAQAQIAQGARLIDIRDADEYTREHIPDAELVPLATLTSGAPLNARAGETV  IFHCQAGSRTQNNAIRLAAAAAPAQTCLLAGGIQAWKAAGLPVVEDSSQPLPLMRRCKLP  LAS |
| 52. | Protein_ID\|573.43724.peg.2107_Multidrug efflux system EmrAB-OMF, membrane fusion component EmrA | MSANAESQTPQQPGSKKGKRKGALLLLTLLFIIIAVAYGIYWFLVLRHYEETDDAYVAGN  QVQIMAQVAGSVTKVWADNTDYVQKGDPLVTLDRTDAQQAFEKAKTQLAASVRQTRQQMI  NSKQLQANIDVKKTALAQAQADLNRRIPLGAANLIGREELQHARDTVASAQAELDVAIQQ  YNANQAIVLGTRLEQQPAVLQAATEVRNAWLALQRTQIVSPISGYVSRRSVQPGAQIGTT  TPLMAVVPATNLWIDANFKETQLAHMRIGQPATVISDIYGDDVKYTGKVVGLDMGTGSAF  SLLPAQNATGNWIKVVQRLPVRIELDEKQLAEHPLRIGLSTLVEVNTTDRDGEMLASQVR  SSRFTRAMPAKSLSIRLIS |
| 53. | Protein_ID\|573.43724.peg.2120_Carbon storage regulator | MLILTRRVGETLMIGDEVTVTVLGVKGNQVRIGVNAPKEVSVHREEIYQRIQAEKSQQSS  Y |
| 54. | Protein_ID\|573.43724.peg.2138_Anaerobic nitric oxide reductase transcription regulator NorR | MLISGETGTGKELVAKAVHQGSPRAANPLVYLNCAALPESVAESELFGHVKGAFTGAISN  RSGKFEMADNGTLFLDEIGELSLALQAKLLRVLQYGDIQRVGDDRSLRVDVRVLAATNRD  LRQEVVEGRFRADLYHRLSVFPLSVPPLRERESDVVLLAGYFCEQCRLRMGLARVILAEA  ARNRLQQWSWPGNVRELEHAIHRAVVLARATQAGDEVVLEPQHFQFAVEAPMLPTETAAA  APATGNINLREATDSFQREAISRALEANQGNWAATARALELDVANLHRLAKRLGLKGSPP  GKSSAG |
| 55. | Protein_ID\|573.43724.peg.2176_Formate hydrogenlyase transcriptional activator | MSYTPMSDLGQQGLFDITRLLLQQPDLAALSETLTRLVQQSALADEAAIILWNAGNHRAA  RYACDEAGHPVSYEDETVLAHGPVRRLLSRPDALHCDHETFADTWPQLIRSGLYRPFGYY  SLLPLAADGRIFGGCEFLRRDNRPWSEKEFQRLHTFAQIVAVVTEQIQNRVSNNVDYDLL  CHERDNFRILVAITNAVLSRLDIDELVSEVAKEIHRYFRIDAISVVLRSDRKGKLNIYST  HYLDASHPVHDQSEVDEAGTLTERVFKSKEMLLLNLHEHDTLAPYEKMLFEMWGNKIQTL  CLLPLMSGNTLLGVLKLAQCDEQVFTTTNLKLLRQIAERVSIAIDNALAYREIQRLKERL  VDENLALTEQLNNVESEFGEIIGRSEAMNNVLKQVEMVAHSDSTVLILGETGTGKELIAR  AIHNLSGRNGRRMVKMNCAAMPAGLLESDLFGHERGAFTGASAQRIGRFELADKSSLCLD  EVGDMPLELQPKLLRVLQEQEFERLGSNKLIQTDVRLIAATNRDLKQMVIDREFRSDLYY  RLNVFPIHLPPLRERPDDIPLLVKAFTFKIARRMGRNIDSIPAETLRTLTRMEWPGNVRE  LENVIERAVLLTRGNVLQLSLPERDIVEAPRTPAVLPEEGEDEYQLIVRVLKESNGVVAG  PKGAAQRLGLKRTTLLSRMKRLGINKDELV |
| 56. | Protein_ID\|573.43724.peg.2294_Acyl-CoA dehydrogenase; probable dibenzothiophene desulfurization enzyme | MTLLSTGTDYDALAAAFRPIFTRIAQGAAEREQQRILPEEPIRWLKEAGFGTLRIPREKG  GWGASLPQLSALLIELAQADSNLPQALRAHFAFVEDQLNQPDSAGRDRWFRRFLDGELVG  SGWTEIGAVKLGEVNTRVTPTEGGWRLDGEKFYSTGALYADWIDVFARRSDTASDVIALV  STQQTGVVREDDWDGFGQRLTGSGTTRFTGARVEPDHVYDFARRFRYQTAFYQHVLLATL  AGIGLAVERDAAQGVKQRSRMYSHGNAAVPRDDAQVLQVVGQISSWAWATRAAVLQAAES  LQQAYVAHISDDDALIARRNQQAEVEAAQAQVIASDWIPRAATELFNALGASDTRTRLAL  DRHWRNARTVASHNPVIYKARNIGNWLVNGEAPTFIWQIGNGEKTAG |
| 57. | Protein_ID\|573.43724.peg.2331_Glycine cleavage system transcriptional activator GcvA | MEKLYAEYLLPVCSPLLLTGDKALKTPADLAQHTLLHDASRRDWQTYTRQLGLSHINVQQ  GPIFSHSAMVLQAAIHGQGVALANNVMAQSEIEAGRLVCPFNDVLVSKNAFYLVCHDSQA  ELGKIAAFRQWILAKAASEQEKFRFRYEQ |
| 58. | Protein_ID\|573.43724.peg.2350_VgrG protein | MDVQNFDHSHHALKIRGLSSGVDVLSFEGKEQLSAPFRYDIQFTSSDKSIAPESVLMQDG  AFSLTAPPVQGIPTQTPLRTLYGVITGFKQLSSSRDEARYEVRLEPRLALLSRSHQNAIY  QNQTVPQIVEKILRERHEMRGQDFVFNLKGDYPSREQVMQYDEDDLTFISRLLSVVGIWF  RFSTDARLKIGVIEFYDDQSGYERGLTLPLRHPSGMSDSGTEAVWDLNTAYSVVSRSVTT  RDYNYREAMAEMTTGQFDVTGGDNTTYGEAYHYADNFLKTGDKATPESGAFYARIRHERY  LNGRAILKGQSTSSLLMPGLEIKVEGNDAPEVFRKGILITGITASAARDRSYELAFTAIP  YSERYGYRPPLIRRPVMAGTLPARVTSTTANDVYAHIDKDGRYRVNLDFDRDTWKPGFES  LWVRQSRPYAGDTYGLHLPLLAGTEVSIAFEDGNPDRPYIAGVKHDSAHTDHVTIQNYKR  NVRAPRRITKSGWTMSGVRSTSRSAQSTAERAS |
| 59. | Protein_ID\|573.43724.peg.2533_Outer membrane usher protein fimD precursor | MTFTVNQNLPDGWGGFYLSGRISDYWNRSGTEKQYQVSYNNSFGRLSWSASAQRVYTPDS  SGHRRDDRISLNFSYPLWFGDNRTANLTSNTSFNNSRFASSQIGINGSLDSENNLNYGVS  TTTATGGQHDVALNGSYRTPWTTLNGSYSQGEGYRQSGIGASGTMIAHSGGVVLSPESGS  TMALIEAKDAAGAMLPGSPGTRVDSNGYAILPYLRPYRINTVEIDPKGSHDDVAFDRTVA  QVVPWEGSVVKVAFGTKVQNNLTLQARQANHEPLPFAASIFSPTARRSALSARAA |
| 60. | Protein_ID\|573.43724.peg.2534_Outer membrane usher protein fimD precursor | MKQRSFCPGRLSTAIAIALCCFPPFSSGQENPGTVYQFNDGFIVGSREKVDLSRFSTSAI  TEGTYSLDVYTNDEWKGRYDLRIARDKDGRLGVCYTKAMLAQYGIAAEKLNPQLSEQEGY  CGSLKSWRNEENVKDNLVQSSLRLNISVPQIYEDQRLKNYVSPEFWDKGITALNLGWMAN  AWNSHTSSVGGSDNSSAYLGVNAGLSWDGWLLKHIGNLNWQQQQGKAHWNSNQTYLQRPI  PQLNSIVSGGQIFTNGEFFDTIGLRGVNLSTDDNMFPDGMRSYAPEIRGVAQSNALVTVR  QGSNIIYQTTVPPGPFTLQDVYPSGYGSDLEVSVKEADGSVEVFSVPYASVAQLLRPGMT  RYALSAGKVDDSALRYKPMLYQATWQHGINNLLTGYTGVTGFDDYQAFLVGTGMNTGIGA  LSFDVTHSRLKSDAHDDSGQSYRATFNRMFTDTQTSIVLAAYRYSTKGYYNLNDALYAVD  QEKNSRSNYTLWRQKTA |
| 61. | Protein_ID\|573.43724.peg.2551_Chaperone protein FimC | MRKTATMAHGLLAGCVLFAASIFSASAQAGVALGATRVIYPAGQKQVQLAVTNNDDNSTW  LIQSWVENADGQRDGRFVITPPLFAMQGKKENTLRIIDATNNQLPQDRESLFWMNVKAIP  SMDKSKLSDNTLQLAIISRIKLYYRPGKLALPPDQAAEKLTFSRSGSSLTLTNPTPYYLT  VTELNAGTRILENALVPPMGKTSVKLPADAGNTITYRTINDYGALTPKMNGVLR |
| 62. | Protein_ID\|573.43724.peg.2553_Outer membrane usher protein FimD | MPWSTVPVLQREGHTRFALTAGEYRSGNSQQETPDFFQGTVMHGLPAGWTLYGGTQLADR  YRAFNLGVGKNMGYFGALSLDITQANATLADDSEHQGQSVRFLYNKSLDETGTNLQLVGY  RYSTRGYYNFADTTYRRMSGYSVETQDGVIQVKPKFTDYYNLAYSKRGKVQLSVTQQLGR  TATLYLSGSHQTYWGTDDADEQLQAGLNAAVDDINWSLSYSLTKTPGSRGAIKCWRSISI  SPSATGCVPTAARSGGTPAPATACRTISMAA |
| 63. | Protein_ID\|573.43724.peg.2554_Outer membrane usher protein FimD | MTNLAGLYGTLLEDNNLSYSVQTGYAGGGNGDNGSTGYTALNYRGGYGNANVGYSRSDGF  KQLYYGVSGGVLAHANGITLSQPLNDTVVLVKAPGAGGVKVENQTGVRTDWRGYAVLPYA  TEYRENRIALDTNTLADNVDLDDAVVSVVPTHGAIVRANFNAQVGMKILMTLTHRGKPVP  FGALATGDSNQSGSIVADNGQVYLSGMALAGKVRVKWGDGPDAQCVADYRLPPESQQQAL  SQLSVACR |
| 64. | Protein_ID\|573.43724.peg.2665_Biosynthetic arginine decarboxylase (EC 4.1.1.19) | MLRTYNIAWWGNNYYDVNELGHISVCPDPDVPEARVDLAELVKAREAQGQRLPALFCFPQ  ILQHRLRSINAAFKRARESYGYNGDYFLVYPIKVNQHRRVIESLIHSGEPLGLEAGSKAE  LMAVLAHAGMTRSVIVCNGYKDREYIRLALVGEKMGHKVYLVIEKMSEIAIVLEEAERLN  VVPRLGVRARLASQGSGKWQSSGGEKSKFGLAATQVLQLVEILREAGHLESLQLLHFHLG  SQMANIRDIATGVRESARFYVELHKLGVNIQCFDVGGGLGVDYEGTRSQSDCSVNYGLNE  YANNIIWAIGDACEENGLPHPTVITESGRAVTAHHTVLVSNIIGVERNEYTEATPPAEDA  ARPLQSMWETWLEMHETGNRRSLREWLHDSQMDLHDIHIGYSSGTFNLQERAWAEQLYLN  MCHEVQKQLDPSNRAHRPIIDELQERMADKIYVNFSLFQSMPDAWGIDQLFPVMPLEGLN  KSPERRAVLLDITCDSDGAIDHYVDGDGIATTMPMPEYDPENPPMLGFFMVGAYQEILGN  MHNLFGDTEAVDVFVFPDGSVEVELSDEGDTVADMLQYVQLDPNTLLTQFRDQVKNTGLD  DALQQQFLEEFEAGLYGYTYLEDE |
| 65. | Protein_ID\|573.43724.peg.2713_hypothetical protein | MQEGVSPWSLQRRRNSFQTQLSQQLGDWGTMYFRASRDDYWGGERTLTGMSLGYSNSLKG  VSYGVNYNIDRTKDANGNWPENRQISFNVSVPFSIFGYSRNLQSMYATTTLTHDNTGRTL  SQTGLSGNTMDGKLSYSASQSWGNQGQISNTNLNTGYQGSKGSISGGYSYSSDMQAINMS  ASGGVMVHSGGITLSRAMGDSVALVSAPGAAGVSVNGGTAVTDWRGYAVVPYLTDYTRNS  VGVDPSTLPENVDLTQTNLNVYPTKGAVVKANFATRVGYQVLMTLKLDNGVVPFGAVATL  LNAGMAEVNSSIVGDDGQVYLTGLPERGSYWLSGARQQHVSAVSVLTSPVYRRRRINRSV  R |
| 66. | Protein_ID\|573.43724.peg.2714_hypothetical protein | MNVRQLPDLKDLPATAEIDNIGALIPQATTMLDLARLRLDISVPQAAMQPEVRGAVDPSQ  WEEGISALMANYSLSAGRTTNSGQNQTSHNNNLFATVRAGANTGPWRLRSTMTHTRVENN  GGNNALTTTQTRFSNTYLARDIRGWRSNLLMGESSTGSDVFDGIPFRGVKLSSNEQMLPS  QLRGYAPAISGVANSNARVTVRQNGNVVYETYVAPGPFYINDIQQAGLSGDYDVKVTEAD  GTERQFIVPYSSLPVMLRPGGWKYELTAGQYDGNLTDGSRRADFMLGTVVYGLPGDVTLF  GGILAAKDYQAFNIGTGVSLDTSVRSRLISRIHRRSLITNPP |
| 67. | Protein_ID\|573.43724.peg.2753_TonB-ExbBD energy transducing system, ExbB subunit | MGNNLMQADLSVWGMYHHADIVVKVVMIGLILASVVTWAIFFGKGAEILASKRRLKREQQ  QLAEARSLDQASDIASAFEAKSLTTQLINEAQNELELSAGAEDNEGIKERTGFRLERRVA  AVGRHMGRGNGYLATIGAISPFVGLFGTVWGIMNSFIGIAQTQTTNLAVVAPGIAEALLA  TAIGLFAAIRRWSSITSSPA |
| 68. | Protein_ID\|573.43724.peg.2795_Outer membrane channel TolC (OpmH) | MKKLLPILIGLSLTGFSAMSQAENLLQVYQQARISNPDLRKSAADRDAAFEKINEARSPL  LPQLGLGADYTYTNGYRDSNGVNSNVTSGSLQLTQVLFDMSKWRALTLQEKTAGIQDVTY  QTDQKL |
| 69. | Protein_ID\|573.43724.peg.2796_Outer membrane channel TolC (OpmH) | MLAAIDTLSYTEAQKQAIYRQLDQTTQRFNVGLVAITDVQNARSQYDAVLANEVTARNDL  DNAVEGLRQVTGNYYPELASLNVNGFKTNKPQAVNALLKEAENRNLSLLQARLSQDLARE  QIRQAQDGHLPTLNLSASTGVSNTRYNGSKTNTPLAYNDSDNGQNQIGLNFSLPLYQGGA  VTSQVKQAQYNFVGASEQLESAHRSVVQTVRSSFNNVNASISSINAYKQAVVSAQSSLDA  MEAGYSVGTRTIVDVLDATTTLYNAKQQLSNARYNYLINELNIKSALGTLNEQDLVALNN  TLGKPISTSADSVAPENPQQDATADGYGNTTAAVKPASARTTQSSGSNPFRQ |
| 70. | Protein_ID\|573.43724.peg.2809_D-glycero-beta-D-manno-heptose 1-phosphate adenylyltransferase (EC 2.7.7.70) | MKVTLPEFERAGVLVVGDVMLDRYWYGPTSRISPEAPVPVVKVENIEERPGGAANVAMNI  ASLGATSRLVGLTGIDDAARALSQALANVNVKCDFVSVPTHPTITKLRVLSRNQQLIRLD  FEEGFSGVDPQPMHERIQQALGSIGALVLSDYAKGALTSVQTMIRLAREAGVPVLIDPKG  TDFERYRGATLLTPNLSEFEAVVGKRAG |
| 71. | Protein_ID\|573.43724.peg.2825_Urease gamma subunit (EC 3.5.1.5) | MLFTAALVAERRLARGLKLNYPESVALISAFIMEGARDGKSVASLMEEGRHVLTREQVME  GVPEMIPDIQVEATFPDGSKLVTVHNPII |
| 72. | Protein_ID\|573.43724.peg.2826_Urease beta subunit (EC 3.5.1.5) | MIPGEYHVKPGQIALNTGRATCRVVVENHGDRPIQVGSHYHFAEVNPALKFDRQQAAGYR LNIPAGTAVRFEPGQKREVELVAFAGHRAVFGFRGEVMGPLEVNDE |
| 73. | Protein_ID\|573.43724.peg.2863_Phosphoenolpyruvate-dihydroxyacetone phosphotransferase operon regulatory protein DhaR | MTLPALLRRAIKHARGLNHVEVTFESQHQFVDAVITLKPIVEAQGNSFILLLHPVEQMRQ  LMTSQLGKVSHTFEQMSADDPETRRLIHFGRQAARGGFPVLLCGEEGVGKELLSQAIHNE  SERAGGPYIAVNCQLYADSVLGQDFMGSAPTDDENGRLSRLELANGGTLFLEKIEYLAPE  LQSALLQVIKQGVLTRLDARRLIPVDVKVIATTTVDLANLVEQNRFSRQLYYALHSFEIV  IPPLRARRNSIPSLVHNRLKSLEKRFSSRLKVDDDALAQLVAYSWPGNDFELNSVIENIA  ISSDNGHIRLSNLPEYLFSERPGGDSASSLLPASLTFSAIEKEAIIHAARVTSGRVQEMS  QLLNIGRTTLWRKMKQYDIDASQFKRKHQA |
| 74. | Protein_ID\|573.43724.peg.2945_DnaA initiator-associating protein DiaA | MLDRIKACFTESIQTQIAAAEALPDAISRAAMTLVQSLLNGNKILCCGNGTSAANAQHFA  ASMINRFETERPGLPAIALNTDNVVLTAIANDRLHDEIYAKQVRALGHAGDVLLAISTRG  NSRDIVKAVEAAVTRDMTIVALTGYDGGELAGLLGQQDVEIRIPSHRSARIQEMHMLTVN  CLCDLIDNTLFPHQDD |
| 75. | Protein_ID\|573.43724.peg.3031_RNA polymerase sigma-54 factor RpoN | MKQGLQLRLSQQLAMTPQLQQAIRLLQLSTLELQQELQQALESNPLLEQTDLHDEVEAKE  VEDRESLDTVDALEQKEMPDELPLDASWDEIYTAGTPSGNGVDYQDDELPVYQGETTQTL  QDYLMWQVELTPFTDTDRAIATSIVDAVDDTGYLTIQIEDIVDSIGDDEIGLEEVEAVLK  RIQRFDPVGVAAKDLRDCLLIQLSQFAKETPWLEEARLIISDHLDLLANHDFRTLMRVTR  LKEEVLKEAVNLIQSPGSSPWPVHSDQRTGICHPGCAGAQGERPLDRRAQRRQHSTVENK  SAVCGNGQQRPQ |
| 76. | Protein_ID\|573.43724.peg.3033_RNA polymerase sigma-54 factor RpoN | MKPMVLADIAQAVEMHESTISRVTTQKYLHSPRGIFELKYFFSSHVNTEGGGEASSTAIR  ALVKKLIAAENPAKPLSDSKLTSMLSEQGIMVARRTVAKYRESLSIPPSNQRKQLV |
| 77. | Protein_ID\|573.43724.peg.3105_RNase E specificity factor CsrD | MRLTTKFSAFITLLTSLTIFVTLIGASLSFYNGIENKVENRVQAVATMLDNRLITTSFDK  LEPQLDELMTPIEIVHIDFMLNGKPLYSHSRPDSYRPLGSHEQFREITVPSLKHPGITLH  LVYVDPMVNYFRSLSITAPLSISIGFMVVIIFFAVRWIRRQLAGQELLELRSTRILSGER  GPQVRGSVYEWPASTSSALDMLLSELQFASDQRSRMDTLIRSYAAQDSKTGLNNRLFFDN  QLATLLEDQEKVGAYGIVMMIRLPEFDLLRDNWGRAAAEEHYFTLINLLSTFIMRYPGAL  LARYHRSDFAVLLPHRTLKEADSIAGLLLKAMDALPPTRILDRDDMMHIGICSFRSGQSA  AQVMEHAEAATRNAVLQGSNSWSVYDDTLPEKGRGNVRWRTLIEQMLSRGGPRLYQKPAV  TRDGRVHHRELMSRMYDGKEEVIAAEYMPMVLQFGLAEEYDRLQVTRLLPFLGFWPEENL  ALQLSVESLIRPRFQRWLRDALMQCEKSQRQRIIFELAEADVCQYIGRLQPVMRLVNALG  VRVAVVQAGLTLVGTSWIKQLDAELIKLHPGLARNIEKRSENQLLVQSLVEACKGMPMQV  LRRAYAAAASGWYCPNAVSRAGRGNFCRLTAT |
| 78. | Protein_ID\|573.43724.peg.3120_DNA-binding protein Fis | MFEQRVNSDVLTVSTVNSQDQVTQKPLRDSVKQALKNYFAQLNGQDVNDLYELVLAEVEQ  PLLDMVMQYTRGNQTRAALMMGINRGTLRKKLKKYGMN |
| 79. | Protein_ID\|573.43724.peg.3122_Multidrug efflux system AcrEF-TolC, membrane fusion component AcrE | MTTHARVTLLSGLIISALLLTGCDNSDNQQPHAQAPQVTVHVVNSAPLSVTTELPGRTSA  FRVAEVRPQVSGIILKRNFVEGSDVEAGQSLYQIDPATYQAAWNSAKGDEAKAEAAAAIA  HLTVKRYVPLLGTKYISQQEYDQAVATARQADADVIATKAAVETARINLAYTKVTSPLAG  ASVNRASPKGRWSPTASRMPCLPSSSSIPSMLT |
| 80. | Protein_ID\|573.43724.peg.3128_Multidrug efflux system AcrEF-TolC, inner-membrane proton/drug antiporter AcrF (RND type) | MAFVSLKPWEARSGDKNSVESIIKRATVAFSQIKDAMVFPFNMPAIIELGTATGFDFELI  DQGGLGHTALTQARNQLLGMVKQHPDQLVRVRPNGLEDTPQFKLDVDQEKAQALGVSLSD  INETISAALGGYYVNDFIDRGRVKKCTFRLMPTSVCCRATLTTCMFVAPTAKWCRSPPLS  LHAGYMARRAWSATTGCHLWRSSVKLRQAKVPGKPWR |
| 81. | Protein_ID\|573.43724.peg.3129_Multidrug efflux system AcrEF-TolC,inner-membrane proton/drug antiporter AcrF (RND type) | MALMETLASKLPSGIGYDWTGMSYRERLSGNQAPALYAISLIVVFLCLAALYESWSIPFS  VMLVVPLGVIGALLAATLRGLNNDVYFQVGLLTTIGLSAKNAILIVEFAKDLMEKEGKGI  IEATLEASRMRLRPILMTSLAFILGVMPLVISHGAGSGAQNAVGTGVMGGMLTATLLAIF  FVPVFFVVVRRRFTRHAE |
| 82. | Protein_ID\|573.43724.peg.3466_Cellulose synthase catalytic subunit [UDP-forming] (EC 2.4.1.12) | MTFCSLVLALMCITQPFNPLSQFIFLMLLWGVALLVRRIPGRFSALMLIVLSLTVSCRYI  WWRYTSTLNWNDPVSLVCGIILLFAETYAWVVLVLGYFQVVWPLNRQPVPLPEDMDLWPT  VDIFVPTYNEDLNVVKNTIYASQGIDWPKDKLNIWILDDGGREAFRQFAKDVGVHYIART  SHEHAKAGNINNALKYAKGEFVSIFDCDHVPTRSFLQMTMGWFLKEKELAMMQTPHHFFS  PDPFERNLGRFRKTPNEGTLFYGLVQDGNDMWDATFFCGSCAVIRRGPLDEIGGIAVETV  TEDAHTSLRLHRRGYTSAYMRIPQAAGLATESLSAHIGQRIRWARGMVQIFRLDNPLFGK  GLKLVQRVCYANAMLHFLSGIPRLIFLTAPLAFLLLHAYIIYAPALMIALFVLPHMIHAS  LTNSKIQGKYRHSFWSEIYETVLAWYIAPPTFVALINPHKGKFNVTAKGGLVEEEYVDWV  ISRPYIYLVLLNLVGVAVGIWRFMYGPENEILTVWVSIVWVFYNLIILGGAVAVSVESKQ  VRRSHRVEMSMPAAIAREDGHLFSCTVHDYSDGGLGIKINGDAQVLEGQNARLLLKRGQQ  EYAFPVRVARVNGSEVGLQLLPLTNQQHIDFVQCTFARADTWALWQDSFPEDKPMESLLD  ILKLGFRGYRHLAEFSPPSVKVVFRALTSLVAWIASFVPRRPERAAPTLSADPAMAQQ |
| 83. | Protein_ID\|573.43724.peg.3515_Biotin sulfoxide reductase (EC 1.-.-.-) / Free methionine-(S)-sulfoxide reductase | MRVSWEQALDLIDAQHRRIRDSYGPASIFAGSYGWRSNGVLHKAATLLQRYMSLAGGYTG  HLGDYSTGAAQAIMPYVVGGNEVYQQQTSWPLVLEHSEVVVLWSANPLNTLKIAWNASDE  QGIPWFDRLRQSGKRLICIDPMRSETVDFFGDSMEWIAPHMGTDVALMLGIAHTLVENDW  QDDAFLTRCTSGYDIFARYLTGESDGVAKTAEWAAAICGVKADKIRELAKLFHENTTMLM  AGWGMQRQQYGEQKHWMLVTLAAMLGQIGTPGGGFGLSYHFANGGNPTRRAAVLGSMQGS  VAGGVDAVEKIPVARIVEALENPGAEYQHNGMARRFPDIRFIWWAGGANFTHHQDTNRLI  QAWQKPELIVISECFWTAAARHADIVLPATTSFERNDLTMTGDYSNQHLVPMKQVVAPRD  EARDDFAVFADLSERWEAGGRERFTEGKSDLQWLETFYQMAARRGAQQQVTLPPFAEFWQ  ANQLIEMPEEPENARFARFAAFRADPRPTR |
| 84. | Protein_ID\|573.43724.peg.3607_ADP-heptose--lipooligosaccharide heptosyltransferase II | MAPAWCRPLLSRMPEVNEAIPMPLGHGALAIGERRKLGHSLRERRYDRAYVLPNSFKSAL  VPFFANIPLRTGWRGEMRYGLLNDARVLDKDAWPLMVERYVALAYDNGVMRCAKDLPQPL  LWPQLQVNEGEKSQACSAFNLSYDRPIVGFCPGAEFGPAKRWPHYHYAALAKKLIDDGYQ  IALFGSAKDNEAGKEIIAALSSEQQAWCRNLAGETQLEQAVILIAACKAVVTNDSGLMHV  AAALDRPLVALYGPSSPDFTPPLSHKARVIRLITGYHKVRKGDAAEGYHQSLIDITPERV  LQELNELLAEKTEHEEA |
| 85. | Protein_ID\|573.43724.peg.3608_Lipopolysaccharide core heptosyltransferase I | MRVLIVKTSSMGDVLHTLPALTDAAQAIPGIRFDWVVEEGFAQIPSWHKSVERVIPVAIR  RWRKAWFSAPIKAERQAFREAVQAVKYDAIIDAQGW |
| 86. | Protein_ID\|573.43724.peg.3609_Lipopolysaccharide core heptosyltransferase I | MDWQTAREPLASLFYNRRHHIAKQQHAVERTRELFAKSLGYAKPQAQGDYAIARHFLQHE  ASAAAPYLVFLHATTRDDKHWPETRWQELLDLLADSGVHIKLPWGPRTKRRGRSGWRKAG  SMLKCCRA |
| 87. | Protein_ID\|573.43724.peg.3610_Lipopolysaccharide core heptosyltransferase I | MLPRMSLEQVAQVLAGARAVVSVDTGLSHLTAALDKPNFTLYGPTDPGLIGGYGKNQHIV  RPENSASTGDIAASRIHLLLQNQGLL |
| 88. | Protein_ID\|573.43724.peg.3618_Lipopolysaccharide core heptosyltransferase III | MTPETLSRGPLNPARIPVIKLRHHGDMLLITPLIHALKQQYPAASVDVLLYEETRDMLAA  NPDIHHIYGLDRRWKKQGKRYQLKMQWQLIQTLRQQRYDMVLNLADQWPSAVISKLTGAA  TRIGFDFPKRRHPVWRYCHTALASTQQHNQLHTVQQNLSILAPLGLQLNDAPARMGYSEA  DWAASRALLPEDFREHYIVIQPTSRWFFKCWREDRMSALINALSAEGYAVVLTSGPDARE  KKMVDTIIAGCPQARLHSLAGQLTLRQLAAVIDHARLFIGVDSVPMHMAAALGTPLVALF  GPSKLTFWRPWQAKGEVIWAGDFGPLPDPDAINTNTDERYLDLIPTDAVIAAAKKVLA |
| 89. | Protein_ID\|573.43724.peg.3623_3-deoxy-D-manno-octulosonic acid transferase(EC 2.4.99.12) | MELLYTTLLYLIQPLVWLRLLLRSRKAPAYRKRWAERYGFCQNKVEPDGILLHSVSVGET  LAAIPLVRALRHRYPSLPITVTTMTPTGSERAMSAFGKDVHHVYLPYDLPGAMNRFLNTV  QPKLVIVMETELWPNMVAALHKRKIPLVIANARLSERSAKGYAKLGGFMRRLLSRITLIA  AQNEEDGNRFLSLGLKRNQLAVTGSLKFDISVTPELAARAVTLRRQWAPHRKVWIATSTH  DGEEQIILQAHKNCWRLSRTCC |
| 90. | Protein_ID\|573.43724.peg.3624_3-deoxy-D-manno-octulosonic acid transferase(EC 2.4.99.12) | MSFTLRSTGEIPSSSTQVVIGDTMGELMLLYGIADLAFVGGSLVERGGHNPLEPAAHAIP  VLMGPHTFNFKDICAKLQQDDGLITVTDADSLVREVSTLLTDEDYRLWYGRHAVEVLHQN  QGALSRLLQLLQPYLPQRSH |
| 91. | Protein_ID\|573.43724.peg.3675_Sulfate-binding protein Sbp | MNKWGVGLTLLLASASVLAKDIQLLNVSYDPTRELYEQYNKAFSAHWKQETGDNVVIRQS  HGGSGKQATSVINGIEADVVTLALAYDVDAIAERGRIDKNWLKRLPDNSAPYTSTIVFLV  RKGNPKQIHDWNDLIKPGVSVITPNPKSSGGARWNYLAAWGYALHQNHGDQAKAQEFVKA  LYKNVEVLDSGARGSTNTFVERGIGDVLIAWKTKRCWRPTNWAKTSLKS |
| 92. | Protein_ID\|573.43724.peg.3726_RND efflux system, membrane fusion protein | MKYIIAPIATALFLLSGCDNAQTSAPQQPTPEVGVVTLQSQPVPVVSQLTGRTTASLSAE  VRPQVGGIIQKRLFTEGDMVKAGQALYQIDPSSYRATWNEAAAALKQAQALVVSDCQKAQ  RYASLVRDNGVSRQDADDAASTCAQDKASVESKKAALESARINLNWTTVTAPIAGRIGIS  SVTPGALVSADQDTALATIRGLDTMYVDLTRSSVDLLRLRKQSLASNSDTLSVTLTLEDG  STYPEKGRLALTEVAVDESTGSVTLRAIFPNPQHVLLPGMFVRARIDEGIMNDAILAPQQ  GITRDAKGDATALVVDAANKVEQRTVETGDTYGDKWLVLSGLKAGDKLIVEGTGKVAPGQ  TVKAVAVNNNGGNA |
| 93. | Protein_ID\|573.43724.peg.3727_RND efflux system, inner membrane transporter | MFSRFFVRRPVFAWVIAILIMLAGVLAIRTLPVGQYPDVAPPAVKISATYTGASAETLEN  SVTQVIEQQLTGLDHLLYFSSTSSSDGSVSITVTFEQGTDPDTAQVQVQNKVQQAESRLP  SEVQQSGVTVEKSQSSFLLILAVYDKTNRATSSDISDWLVSNMQDPLARVEGVGSLQVFG  AEYAMRVWMDPTKLASYSLMPSDVQSAIEAQNVQVSAGKIGALPSSNAQQLTATVRAQSR  LQTPDQFKAIIVKSQADGSVVRLSDVARVEMGSEDYTATANLNGHPAAGIAVMMAPGANA  LDTATLVKSKIAEFQRQMPQGYDIAYPKDSTEFIKISVEDVIQTLFEAIILVVCVMYLFL  QNFRATLIPAVAVPVVLLGTFGVLALFGYSINTLTLFAMVLAIGLLVDDAIVVVENVERI  MRDEGLPAREATEKSMGEISGALVAIALVLSAVFLPMAFFGGSTGVIYRQFSVTIISAMM  LSVVVALTLTPALCGALLSHSKPHTKGFFGAFNRLWGRTEAGYQRRVLGGLRRGAVMMGA  YALICGAMALAMWKLPGSFLPVEDQGRSWSSTPCPPGRRRCVPPRCAAR |
| 94. | Protein_ID\|573.43724.peg.3729_RND efflux system, inner membrane transporter | MTPPSVDGLGQSNGFTFELMASGGTDRDSLMKLRSQLLAAANQSSELQSVRANDLPQMPQ  LQVDIDNNKAVSLGLSLSDVTDTLSSAWGGTYVNDFIDRGRVKKVYIQGESDARAVPSDL  GKWFVRGSDNSMTPFSAFATTHWQYGPESLVRYNGSAAFEIQGENAAGFSSGAAMDKMEK  LADSLPAGSTWAWSGISLQEKLASGQAMSLYAISILVVFLCLAALYESWSVPFSVIMVIP  LGLLGAALAATLRGLSNDVYFRWRC |
| 95. | Protein_ID\|573.43724.peg.3733_Efflux transport system, outer membrane factor(OMF) lipoprotein | MLLRRPDIQEAEHNLKSANADIGAARANFFPSISLTASAGVGSDSLSSLFSHGMQVWSFA  PSISLPLFTGGSNLAQLRYAEAEKKGLIATYEKSIQSAFKDVADALARRETLSEELDAQR  QYVAAEQTSLDIAMKSYQAGVGDYLSVLTAQRTLWSAKTTLLSLQQTDLNNRITLWQSLG  GGAS |
| 96. | Protein_ID\|573.43724.peg.3861_Uncharacterized glycosyl hyrdrolase YieL | MKRHAIYFALALAGAAFTLQAAPLPAMPDPSLPVSHFITQVNADKSITYRLFAPDARRVS  IVTGATPDSFVSHDMTKAADGVWTWKSEPMKPNLYEYYFDVDGFRSVDTGSRYQKPQRQV  NTSLILVPGSILDDREVAHGDLRTLTYHSKALNAERRLYVWTPPGYSGTGDPLPVLYFYH  GFGDSGLSAIDQGRIPQIMDNLLAEGKIKPMLVVVPDTETDIPEAVAENFPPQERRKTFY  PLNAQAADKELMQDIIPLIDARFNVRKDADGRALAGLSQGGYQALVSGMNHLESFGWLAT  FSGVTTTTVPNAGVEAQLKQPDAINKQLRNFTVVVGEKDSVTGKDIAGLKSELEKQQIKF  DYHQYPGLNHEMDVWRPAYAEFVQKLFK |
| 97. | Protein_ID\|573.43724.peg.3968_Phosphate ABC transporter, substrate-binding protein PstS (TC 3.A.1.7.1) | MSLLSLVPLIAVVFALFAAFPMFSEVSVQIRHFIFANFIPATGDVIQGYIEQFVANSSRM  TAVGAFGLIVTSLLLMYSIDSALNTIWRSTRSRPKVYSFAVYWMILTLGPLLAGASLAIS  SYLLSLRWASDLDGVIDNLLRLFPLILSWAAFWLLYSIVPTTQVRNRDAVIGALVAALLF  EAGKKAFALYITTFPSYQLIYGVISVVPILFVWVYWTWCIVLLGAEITVTLGEYRKLKTE  ETEQP |
| 98. | Protein_ID\|573.43724.peg.3978_Formate dehydrogenase O beta subunit (EC 1.2.1.2) | MAKLIDVTTCIGCKACQVACSEWNDIRDEVGHNVGVYDNPADLTAKSWTVMRFSEVEQND  KLEWLIRKDGCMHCADPGCLKACPSEGAIIQYANGIVDFQSEQCIGCGYCIAGCPFDVPR  LNPEDNRVYKCTLCVDRVTVGQEPACVKTCPTGAIHFGSKEDMKTLAGERVAELKTRGYD  NAGLYDPAGVGGTHVMYVLHHADKPNLYHGLPENPEISQTVKFWKGIWKPLAAVGFAATF  AASIFHYVGVGPNRAEEEEDNLHEEKDEVRK |
| 99. | Protein_ID\|573.43724.peg.4123_Lipopolysaccharide biosynthesis protein WzzE | MERHGDHRSAYCQYVGGYYSQQQFLRNLDIKADLASVDQPSAMDEAYKEFIMQLASWDTR  RDFWLQTDYYKQRQSGNARADAAMLDDLINNIQFMPGDAAKSINDSVKLTAETGQDANNL  LRQYVAFASQRAAGHLNDELKGAWAARTVQMKAQVKRQEEVAEAIFNRRTHSVEQALKVA  QQHNISRSETDVPADQLPDSELFLLGRPMLQARLENLQAVGPEYDLDYDQNRAMLSTLNV  GPTLDPRFQTYRYLRTPEEPVKRDSPRRVFLMVMWGIVGALIGAGVALSRRRVL |
| 100. | Protein_ID\|573.43724.peg.4142_Lipopolysaccharide N-acetylmannosaminouronosyltransferase (EC 2.4.1.180) | MQRAGAEGTPVFLVGGKPEVLAQTESRLRQRWQVNIVGSQDGYFTPEQRQALFERIRDSG  AKIVTVAMGSPRQEIFMRDCRRLYPHALYMGVGGTYDVFTGHVHRAPKFWQDLGLEWFYR  LLLQPSRIKRQFRLLRYLRWHYSGKL |
| 101. | Protein_ID\|573.43724.peg.4149_Uroporphyrinogen-III synthase (EC 4.2.1.75) | MLQLPELQNIAGKNALILRGNGGRELLGATLTERGARVTFCECYQRSAKHYDGAEEAMRW  QSRGVTTLVVTSGEMLQQLWSLIPQWYREQWLLHCRVVVVSERLALQARELGWQEIQVAD  SADNDALLRALQ |
| 102. | Protein_ID\|573.43724.peg.4150_Uroporphyrinogen-III synthase (EC 4.2.1.75) | MSILVTRPLPQGEALVSRLRAMGRVAWSFPLIEFTPGRELPALGDALARLGPDDLLFALS  QHAVEFAHARLQQQGLPCPLLRAILPLAARRRWPCIRSVGSISITR |
| 103. | Protein_ID\|573.43724.peg.4179_Mobile element protein | MAETATPHDAIFKTFLSRVETARDFIELHLPPSLTQICKLDTLRLESGSFLEDDLRPYYS  DILYSLETTRGSGYVHVLIEHQSAPDKLMAFRLMRYAIAAMQRHRRAAIKRCRWLSLFSS  IRDDAAPIPGRCAGWIILPILRPPASCIAGRFRWSISR |
| 104. | Protein_ID\|573.43724.peg.4221_Protoporphyrinogen IX oxidase,oxygen-independent, HemG | MKTLILFSTRDGQTREIASFLASELKELGIDADTLNLNRTDVVEWHHYDRVVIGASIRYG  HFHPAVDRFVKKHLAALQALPGAFFSVNLVARKPEKRTPQTNSYTRKFLLNSPWQPQSCA  VFAGALRYPRYSWYDRFMIRLIMKMTGGETDTRKEVVYTDWQQVSRFAREIAQMARK |
| 105. | Protein_ID\|573.43724.peg.4258_Regulator of sigma D | MLNQLENLTERVGGSNELVDRWLQVRKHLLVAYYNLVGLKPGKESFMRLNEKALDDFCQS  LVDYLSSGHFSIYERIIGEMEGDTLF |
| 106. | Protein_ID\|573.43724.peg.4273_Response regulator of zinc sigma-54-dependent two-component system | MMQVRLLRAIQEREVQRVGSNQTLAVDVRLIAATHRNLAEEVSAGRFRQDLYYRLNVVTI  EIPPLRRRREDIPQLAQHFLKRYTERNRKTVKGFTPQAMDLLIHYPGRAIYASWKTRWSG  RWCY |
| 107. | Protein_ID\|573.43724.peg.4283_Isocitrate lyase (EC 4.1.3.1) | MQQAKAGIEAIYLSGWQVAADANLASSMYPDQSLYPANSVPAVVDRINNTFRRADQIQWS  AGIEPNDPRFIDYFLPIVADAEAGFGGVLNAFELMKSMIEAGAAAVHFEDQLASVKKCGH  MGGKVLVPTQEAIQKLVAARLAADVMGVPTLVIARTDADAADLITSDCDPYDREFITGDR  TSEGFFRTHAGIEQAISRGLAYAPYADLVWCETSKPDLEQARRFAEAIHARFPGKLLAYN  CSPSFNWKKNLDDKTIASFQQQLSDMGYKYQFITLAGIHSMWFNMFDLAHAYAQGEGMRH  YVEKVQQPEFAAGPEGYTFVSHQQEVGTGYFDKVTTIIQGGTSSVTALTGSTEEDQF |
| 108. | Protein_ID\|573.43724.peg.4367_Outer membrane protein assembly factor YaeT | MLKKTHIISGLLIAPLTLYAATSYQVDDIRFEGLQRVTVGAALLSMPLHAGDAVTPEDVS  EAVRALYASGNFENVQILRDGKTLVVQVKERPTIASVSFSGNKAVKDDALKENLTASGIS  AGSALDRNSLSEIEKGLQDFYYSAGKYSAQVHAVVTPLPRNRVDLTFVFQEGISAKIAQI  NIIGNQAFREEALLDQLQLRDNVPWWNVVADKKYQKQKLEADLETLRSFYLDRGYARFAI  ESTQVSMTPDKKSLYITIALNEGERYRVDRTQVTGDLAQHGPEIEALAQPLAGAWYSGAQ  VTTVENEIKKHFGKYGYAWPQVTSTPEIDDAHHRVVLHIQVNAGRRYSVRQIRFSGNDTS  RDAVLRREMRQMEGAWLNNEKVDQGKVRLDRTGFFENVEQQIVPVNGTADQVDVVYKVKE  RNTGSFNVGLGFGTDSGVSYQLGVTQDNWLGTGNSVSFNGTRNSYQSYLELGATNPWFTV  DGISLGGKIFYNSYDASDADAGSYNQQSYGLGSTLGFPISENNSLNLGLDYVHNRLTNMD  PELTTWRYLSSRGIEPSVVTKDGDSGAKYSANDYFVSLGWGYNDLDRGFFPRAGNKSSLS  GKVTLPGSDNSYYKLSFDTAQYLPLSENKRWVWMERLRAGYAGGLDGKSVPFYDNFYAGG  SSSVRGFTSNTIGPKAAYYRCNGSESSYSACPLDASSDAVGGNAMAVLNSEFIIPTPL |
| 109. | Protein_ID\|573.43724.peg.4385_c-di-GMP phosphodiesterase (EC 3.1.4.52)/PdeC | MWSRTRRQYHSPRNMLQRALSCRQLRLPYQPIIDIKNNRCVGAEALLRWPGFDGPVMNPA  EFIPLAENEGMIAQVTDYVVDELFYEMGEFLASHPQLYIAINLSASDFHSARLISQISEK  AHSYAVCIGQIKIEVTERGFIDVPKTTPVIQAFREAGYEIAIDDFGTGYSNLHNLHALNV  DILKIDKTFVDTLTTNNTSHLIAEHIIEMARGLRLKTIAEGVETPEQVSWLYKRGVQYCQ  GWLFAKAMPAREFMQWLANAPAPANVPQQPRHAEI |
| 110. | Protein_ID\|573.43724.peg.4401_Outer membrane usher protein fimD precursor | MIMVCQGRNIPLSFSSVELLFSTKSICVAIILCLSTLPSRAEAGEYFNPNLLEVAESPAA  SVDLSYFSQDGIPPGTYHLDVYINDKYVSSDSLTFQEISHDAGATASPCLSAEYLNSWLI  NTTAYPQLFEAGETCARLSAIPGMTFSVSLAQQRIDFTVPQAAMLNRPRDYIPESQWQQG  INAGLLNYSVTGQRNAPRHNGATIDSQFVSLQPGLNLGPWRLRNYSTYSHSDNNSRWESV  YSYLARDIHTLRSQLVVGNTYTSSGIFDSLSFTGLQLSSDKEMLPDSLHGFAPTIRGIAR  TTAEVSVYQNGYSIYKTTVAPGAFEINDLYATGSAGDLYVNIKESDGSEQNFVVPFASLA  ILQREGQLDYALSSGRTRSGSSDDKEYNFIQSSLAYGATSNITLYTGFQQAEDKYTNLLL  GAGFNLGTIGALSFDGSQSWADVKTSDTASSTSKEQGQSYRVRFSKSFLQTGTSFSVAGY  RYSTSGYYSFQDFVDNSSTQRDCCTQSGRTKGRFDASLSQTLFGYGSLSLSLVNETYWDS  SRMESVGVGYSGSIGKASYFINYSYNRNVQSSDDSDNNRPASDTVVSLTLSIPLGETLSA  NYTLNHGRHNDTTHSVGLNGSAFEDRSLNWSLMEGYNTKDKSTSGNLSVNYQGSKGDVAG  GYGYDRYSNNYNYSLRGAWSPMRADSRYHAFWARVLRWSKRLASAM |
| 111. | Protein_ID\|573.43724.peg.4501_Outer membrane usher protein fimD precursor | MPGTYTSYQLTNSNPGSTDQSVSIGGNALENDSLEWSLQQGYSNREYYSGDMRATYNGAR  GSVNAGYSYDRSSERIDYGANGSIVAHADGITLGQDITDAAVLVKAPGLDNVKLTNDNTI  STDYRGYAIVPYVTPYRRTDITLDSSTLGEDMELPETTQSVVPTRGAIVRANYAGNIGRR  ALIQLEMPGDKPVPYGSTVTIKNSANTQANIVSDEGMVYLSGLQDSGELLVLWGQRQDQR  CNATYQLLPTQGSLTTSKAICR |
| 112. | Protein_ID\|573.43724.peg.4502_Outer membrane usher protein fimD precursor | MPFATLPVMVRENQLEYEITSGKYRPYDGGVDETPFTQATATYGVSSSLTLYGGMQAASR  YQALSTGLGYNLGELGAASADVTQAWSKMKDDEKTSGQSWRVRYGKNIVETGTNVTIAGY  RYSTRGFHTLSEVLDSYSNDGYYTSRSLRNRTNLTVNQSLGKGLGSLSISGLIEDYWDDK  RTNKSISVGYNGGFRNVNYYLGYSYNRYTWVEITQGKTLRTTNVLH |
| 113. | Protein_ID\|573.43724.peg.4503_Outer membrane usher protein fimD precursor | MFNGRYLDTRTIKFVANNRASSDNREPTLVPCLSLKTLAEYGVRIKAFPELAEDKNGCAN  FSVIPDTKADFDFTAQRLNISIPQAALSTTAQGYIPPDQFDDGINALLVNYQFSGSNDMQ  ANDEYYSLNLQSGLNVGPWRIRNLSTWNKNNSGAGDWDSAYLYMQRSIRSINSNLVMGES  SSLNGIFDSVPFTGIQLATDTTMLPESMRGYAPIIRGIARTNARVTIKQNGYQVYQTYVA  PGHLKSPICTRAAVAATFTSAWKSPTDQARICCAVRHPAGHGA |
| 114. | Protein_ID\|573.43724.peg.4504_Putative fimbrial chaparone | MFRTLLNLTASCALLLSSIAHAGIVVESTRYLYKEGAREITAQIENKDDIPYLIKSWVET  PAGKAPSFMATPPLFRLEGKQQNTVRLFATGNVNAPTDRESMYYFNVMAIPPADDAKANN  NTIQLAVRHRMRLVYRPKTLFDLSPNTEAKKLEWRKAGKKLTIKNPTPFFFYFNSIQIGG  KEVKPEVNSVAPMTTKEVTLKENINASSITWKVVNDYGGAGSLYSSSL |
| 115. | Protein_ID\|573.43724.peg.4575_FKBP-type peptidyl-prolyl cis-trans isomerase FklB (EC 5.2.1.8) | MLSHAFILMFITQKGTTMATPTFDTIEAQASYGIGLQVGQQLSESGLQGLLPEALVAGIA  DALEGNQPQVPVEAVHRALREIHERADAVRRERFQAMAADGQKYLDENREKEGVNSTESG  LQFRVLTQGEGPIPARTDRVRVHYTGATDRRHRLRQLRRSR |
| 116. | Protein_ID\|573.43724.peg.4586_2',3'-cyclic-nucleotide 2'-phosphodiesterase(EC 3.1.4.16) / 3'-nucleotidase (EC 3.1.3.6) | MAAKELKEGDVHPVYKAMNTLNYAVGNLGNHEFNYGLDFLHKALAGAKFPYVNANIIDAK  TGKPMFTPYLIQDTRVVDSDGQIHTLRIGYIGFVPPQIMTWDKANLNGKVTVNDITETAR  KYIPEMRAKGADVVVVVAHSGLSADPYQAMAENSVYYLSQVPGVDAIMFGHAHAVFPGKD  FANIKGADIAKGTLNGVPAVMPGMWGDHLGVVDLVLNNDSGKWQVTQSKAEARPIYDAVA  KKSLAAEDAKLVAVLKADHDATREFVSKPIGKSADNMYSYLALVQDDPTVQVVNMAQKAY  VEHYIQGDPDLAKLPVLSAAAPFKVGGRKNDPASFVEVEKGQLTFRNAADLYLYPNTLVV  MKVSGKEVKEWLECSAGQFNQIDPASSKPQSLINWDGFRTYNFDVIDGVNYQIDVTQPAR  YDGECQMIHPQAERIKHLTFNGKPVDPQATFLVATNNYRAYGGKFAGTGESHIAFASPDE  NRSVLAAWIGAQSKKEGAIHPAADNNWRLAPIHSNTPLDIRFETSPGDKAAAFIKEKAQY  PMRQVATDDIGFAIYQLDLSK |
| 117. | Protein_ID\|573.43724.peg.4602_Outer membrane component of TAM transport system | MGLGHQLHRDAGQDVWIRTLYDRHRFVVRGNLGWIEADNFDKVPPDLRFFAGGDRSIRGY  KYKSISPKDDDGKLIGASKLATGSLEYQYNVSGKWWGAVFIDSGEAVSDIRESDFKTGAG  VGVRWQSPVGPIKLDIARPIGDKEEHGLQFYIGLGPEL |
| 118. | Protein_ID\|573.43724.peg.4630_UPF0307 protein YjgA | MLRNRDVDPIRQALDKLKNRHNQQVALFHKLEQIRDRLIDDGDDAVAEVLNLWPDADRQQ  LRSLIRNAKKEKEGNKPPKSARLIFQYLRELAENEG |
| 119. | Protein_ID\|573.43724.peg.4704_Glycerol-3-phosphate-binding protein | MFNPLTALTVGLSLALSGAALAKEKIDFMFPAPVDGKLTMEMTRVIKTFNDSQQDVEVRG  IFTGNYDTTKIKAESAQKAGQPPALVIMSANFTTDLALKDEILPMDELFKYGDQKAGDFL  QKEFWPAMHKNAQVMGTTYAIPFHNSTPILYYNKTLFDRAGIAQPPQTWAELLSDAKKLT  DESKGQWGIMLPSTNDDYGGWIFSALVRANGGKYFNEDYPGEVYYNSPTAIGALRFWQDL  IYKDKVMPSGVLNSKQISASFFSGKLGMAMLSTGALGFMRENSKDFELGVAMLPAKEQRA  VPIGGASLVSFKGINDAQKKAAYQFLTYLVSPQVNGAWSRFTGYFSPRKASYDTPEMKAY  LQQDPRAAIALEQLKYAHPWYSTWETVAVRKAMENQLAAVVNDAKVTPEAAVQAAQKEAD  ALMKPYVDKTALAEVK |
| 120. | Protein_ID\|573.43724.peg.4713_Carbonic anhydrase, beta class (EC 4.2.1.1) | MTTLKPLLARNRSWALQKCQHDPDYFEKWVDGQRPHSLWIGCSDSRVPAEVLTGSQPGEL  FVHRNIANMLDPADDNVMSVLQYALHYLEVERVVLCGHYGCGGVQAALSLPTLPLAQESS  ALARRIGQLRNTLHHEIVQIADGCGVAASPAPAPTRSPLAMRWTPWWKPTFAPSSPVCWR  VNRCRRCSPAGDRSACTAASTIWLPAI |
| 121. | Protein_ID\|573.43724.peg.4733_Mobile element protein | MSLRQACRILSLSRTVFRYQPDTRRDEPVIMALTVAAERYPRYGYKRFFQVLRRQGNAWN  HKRVHRIYFLLKLNFRRKGKQHLPVRNPVPLVTPEAMNQSWSIDFMHDALVCGRRFRTFN  VVDDFNREALAIEIDLNIPAQRVVRVLDRIVANRGYPLKMRMDNGPELVLLTLAQWAEEH  GVMLEFLRPGKPTQNAFIE |
| 122. | Protein_ID\|573.43724.peg.4778_Copper/silver efflux RND transporter,transmembrane protein CusA | MIEWIIRRSVANRFLVMMAALFLSICGTWTIVHTPVDALPDLSDVQVIVKTRYPGQAPQI  VENQVTWPLTTTMLSVPGAKTVRGFSQFGDSYVYVIFEDGTDPYWARSRVLEYLNQVQGK  LPAGVSAEMGPDATGVGWVFEYALVDRSGKHDLAELRSLQDWFLKYELKTIPNVSEVASV  GGVVKEYQIVVDPMKLTQYGISLGEVKSALDASNQEAGGSSVELAEAEYMVRASGYLQTL  DDFKNIVLKTGDNGVPVYLGDVARVQIGPEMRRGIAELNGEGEVAGGVVILRSGKNAREV  ISAVKAKLASLQSSLPEGVEVVTTYDRSQLIDRAIDNLSYKLLEEFIVVALVCALFLWHV  RSALVAIISLPLGLCFAFIMMHFQGSTPTSCRWAGSPLRWERWSMPPS |
| 123. | Protein_ID\|573.43724.peg.4783_Copper/silver efflux RND transporter, outer membrane protein CusC | MKSLISTALTNNRDLRMATLKVQEARAQYRVTDADRYPQLNGDGSTTYGGKLKGDTTTSS  DYAAGLNLSYDLDFFGRLKNLSEADRQNFFASEEARRAVHILLIANVSQSYFNQRLAAAQ  LQVANDTLQNYQQSYAFVEKQLLTGSTTVLALEQARGMIESTRADIAKRQGQLAQANNAL  QLLLGSYQHLPDDSASSAVDLQGVTLPPSLSSTILLQRPDILEAEHSLQAANANIGAARA  AFFPSITLTSSLSGSSSELSSLFNAGGAMWNFIPKIELPIFNAGRNQASLDLAEIRQQQQ  VVNYEQKSSPPLKRWPTRWPCARAWPIKLPRRSAISPR |
| 124. | Protein_ID\|573.43724.peg.4954_Soluble lytic murein transglycosylase (EC 4.2.2.n1) | MEKAKKMTWHLLAASVGLLTLSQLAHADSLDEQRSRYAQIKQAWDNRQMDVVDQLMPTLS  TYPLYPYLQYRQITDDLMNQPALVVKNFIDANPTLPPARSLRSRFVNELARRSDWRGLLA  FSPDKPTSTEAQCNYYYAKLSVGQSQEAWSGAKELWLTGKNQPGACEPLFSAWRDSGQQD  PLAYLERIRLAMKAGNIGLVKSLAQQMPANYQSIASAVVALANDPNSVLTFARTTGATDF  TRQMAAVAFASVARQDVENARLMIPSLVQAQQLNEDQTQELRDIVAWRLMGSDVTEEQAI  WRDDAIMRSQSTPLVERRVRIALGTGDRHGLNTWLARLPMEAKEKDEWRYWQADLLLERG  RDEEAQAILRSLMQQRGFYPMVAAQRLGEEYTFRIDKASGTIDPALASGPEMARVRELMY  WNMDNTARTEWANLVTSRTKSQQAQLARYAFDQHWWDLSVQATIAGKLWDQLEERFPLAY  NDLFARYVSGKDIPQSYAMAIARQESAWNPKVRSPVGASGLMQIMPGTATHTVSMFSIPG  YSGPSQLLDPETNINIGTSYLQYVYQQFGNNRIYASAAYNAGPGRVRSWQGNSAGRIDAV  AFVESIPFSETRGYVKNVLSYDAYYRYFMGQQDKILSDAEWRQRY |
| 125. | Protein_ID\|573.43724.peg.5024_4-hydroxythreonine-4-phosphate dehydrogenase(EC 1.1.1.262) | MGTEEIDTIIPVLEEMRAKGMNLSGPLPADTLFQPKYLDHADAVLAMYHDQGLPVLKYQG FGRGVNITLGLPFIRTSVDHGTALELAGQGKADVGSFITALNLAIKMIVNTQ |
| 126. | Protein_ID\|573.43724.peg.5025_4-hydroxythreonine-4-phosphate dehydrogenase(EC 1.1.1.262) | MPETRKVVITPGEPAGIGPDLVVQLAQRDWPVELVICADGALLTDRAKRLGLPLSLLPYD  PAQPPVPQRAGTLTLLNVALNVPAEPGVLNVQNGAYVVETLARACDGCLSGEFAALVTGP  VHKGNINDAGIAFTGHTEFFEERAKASKVVMMLATEELRVALVTTHLPLKAISEAITPSY  CARSSPFSITTCARNLALPSRMCWFAGLTRTLAKAAIWEQKR |
| 127. | Protein_ID\|573.43724.peg.5056_Thiamin ABC transporter, transmembrane component | MATRRQPLNLRGLMPGLFAATLLCAVALAAFLALWFSAPGAGWQSVFSDSYLWHVVRFSF  WQASLSALISVGPAIFLARALYRRRFPGRTLLLRLCAMTLILPVLVAVFGILSVYGRQGW  LASLFHALGWQWEFSPYGLQGILLAHVFFNMPMATRLLLQALENIPGEQRQIAAQLGMRG  YAFFRLVEWPWLRRHIPAVAALIFMLCFASFATVLSLGGGPKATTIELAIYQALSFDYDP  ARAAMLALIQMLCCLGLVLLSQHLSKAVAIGVSHVRGWRDPDDRLHSRLSDGLLIGAALL  LLLPPLLAVIVDGINRNMLDVLAQPALWQALSTSLRIAIAAGLLSVILTMMLLWSSRELR  ARQRSLAGQAMELSGMLILAMPGIVLATGFFLLLNNTIGLPESADGIVIFTNALMAIPYA  LKVLENPMRDIAARYSMLCQSLGIEGFARLRVVELRALRRPLARRSPSPAYCRLAISAWW  RCSATRRSAPCLSICTSR |
| 128. | Protein_ID\|573.43724.peg.5079_3-isopropylmalate dehydratase small subunit (EC 4.2.1.33) | MTLSDEQVDELFKLVQANPGITFEVDLEAQVVKAGDKTYSFKIDDFRRHCMLNGLDSIGL  TLQHEAAISDYERKLPVFMN |
| 129. | Protein_ID\|573.43724.peg.5080_3-isopropylmalate dehydratase small subunit (EC 4.2.1.33) | MAEKFTQHTGLVVPLDAANVDTDAIIPKQFLQKVTRTGFGAHLFNDWRFLDDKGQQPNPE  FVLNFPEYQGIDTAGAGKLRLRLLARARAVGADRLRL |
| 130. | Protein_ID\|573.43724.peg.5117_UDP-3-O-[3-hydroxymyristoyl]N-acetylglucosamine deacetylase (EC 3.5.1.108) | MIKQRTLKRIVQATGVGLHTGKKVTLTLRPAPANTGVIYRRTDLNPPVDFPADAKSVRDT  MLCTCLVNEHDVRISTVEHLNAALAGLGIDNIIVEVDAPEIPIMDGSAAPFVYLLLDAGI  DELNCAKKFVRIKETVRVEDGDKWAEFKPYNGFSLDFTIDFNHPAIDASTQRYTLNFSAD  AFMRQISRARTFGFMRDIEYLQSRGLCLGGSFDCAIVVDDYRVLNEDGLRFEDEFVRHKM  LDAIGDLFMCGHNIIGAFTAYKSGHALNNKLLQAVLAKQEAWEYVTFEDDAKLPMAFRAP  SMVLA |
| 131. | Protein_ID\|573.43724.peg.5177_Carbonic anhydrase, beta class (EC 4.2.1.1) | MNDIDTLISNNALWSKMLVEEDPGFFEKLSQTQKPRFLWIGCSDSRVPAERLTGLEPGEL  FVHRNVANLVIHTDLNCLSVVQYAVDVLEVEHIIICGHYGCGGVQAAVENPELGLIDNWL  LHIRDIWFKHSSLLGEMPEERRLDTLCELNVMEQVYNLGHSTIMQSAWKRGQKVTIHGWA  YGIHDGLLRDLDVTAVSRETLEQRYRHGSPTSRSSTSTTDSLQTSPPWRARRALSRVGYP  LTRPAG |
| 132. | Protein_ID\|573.43724.peg.5185_Pantoate--beta-alanine ligase (EC 6.3.2.1) | MLEGASRPGHFRGVSTIVSKLFNLVQPDVACFGEKDFQQLALIRKMVADMGYDIEIIGVP  IVRAKDGLALSSRNGYLTADQRKIAPGLYKVLSAVAEKLAAGDRQLDEIIAIAEQELNEK  GFRADDIQIRDADTLLELTDASQRAVILMAAWLGQARLIDNRIVTLAQ |
| 133. | Protein_ID\|573.43724.peg.5265_Outer membrane protein assembly factor YaeT | MAMKKLLIASLLFSSATVYGAEGFVVKDIHFEGLQRVAVGAALLSMPVRPGDTVTDDDIS  NTIRALFATGNFEDVRVLRDGDTLLVQVKERPTIASITFSGNKSVKDDMLKQNLEASGVR  VGESLDRTTIADIEKGLEDFYYSVGKYSASVKAVVTPLPRNRVDLKLVFRKASPQKFNRS  TSSATMRFRPMS |
| 134. | Protein_ID\|573.43724.peg.5267_Outer membrane protein assembly factor YaeT | MTGNLAGHSAEIEALTKVEPGELYNGAKVTKMENDIKKLLGRYGYAYPRVQSQPEINDSD  KTVKLHVNVDAGNRYYVRKIRFEGNDTSKDAVLRREMRQMEGAWLGSDLVDQGKDRLNRL  GFFETVDTDTQRVPGSPDQVDVVYKVKERNTGSFNFGIGYGTESGVSFQAGVQQDNWLGT  GYAVGINGTKNDYQTYTELSVTNPYFTVDGVSLGGRVFYNDFDANDADLSDYTNKSYGTD  ITLGFPVNEYNTLRAGVGYVHNSLSNMQPQVAMWRYLNSMGQYPDNTNDRNSFSANDFTF  NYGWTYNKLDRGFFPTEGSRVNLNGKVTIPGSDNEYYKATLDTATYVPIDNDHQWVVLGR  TRFGYGDGIGGKEMPFYENFYAGGSSTVRGFQSNTIGPKAVYFPASSRHDDDDSYDNECK  STESAPCKSDDAVGGNAMAVASLELITPTPFISDKYANSVRTSVFWDMGTVWDTHWDSSA  YAGYPDYSDPSNIRMSAGIAVQWMSPLGPLVFSYAQPFKKYDGDKAEQFQFNIGKTW |
| 135. | Protein_ID\|573.43724.peg.5272_Acyl-[acyl-carrier-protein]--UDP-N-acetylglucosamine O-acyltransferase (EC 2.3.1.129) | MIDKTAFVHPTAIVEEGAVIGANVHIGPFCIVGANVEIGEGTVLKSHVVVNGHTKIGRDN  EIYQFASIGEVNQDLKYAGEPTRVEIGDRNRIRESVTIHRGTVQGGGLTKVGNDNLLMIN  AHVAHDCTLGDRCILANNATLAGHVSLDDYVIIGGMTAVHQFCVIGSHVMVGGCSGVAQD  VPPFVIAQGNHATPFGVNIEGLKRRGFSREAITAIRNAYKLLYRSGKTLEEAKPEIAELAAQHPEVQPFVDFFARSTRGLIR |
| 136. | Protein_ID\|573.43724.peg.5273_Lipid-A-disaccharide synthase (EC 2.4.1.182) | MAEQRPLTIALVAGETSGDILGAGLIRALKARIPNARFVGVAGPLMQAEGCEAWYEMEEL  AVMGIVEVLGRLRRLLHIRADLTRRFGELRPDVFVGIDAPDFNITLEGNLKKQGIKTIHY  VSPSVWAWRQKRVFKIGRSTDLVLAFLPFEKAFYDKFNVPCRFIGHTMADAMPLDPDKGA  ARDRLGIPHSVRCLALLPGSRGAEVEMLSADFLKTAQLLRATYPDLQVVVPLVNAKRREQ  FERIKAETAPGMIVHMLDGQARDAMIASDAALLASGTAALECMLAKCPMVVGYRMKPFTF  WLAKRLVKTDYVSLPNLLAGRELVKELLQDECEPQALAAALQPLLADGKTSHEMHETFRA  LHQQIRCNADEQAADAVLELAKQ |
| 137. | Protein_ID\|573.43724.peg.5301_D-glycero-beta-D-manno-heptose-1,7-bisphosphate 7-phosphatase (EC 3.1.3.82) | MAKSVPAIFLDRDGTINVDHGYVHEIDNFEFIDGVIDAMRELKEMGYALVLVTNQSGIAR  GKFTEAQFETLTEWMDWSLADRGVDLDGIYYCPHHPQGAVEEYRQTCDCRKPHPGMLISA  RDYLHIDMAASYMVGDKLEDMQAAAAADVGTKVLVRTGKPLTEEAEKAADWVLNSLAELP  AAIKKQQK |
| 138. | Protein_ID\|573.43724.peg.5323_D-sedoheptulose 7-phosphate isomerase (EC 5.3.1.28) | MYQDLIRNELNEAAETLANFLQDEANIHAIQRAAVLLADSFKAGGKVLSCGNGGSHCDAM  HFAEELTGRYRENRPGYPAIAISDVSHLSCVSNDFGYEYVFSRYVESVGRAGDVLLGIST  SGNSGNVIKAIEAARAQGMKVITLTGKDGGKMAGSADVEIRVPHFGYADRIQEIHIKVIH  ILIMLIEKEMAKG |
| 139. | Protein_ID\|573.43724.peg.5396_Outer membrane porin PhoE | MKKSTLALMMMGFVASTATQAAEVYNKNANKLDVYGKIKAMHYFSDYDSKDGDQTYVRFG  IKGETQINDDLTGYGRWESEFSGNKTESDSSQKTRLAFAGVKLKNYGSFDYGRNLGALYD  VEAWTDMFPEFGGDSSAQTDNFMTKRASGLATYRNTDFFGLVDGLDLTLQYQGKNEGREA  KKQNGDGVGTSLSYDFGGSDFAVSAAYTSSDRTNDQNLLARGRGSKAEAWATGLKYDANN  IYLATMYSETRKMTPISGGFANKAQNFEAVAQYQFDFGLRPSLGYVLSKGKDIEGVGSED  LVNYIDVGLTYYFNKNMNAFVDYKINQLKSDNKLGINDDDIVALGMTYQF |
| 140. | Protein_ID\|573.43724.peg.5422_CFA/I fimbrial chaperone | MSRRRGATLTKALLTVGCLLAAPLAQAISVGNLTFSLPAEADFASKRVVNNNKSARLYRI  AVSAIDRPGGSEVRSRPVDGELLFAPASWCCRQVRASILNFTIMGRGITASATIGSRFAK  SPPAT |
| 141. | Protein_ID\|573.43724.peg.5423_CFA/I fimbrial chaperone | MEPVVVMDTILVVRPREVQFKWSFDKVAGTVSNTGNTWFKLLIKPGCDSTEEEGDAWYLR  PGDVVRQPALRQPGNHYLVYNDKFIKISDTCPLKPRPAE |
| 142. | Protein_ID\|573.43724.peg.5598_Peptidyl-prolyl cis-trans isomerase PpiD (EC 5.2.1.8) | MAEIGRGQFENAVASERNRMQQQLGDQFSELAANENYMKTMRQQVLNRLIDESLLDQYAR  ELGLSISDEQVKQAIFQTQAFQTNGKFDNQRFSGIVAQMGMTTDQYAQALRNQLTTQQLI  NAIAGTDFMLPGESDQLAALVSQQRVVREATINVNALAAKQTASDEEINAFWQQNQARFM  APEQFRVSYIKMDAASMQESASDEEIQSWYDQHKDQFTQPQRNRYSVIQTKTEADAKAVL  AELQKGADFATLAKEKSTDIISARNGGDMGWMEDASTVPELKDAGLKEKGQLSGVIKSSV  GFLVARLDDVQPAQVKPLADVRNDIAAKVKQEKALDAYYALQQKVSDAASNDNESLASAA  QVAGMKVVETGWFGRDNLPEELNFKPVADAIFNGGLVGENGAPGSNSDIITVDGDRAFVL  RISEHKAEAVKPLAEVKAQVSDIVKHNKAEQQAKLEADKLLAELKDGKGDEAMKAAGLSF  GAPQTLSRTGQDPLSQLAFTLPLPQQGKPVYGVGSNMQGDVVLVALDEVKPAACRKSRRR  PWFRGSPRTMPKSLSKR |
| 143. | Protein_ID\|573.43724.peg.5626_c-di-GMP phosphodiesterase (EC 3.1.4.52)/PdeB | MTAMGSRQHMVMIDPVSFIDVVPASEEKIHTMLFGLDHQKLVISSQPLPAKVWQRIKDPH  VDMLTLDNTVYRIQRIPELGSGIVTWSSTLPLQQRIRQQLFFWLPAGIFTSLLATWLLLR  LLRHLRSPRNSMLDALNSEAIQVHYQPIISLQDGKIAGAEALARWQQPDGTFLSPDIFIP  LAEQTGLITQLTEDIVRKIFTDLGPWLRQRPEVHISINLSVDDLRSPTLPTLLHDQLQHW  GIAAEQIILEITERGFVDPETTMPVIAHYRQAGHRISIDDFGTGYSSLSYLQKLDVDTLK  IDKSFVDTLEYRPLTPHIIEMAKALNLATVAEGVETESQRDWLRQHGVQYAQGGFTAKRC  QKSSSFCGQSITSMRIKRRVA |
| 144. | Protein_ID\|573.43724.peg.5662_Multidrug efflux system AcrAB-TolC,inner-membrane proton/drug antiporter AcrB (RND type) | MKIVYPYDTTPFVKISIHEVVKTLVEAIILVFLVMYLFLQNFRATLIPTIAVPVVLLGTF  AVLAAFGFSINTLTMFGMVLAIGLLVDDAIVVVENVERVMAEEGLPPKEATRKSMGQIQG  ALVGIAMVLSAVFIPMAFFGGSTGAIYRQFSITIVSAMALSVLVALILTPALCATMLKPI  QKGSHGATTGFFGWFNRMFDKSTHHYTDSVGNILRSTGRYLVLYLIIVVGMAWLFVRLPS  SFLPDEDQGVFLSMAQLPAGATQERTQKVLDEMTNYYLTKEKDNVESVFAVNGFGFAGRG  QNTGIAFVSLKDWSQRPGEENKVEAITARAMGYFSQIKDAMVFAFNLPAIVELGTATGFD  FELIDQGGLGHEKLTQARNQLFGMVAQHPDVLTGVRPNGLEDTPQFKIDIDQEKAQALGV  SISDINTTLGAAWGKLCQRLYRPRPREESVHHVRSEIPYAAGRYRQVVCSRQRWSDGAVL  CFLDLALGIRFAASGTLQRSAVTGNPRSGRAQARVPVRRWR |
| 145. | Protein_ID\|573.43724.peg.5664_Multidrug efflux system AcrAB-TolC,inner-membrane proton/drug antiporter AcrB (RND type) | MPNFFIDRPIFAWVIAIIIMLAGGLSILKLPVAQYPTIAPPAISITAMYPGADAETVQNT  VTQVIEQNMNGIDHLMYMSSNGDSTGTATITLTFESGTDPDIAQVQVQNKLALATPLLPQ  EVQQQGISVEKASSSFLMVVGVINTNGTMNQDDISDYVAANMKDPISRTSGVGDVQLFGS  QYAMRIWMDPNKLNNFQLTPVDVISALKAQNAQVAAGQLGGTPPVKGQQLNASIIAQTRL  TNTEEFGNILLKVNQDGSRFVCVMSPKLSWAAKAMT |
| 146. | Protein_ID\|573.43724.peg.5665_Multidrug efflux system AcrAB-TolC, membrane fusion component AcrA | MDQLTVKRYQKLLGTKYISQQDYDTAVATAQQSNAAVVAAKAAVETARINLAYTKVTSPI  SGRIGKSAVTEGALVQNGQTTALATVQQLDPIYVDVTQSSNDFLRLKQELADGRLKQENG  KAKVELVTNDGLKYPQSGTLEFSDVTVDQTTGSITLRAIFPNPDHTLLPGMFVRARLEEG  INPDALLVPQQGVTRTPRGDASVMVVGEGDKVEVRQVTASQAIGDKWLVTDGLKSGDRVI  VTGLQKSNQVCR |
| 147. | Protein_ID\|573.43724.peg.5666_Multidrug efflux system AcrAB-TolC, membrane fusion component AcrA | MNKNRGLTPLAVVLMLSGSLALTGCDDKPAQQGAQHMPEVGIVTLKSAPLQITTELPGRT  SAYRIAEVRPQVSGIILKRNFVEGSDIQAGVSLYQIDPATYQASYDSAKGDLAKARRRQT  WIN |
| 148. | Protein_ID\|573.43724.peg.5677_Inner membrane protein YbaN | MPPVILTIIGWLAVALGTLGVFLPLLPTTPFILLAAWCFARSSPRFHQWLLYRSWFGGYL  RHWQQYRAMPRGAKPRAIAMIVVTLLFRYGWLSLRGCASCCWSFWPVC |
| 149. | Protein_ID\|573.43724.peg.5727_UDP-2,3-diacylglucosamine diphosphatase (EC 3.6.1.54) | MPCYFIHGNRDFLVGQRFARQSGMILLAEEERLDLYGREVLIMHGDTLCTDDQGYLAFRA  KVHTPWIQRLFLALPLFIRRRIAARMRADSKAANSSKSMEIMDVNPQAVVDAMERHHVQW  LIHGHTHRPAVHELQANGQPAWRVVLGAWHSEGSMVKVTPDDVKLIHFPF |
| 150. | Protein_ID\|573.43724.peg.5739_Lipid A biosynthesis lauroyl acyltransferase(EC 2.3.1.241) | MRSNKTMLDRKNLKGMVRALKEGEILWYAPDHDYGPASSVFAPLFAVEQAATTTGTWMLA  KMSGATIVPFVPRRKPNGMGYELISLTPERTPPLASAEVTAAWMNQIIEQCILMAPEQYM  WLHRRFKTRPEGVPPRY |
| 151. | Protein_ID\|573.43724.peg.5825_Putative periplasmic protein | MLNNTLYVGAVADPAQIWVVDATTLKLKTRIKNTGKWMTGLHYSAQTGRVYAANGSGEIL  VINPRNRRIEQRWKPLGDKPALLLNMAEDSDTGRLFVTDNSKAKTTLVLDIHNGKLLKQL  DVGDSLAVQFNKKRHEIYISQRESGKVISLDASRYTLKKSWALPANPNSLLLSADGQTLF  VTVKQPFNKDHSTKGPDSVVRIDLNAQ |
| 152. | Protein_ID\|573.43724.peg.5826_Ferrichrome-iron receptor | MGQIMHTTHYSSFPLRKTLLALAIGAASQTAMAANAAAAKQPGEETLIVEANETSDFKSG  GDLVVPAFLDGQIAHGGRLGMLGEQKAMDVPFNVIGYTSKLIQDQQAKTIADVVSNDAGV  QAVQGYGNFAETYRIRGFKLDGDDMTMGGLAGVVPRQVMDTQMLERVEIFKGANSLLNGA  ASSGVGGMINLEPKRAEDLPTARVGVDYTSDSQVGGTLDLGRRFGDNNQFGARVNLVHRE  GEGAIDNDKRRTTLASLGLDYRGDRFRSSLDFGYQKKTFHGGTMGVNISGVDFVPALPDN  SKNYSQKWGYSDIESEFGMAKAEYDLTDSWTVYSALGGQHSHEIGTYSAPKLLNKNGDAT  VGRLDTNRIIDAISGMGGVRGDFNTGAISHKVNLGYAAQVHTDATAWRMSARNPTTNIYD  NHDVAMPDNAYFGGNYHDPLVTSRSRTQGWLLSDTLGFFNDKVLFTAAARHQKVVVRNYS  NATGLEDTSSRYTQSRWMPTFGLVYKPWEQLSLYANHTEALQPGSVAPTTAANAGQSTGI  AHSKQDEVGVKIDYGTIGGSLALFEIKKPNAISDTAGNYGLDGEQRNRGVEMNVFGEPML  GLRLNASTVWLDAKQTKTAEGATDGKDAIGVANFYAVLGAEYDIKPVEGLTATARVNHSG  SQYADAANTKKLDSYTTLDLGLRYRMRLNADQNEMIWRVGVTNVTNEKYWSGIDDTGTYL  FEGDPRTVRVSMSYEF |
| 153. | Protein_ID\|573.43724.peg.5857_Phenylalanine-specific permease | MIKTFRFALLACGLVGGALTATAAPAVPVRVATVELAPHAEERAIPGRVEAIRAVDIRAR  TEGVIVQRHFQDGQYVTEGDLLFTLDDAQPRAALALAQAELKSAEASLRQSQQLLTRYER  LINNHSISRNDVDTARMQRDVAAAAVQQAKARVEAQRIVLSYTQIAAPVTGRVGHSAFHV  GTLVNSSSGVLVDIVQLDPVRVSFALDEAAFFSKTGQHADIHALKQAWLAQIEIDGKRRD  GVLTSIDNRIDARTGSVAVRAEFANPQHRLLPGGSVTILFRPQELPSRVMVPAAAVQQDP  QGFFSWVLKPDHTAGQRRLTLAGQQGQQFAVEKGLQAGEQVITDGAQRLREGAAVQVLN |
| 154. | Protein_ID\|573.43724.peg.5859_RND efflux system, inner membrane transporter | MRGRTALSITFAAGTDADLAAIDVQNRVAQALAQLPAEVQQNGVQVRKRASNLLMGVSLY  SPLGTLSPLFVSNYASTQVREALARLPGVGEVQMFGARDYSMRIWLRPDRMNALNITTDD  VARALREQNVQGAAGQVGTPPVFNGQQQTLTINGLGRLNEAASFGEIILRRGAQGQLVRL  ADVATIELGARSYSSGAQLNGKASAYLGIYPTPTANALQVASAVRAELNRLHTRFPADLT  WEVKFDTTRFVAATIKEIGVSLALTLLAVVVVVSLFLQSWRATLIVVLAIPVSLIGTFAV  LYLLGYSANTLSLFAIILALTMVVDDAIVVVENVETKMAEGLDRLQATAQALRQIAGPVI  ATTLVLLAVFVPVALLPGIVGELYRQFAVTLSTAVALSSLVALTLTPALCALLLRAASRP  AGGGVAGV |
| 155. | Protein_ID\|573.43724.peg.5861_RND efflux system, inner membrane transporter | MTREILQLANQHPQLSRVFTTWSSNVPQLTLTVDRDRAALLDVPVAQIFSSLQTAFGGTR  AGDFSRNNRVYHVVMQNEMQWRERAEQISELYVRSRDGERVRLSNLVTITPTVGAPFIQQ  YNQFPSVSVSGSAAEGVSSRTAMAAMEQILQAHLPPGYDYAWSGISWQEQQTGNQAVWIV  LAAVAMAWLFLVAQYESWTLPASVMLSVLFAIGGALLWLWTAGYANDVYVQIGLVLLIAL  AAKNAILIVEFARSRREEGLSIVDAARARGYPPLPRGNDDRGVVYYRHYADDARHRRRGA  EPADHRHHGIQRDAGGDDG |
| 156. | Protein_ID\|573.43724.peg.5922_Uncharacterized protein STM0479 | MAQATGGAGLTHDAPFSEALRKISSKNPARRQAVVASRHSALPGKEAMKKRPHSTPHDSL  FKRFLRDSETARDFLDIHLPPALRQLCDLSTLHLESGSFVEDNLRASYSDILYSLHTSKG  EGYIYVVIEHQSTPDSHMAFRLLRYALAAMQQHLDRGHKALPLVIPMLFYHGSISPYPFS  LCWLDEFPPDSPAKQLYLGAFPLVDVTDIPDDAILQHRRIALLELVQKHIRQRDLSHILQ  QLVEVVLMGYTDHQQFKTLFTYMPLHGNSADPENFIDQLVERLPQYEDTLMTIAEYLKQK  GREEGKQRWLQEGLNEGLQAGEHREACRIALLMLENGMDTETVLRMTRLSADELATLARQ  ARQSSPPLNP |
| 157. | Protein_ID\|573.43724.peg.5931_4'-phosphopantetheinyl transferase (EC 2.7.8.-)[enterobactin] siderophore | MRHHRTALPLAGYTIQQIDFDPATFQPEDLFWLPYHASLTGWGRKRQAEHLAGRIAAAYA  LREVGDKQLPAIGDQRQPLWPTPWFGSISHCGQRALAVVADRPVGVDIERRFTPSWRRNW  RAALSVRQKKQRCCAAVCPSRWR |
| 158. | Protein_ID\|573.43724.peg.5951_Ferric enterobactin-binding periplasmic protein FepB (TC 3.A.1.14.2) | MNFFSFCRRGALTGMLLLLGITSAQAADWPRQVTDSYGTHTLPSQPLRIVSTSVTLTGSL  LAIDAPVVASGATTPNNRVADSQGFLRQWSEVAKARKLARLYIGEPSAEAVAAQMPDLIL  VSATGGDSALPLYDQLKTIAPTLVINYDDKSWQTLLTQLGQITGHEQQASARIADFNKQL  VSLKEKMKLPPQPVTALVYTAAAHSANIWTPESAQGQMLEQLGFSLATLPGGLPASHSQG  KRHDIVQLGGENLAAGLNGQSLFLFAGDQKDADAIYANPLLAHLPAVAGKRVYPLGTETF  RLDYYSALLVLQRLSSLFG |
| 159. | Protein_ID\|573.43724.peg.6069_Magnesium and cobalt efflux protein CorC | MDTIGGLVMQAFGHLPARGESIDIDGYQFKVAMADSRRIIQVHVKLPDDAPQPKLEE |
| 160. | Protein_ID\|573.43724.peg.6148_Tol-Pal system protein TolQ | MRISMNRELETLETHIPFLGTVGSISPYIGLFGTVWGIMHAFIALGAVKQATLQMVAPGI AEALIATAIGLFAAIPAVMAYNRLNQRVNKLELNYDNFMEEFTAILHRQAFTTSESNKG |
| 161. | Protein_ID\|573.43724.peg.6154_Tol-Pal system peptidoglycan-associated lipoprotein PAL | MQLNKVLKGLMIALPVMAIAACSSNKNASNDQSGEGMLGAGTGMDANGNGGNMSSEEQAR  LQMQQLQQNNIVYFDLDKYDIRSDFAAMLDAHANFLRSNPSYKVTVEGHADERGTPEYNI  ALGERRANAVKMYLQGKGVSADQISIVSYGKEKPAVLGHDEAAYAKNRRAVLVY |
| 162. | Protein_ID\|573.43724.peg.6166_2-keto-3-deoxy-D-arabino-heptulosonate-7-phosphate synthase I alpha (EC 2.5.1.54) | MNKYGNAGKKMNYQNDDLRIKEINELLPPVALLEKFPATENAANTVAHARKAIHQILKGD  DDRLLVVIGPCSIHDPAAAKEYAARLLTLREALKGELEIVMRVYFEKPRTTVGWKGLIND  PHMDNSFRINDGLRIARKLLLDINDSGLPAAGEFLDMITPQYVADLMSWGAIGARTTESQ  VHRELASGLSCPVGFKNGTDGTIKVAIDAINAAGAPHCFLSVTKWGHSAIVNTSGNGDCH  IILRGGKEPNYSAKHVAEVKIGLAKAGLPAQVMIDFSHANSSKQFKKQMEVGADVCQQIA  GGERAIMGVMIESHLVEGNQSLESGEPLTYGKSVTDACIGWEDTETILRQLAEAVKTRRG |
| 163. | Protein_ID\|573.43724.peg.6257_Endonuclease/exonuclease/phosphatase family protein | MAEPTAGFSLNVLTINTHKGFTAFNRRFILPELRDAVRSVSADIVCLQEVMGAHEVHPLH  IENWPDTTHYEFLADTMWSDYAYGRNAVYPEGHHGNAVLSRFPIEYYENRDISVGNGEKR  GLLYCRIVPPRSGITIHVICVHLGLRADQRQAQLTMLAEWVNTLPAGEPVVVAGDFNDWR  QQANQPLKAQAGLEEIFTRARGRPARTFPVSFPLLRLDRIYVKNAHASRPKALALKQWRH  LSDHAPLSVEIHL |
| 164. | Protein_ID\|573.43724.peg.6393_Diguanylate cyclase (EC 2.7.7.65)/DgcY | MFYFFWTLPFAWPLLVILMTTGLTALYHHWPGITAFMLPLWVTALLAGIQLHYHTEIRFL  ILWAIFTAILLYGRRILQRWYDEAWDTHQENMQLIQRLESIANQDALTGTANRRALNAYL  AAIWQQKTPLALMMIDVDYFKRYNDRYGHQAGDECLSSVAQVLKMAVRAEGDLVARYGGE  EFVVVLPGVSLAHATAIAERIQQKIREAGLPHAASAVASEVTVSIGIVASDGTVPIETLI  ARADSALYQAKNKGRNQWSY |
| 165. | Protein_ID\|573.43724.peg.6478Tetraacyldisaccharide 4'-kinase (EC 2.7.1.130) | MIARIWSGESPLWRLLLPLSWLYGLVSGVIRLSYQLGWQKAWRAPVPVVVVGNLTAGGNG  KTPVVIWLVEQLQQRGIRVGVVSRGYGGKAERYPLVLDDRTSTALAGDEPVLIHQRTGAP  VAVAPLRSDAVKALLSAHDLQMIVTDDGLQHYNLARDREIVVIDGVRRFGNGWWLPAGPM  RERASRLQSVDAVIVNGGVARPGEIPMRLRQGWRLTCLPANVAMSPPLRTWWRWPASAIR  RASSPPSKAAASSRSKPWRWRTIRR |
| 166. | Protein_ID\|573.43724.peg.6484_3-deoxy-manno-octulosonate cytidylyltransferase(EC 2.7.7.38) | MVMDAKGYALYFSRATIPWDRDRFAQSRETIGDSLLRHIGIYGYRAGFIRRYVSWAPSPL  EQIEMLEQLRVLWYGEKIHVAVAAEVPGTGVDTPEDLERVRAELR |
| 167. | Protein_ID\|573.43724.peg.6549_Outer membrane protein A precursor | MDDNEAQKMKKTAIAIAVALAGFATVAQAAPKDNTWYAGGKLGWSQYHDTGFYGNGFQNN  NGPTRNDQLGAGAFGGYQVNPYLGFEMGYDWLGRMAYKGSVDNGAFKAQGVQLTAKLGYP  ITDDLDIYTRLGGMVWRADSKGNYASTGVSRSEHDTGVSPVFAGGVEWAVTRDIATRLEY  QWVNNIGDAGTVGTRPDNGMLSLGVSYRFGQEDAAPVVAPAPAPAPEVATKHFTLKSDVL  FNFNKATLKPEGQQALDQLYTQLSNMDPKDGSAVVLGYTDRIGSEAYNQQLSEKRAQSVV  DYLVAKGIPAGKISARGMGESNPVTGNTCDNVKARAALIDCLAPDRRVEIEVKGYKEVVT  QPAA |
| 168. | Protein_ID\|573.43724.peg.6575_Type I secretion membrane fusion protein, HlyD family @ Type I secretion system, membrane fusion protein LapC | MNKPTAAIFPLVKELDPVAAMADNERDEAELVKSRRLIALLALLLAVTGVWAWFATLDEV  STGTGKVIPSSREQVLQTLDGGILTELNVREGSRVAAGGGRTIDPTRSESNVGESQAKYR  ASLAASIRLTAEVNNQPLIFPPSLKAWPGLLAEETRLYHSRREQLTKSMRQLEQSLSLVN  SELAINEKLAKTGAASNVEVLRLRQQAADIELKKIDLNTRYYVDAREQLSKANADVASLA  EVIKGRADSVARLTVRSPVQGIVKNIKVNTIGGVIAPNGELMDIVPIDGRLLIEARISPR  DIAFIHPDQKALVKITAYDYAIYGALNGVVETISPDTIQDEAKPDVYYYRVFIRTDHNYL  ENKRGKRFLIGPGMIATVDIKTGEKTVMDYLVKPFNRAKEALRER |
| 169. | Protein_ID\|573.43724.peg.6660_Peroxyureidoacrylate/ureidoacrylate amidohydrolase RutB (EC 3.5.1.110) | MLIIWFQNGWDDQYVEAGGPGSPNYHKSNALKTMRQRPELQGKLLAKGGWDYQLVDELTP  QEGDIVLPKPRYSGFFNTPLDSILRSRCIRHLVFTGIATNVCVESTLRDGFFLEYFGIVL  EDATHQAGPAFAQQAALFNIETFFGWVSDVESFCHALSPAAPLALAKEKRYA |
| 170. | Protein_ID\|573.43724.peg.6711Lipid A biosynthesis lauroyl acyltransferase(EC 2.3.1.241) | MTHLPKFSPALLHPRYWLLWLGIGLLWLVVQLPYPLIYRLGNAIGRLAMRFMKRRAKIAY  RNLELCFPEKSEQERHRMVVMNFESVGMGLMETGMAWFWPVRRIARWTDTIGFEHIRDVQ  AQQRGILLIGIHFLTLEMGARMFGMNEPGIGVYRPNDNPVIDWLQTGAACAPTRI |
| 171. | Protein_ID\|573.43724.peg.6726_Peptidoglycan lipid II flippase MurJ | MNLLKSLAAVSSMTMFSRVLGFARDAIVARIFGAGMATDAFFVAFKLPNLLRRIFAEGAF  SQAFVPILAEYKSKQGEDATRVFVSYVSGLLTRPWLSLPLSGCWRRRG |
| 172. | Protein_ID\|573.43724.peg.6727_Peptidoglycan lipid II flippase MurJ | MGAILNTWNRFSVPAFAPTFLNVSMIGFALFAAPYFHPPVLALAWAVTVGGVLQLAYQLP  HLKKIGMLVLPRINLKDAGAMRVVKQMGPAILGVSVSQISLIINTIFASFLVSGSVSWMY  YADRLMEFPSGVLGVALGTILLPSLSKSFASGNHDEYCRLMDWGLRLCFLLALPSAVALG  ILAKPLTVALFQYGKFSAFDAAMTQRALVAYSVGLMGLIVVKVLAPGFYSRQDIKTPVKI  AIITLIMTQVMNLAFIGPLKHAGLSLSIGLAACLNAALLYWQLRKQKIFTPQPGWLAFLL  RLIIAVLVMAAALLGVMHLMPEWSLGTMPFRLMRLLAVVIAGVVAYFATLLVLGFRVKEF  VRRTA |
| 173. | Protein_ID\|573.43724.peg.6790_4-hydroxythreonine-4-phosphate dehydrogenase(EC 1.1.1.262) | MQTVPDRLGTDYQPGDENHGCDFAIHKIAHPQEARFSWGTLDVLETGDYDCDSIEWGKVQ  KLAGQMSLDYVMKSIELGKAGLIDVVSTAPIHKEAIKLAGCKLPGHTEIYQVETQSDYGL  TMFHVHNLRVFFVSRHMALKAACDYANKARVLACVQQIHHEFTALNIKNPRIAVAALNPH  GSDNGLFGHEEADNLIPAVKAAQEMGINAIGPVPADSVFHLGKQGRYDAILSLYHDQGHI  ACKTLDFERSITITFGLPFMRSSVDHGTAFDIAGTGKAGTVSMLESTLVAARYWKMKHQ |
| 174. | Protein_ID\|573.43724.peg.6819_Sensor histidine kinase PhoQ (EC 2.7.13.3) | MKGLLRHIFPLSLRVRFLLATAGVVLVLSLAYGMVALVGYSVSFDKTTFRLLRGESNLFY  MLARWENGAIDVDIPENLNMESPTVTLIYDEQGKLLWAQRDVPWLAKRIQPEWLKRNGFH  EIEADVDSSSMLLRNNHEIQEQLDAIREQGDDSEMTHSVAINLYPATSKMPQLSIVVVDT  IPVELKRSYMVWSWFVYVLAANLLLVIPLLWVAAWWSLRPIESLAKEVRELEEHHREKLN  PNTTRELTRLVSNLNRLVRSERERYDKYRTTLTDLTHSLKTPLAVMQSTLRSLRGEKISV  DEAEPVMLEQISRISQQIGYYLHRASMRSGGTLLSRELHPIAPCLTA |
| 175. | Protein_ID\|573.43724.peg.6865_Lipid A biosynthesis palmitoleoyltransferase(EC 2.3.1.242) | MACVFNKQLLHPRNWLTWFGLGILWLIVQLPYPLLHFIGTSAGRLSRRFLKRREHIARRN  IELCFPDMSPAARETLIDQNFMSLGMGLIETGMAWFWSDERVKKWFDVEGFANLNHALSG  GKGVMVVGVHFMSLELGGRAMGLCRPMMATYRPHNSPLMEWVQTRGRLRSNKAMIDRNNL  TGLVHALKSGEAVWFAPDQDYGPKGSVFAPFSPSRRRRPPTAPMSCPAYRARKC |
| 176. | Protein_ID\|573.43724.peg.6933_2-nitroimidazole transporter NimT | MTLFLGINSLVYYVIIGWLPSILQSMGYSEAQAGSLHGLLQLATAAPGLAIPLILHRLRD  QRGIAVLVALMCAISAAGLWLLPELAIGWTLLFGFGSGATMILGLTFIGLRASSAHQAAA  LSGMAQSVGYLLAACGPPLMGRIHDANGDWHIPLLAVALISLVMAVCGALAGRDREIHP |
| 177. | Protein_ID\|573.43724.peg.7138_Psp operon transcriptional activator | MAKFIMAQYKDNLLGEANSFLEVLEQVSRLAPLDKPVLVIGERGTGKELIANRLHYLSSR  WQGPFISLNCAALNDNLLDSELFGHEAGAFTGASKRHPGRFERADGGTLFLDELATAPML  VQEKLLRVIEYGELERVGGSQPLQVNVRLVCATNADLPQMVEEGHFRADLLDRLAFDVVQ  LPRYATGRATSCCSPTSLPSRCAANSVCRCFPALANERPRRCSATAGRGIFAN |
| 178. | Protein_ID\|573.43724.peg.7152_Transcriptional repressor protein TyrR | MRLEVFCEDRLGLTRELLDLLVLRGIDLRGIDIDPIGRIYLNFAELEFATFSSLMAEIRR  IAGVTDVRTVPWMPSEREHLALSALLVAMPEPVLSLDTKGRVELANPASCLLFGQSQAKL  RNHPVAQLIADFNVQRWLESSPQETHAEHVVVNGQNYLLEVTPVYLEGEHNERVLTGAVA  MLRSTVRMGRQLQTMTSQDTSAFSQILAVGPKMRHVVEQARKLAMLSAPLLIVGDTGTGK  DLLAHACHLASPRAGKPYLALNCGSIPEDAVESELFGDALQGKKGFFEQANGGSVLLDEI  GEMSPRMQTKLLRFLNDGTFRRVGEDHEVHVDVRVICATQKNLIELVQKGLFREDLYYRL  NVLTLYLPPLRDCPQDIMPLTELFVARFADEQGIPRPKLSADLSTVLTRYSWPGNVRQLK  NAVYRALTQLEGFELRPQDILLPDHDVASLPVGEEAMEGSLDDITRRFERSVLTQLYRSY  PSTRKLAKRLGVSHTAIANKLREYGLSQKKGDE |
| 179. | Protein_ID\|573.43724.peg.7156_Thiol peroxidase, Tpx-type (EC 1.11.1.15) | MSQTVHFVGNPVSVQGTIPQAGAKAQPFTLVAKDLSDVALSQYAGKRKVLNIFPSIDTGV  CAASVRKFNQLAAELDNTVVLCISADLPFAQSRFCGAEGLSNVVTLSTLRGASFLADYGV  AIATGPLAGLAARAVVVIDENDQVVYSQLVNEITEEPDYDAALAALKA |
| 180. | Protein_ID\|573.43724.peg.7210_T6SS component TssC (ImpC/VipB) | MSVTTENAPVQGQTTLQENRAGEGVYASLFEKINLTPASRLGDINDFLDDAALSEAPAAE  RLTAAMQVFMERIRQSGQRVEKLDKTLIDHHIAELDFQISRQLDAVMHHQEFQQVESLWR  GLKQLVDNTDYRQNVKTEILDVAKDDLRQDFEDAPELIQSGLYWHTYTAEYDTPGGEPIG  SVISAYEFDASPQDVALLRNISRVSAAAHMPFIGAVGPAFFLKETMEEVAAIKDIGNYFD  RAEYIRWKAFRETDDARYIGLVMPRVLGRLPYGPDTVPVRSFNYVEQVKGPDHEKYLWTS  AAFSFASNMVKSFVNNGWCVQIRGPQAGGAVKDLPIHLYDLGTGNQVKIPSEVMIPETRE  FEFASLGFIPLSYYKNRDYACFFSANSAQKPALYDTADATANSRINARLPYIFLLSRIAH  YLKMIQRENIGTTKDRRLLELELNTWVRSLVTEMTDPGDELQASHPLRDASVVVEDIEDN  PGFFRMKLYAVPHFQVEGMDVNLSLVSQMPKAKA |
| 181. | Protein_ID\|573.43724.peg.7219_VgrG protein | MSSVKSLLFSHNHHLLSVKGCEAGLDVLAFEGDEALSQPFRYRIEFTSADHAISKEMMLM  KAASLTLQAPVAQGFGINVQQPVRVIQGVVTGFERLSTSRDETHYALTLQPRLALLNRSH  QNAIYQDQSVPQIVEKILRERHGLRGQDFLFSLTKTYPRREQVMQYGEDDLRFITRLLGE  VGIWFRFTADTRLHIDVAEFCDSQQGYEKGLTLPSVPPSGQQSAGVDAVWEMACRHRVVE  QQVSTRDYNYREATADMNAQVDVTRGETTTFGEAYHWGDNYLTAGNVHDRHPAPESGAFY  ARLRHERYLNGQTRMQATTSCPTLCPGQVLKVTGGEEVAGEFADGVLVTAMHSHARRDAD  FAVEFAGIPDSPDVGYRPEPGARPVMAGTLPARVTSTRENDTYGHIDKHGRYRVNMLFDR  ARWETGFESLWVRQSRPYAGDTYGLHLPLLAGTEVAIGFEDGNPDRPYIAGVLHDSAHGD  HVTIRNDKRNVLRTPANNKIRLDDERGKEHIKVSTEYGGKSQLNLGHLVDSDRQPRGEGF  ELRTDSWGAIRAQKGIFISADGQAQAQGQVLAMEPAVSNLAEAREQMMSISGDAQKATAN  PADLQAQITLLEQQLTDLKKSVLLVSAPEGIALTSGEHLQVSAGHNLIATAGKNADVSVV  KNLFIGVGSALSVFVRKLGIRLIANQGPVQMQAQNDLMALLARKEISIVSTEDSIEIIAK  KRVTINGGGSYITLNASGIESATAGEYRTRAGYYVRREKAQHKPDIAPLANAINDGSHNI  RYLCTDDNGVPMMNTPYRAFLADGSVLEGVSDGEGYTKLFTSAQVQDVLLHMIPEAINA |
| 182. | Protein_ID\|573.43724.peg.7238_T6SS component TssF (ImpG/VasA) | MDDLTLRYYDAEMRYLLEAGEEFARAHPEQAAMLNLDKAGARDPFVERLFEGFAFLMGRM  REKLDDDLPELTEGVVSLLWPHYLRTIPSMSVVEFTPDWPEMKEPMTMAKGFEVLSRPIG  EKGTRCRYTTTKAIALQPLSLERVQLATDTDGRSVITLRFTCSQLTDWRRVDLSHIPLYC  NADAPLACAMHEAFTLNVARMWLRMPDEVDRRPLDGYFSALGFGEDDGLWPEDGRSFRGY  QLLLEYFTFREKFMFIDLRGLETVAFPAGLAWFEIDVVLAERWEHDFRFSEKQLRLHCVP  VINLFPLESDPLTINSLQTEYPLRPMRVQDGHTEIYTVDSVISSHQQVYAPFSSFRHKGG  MMRHDTADYYYHTRVRRGPSGLYNTWLIVGGEAFDNHTVPEDESLSLTLTGTNGQLPRRA  LQSTVLDTVMKTTSASIAVRNLCAPTLPCYPPPRSVFTGGC |
| 183. | Protein_ID\|573.43724.peg.7331_Efflux transport system, outer membrane factor(OMF) lipoprotein | MIRPVALAIVLALVGCQSADVPRAQPTLTIPAAWRADVGPASPVEGVWWRNFHDSTLNQY  VDQALRYNSDVLIARERVNEYQARAYAADSSLFPSLDASLTGTRARTQSAATGLPIHSTL  YKGGLTASYDVDIWGANRSAANTAGASLEAQKAAAAAANLSVASSVAVGYVTLLSLDEQL  RVTQQTLTSREDAWRLAKRQFETGYTSRLELMQADSELRSTRAQIPPLQHQIAQQENALS  VLLGDNPGAVKRGEFAQLTPLRLPSQLPSTLLNRRPDIAQAERQLVAADATLASSQAQLL  PSINLTATGSLQDRTLPDLLDNPLRLWSLGGSILVPLLNRQALNAQVDVSMAQRNQALYS  YEKTVRSAFKEVNDSLDAISRYGEQLTELQEQETVAQETLRIAQNRYRNGYSSYLDVLDA  QRTLFSTQLSVVQVKNNLLLAQIDLYRALGGGWSDSSGS |
| 184. | Protein_ID\|573.43724.peg.7332_Membrane fusion component of MSF-type tripartite multidrug efflux system | MSQQDAAKQQANTRNNIRVVSIFTAAAIGLVGVLVILYAWQLPPFTRHSQFTDNAYVRGQ  TTFISPQVNGYITAVNVKDFAIVQPGEVLFQIDDRIYKQRVHQAQATLAMKEAALRNNLQ  QRKSAEATIAKNEAALQNARAQNLKIQADLKRIQQLTADGSLSIRERDSARASAAQGAAD  IEQAKAALEMSRQDRESTIVNRDSLEADVASAKAALELAQIDLQNTQIIAPTGGQLGQIS  VRLGAYVSAGTHLTSLVPPQHWVIANLKETQLAEVRVGQPVTLTVDALNGETFHGKVQSI  SPATGVEFSAISPDNATGNFVKIAQRIPVRITVNDGQNNSERLRPGMSVQVTIDTRAEKQ  P |
| 185. | Protein_ID\|573.43724.peg.7481_Anaerobic selenate reductase, molybdenum cofactor-containing periplasmic protein | MSNQESPGGVSRRALLKSTALGSLALAAGGLTLPFTLRRAAAAVQQATGDTTRIVWGACS  VNCGSRCALRLHVRDDEVVYVETDNTGDDRYGDHQVRACLRGRSIRRRINHPDRLNYPMK  RVGKRGEGKFVRISWQEALDTLADRLKSVVAQYGNEAVYINYSSGIVGGNITRSSPSASP  VARLMNCYGGSLNQYGTYSTAQIACAMPYTYGSNDGNSTSDIENSKLVVMFGNNPAETRM  SGGGITWYLEQARERSNARMIVIDPRYTDTAAGREDEWIPIRPGTDAALVAGIAWVLINE  DLVDQPFLDKYCVGYDEKTLPAGAPANGHYKAYILGEGDDGIAKTPQWASRITGIPTERI  IKLAREIGMSKPAYICQGWGPQRQANGELTARAIAMLPILTGNVGINGGNSGARESTYTI  TIERLPVLENPVKTAISCFTWTDAIARGPEMTASRDGVRGKEKLDVPIKFLWNYAGNTII  NQHSDINKTHEILQDESKCETIVVIDNFMTSSAKYADLLLPDLMTVEQEDIIPNDYAGNM  GYLIFIQPATSAKFERKPIYWILSEVAKRLGDDVHQRFTEGRTQEQWLQYLYAKMVAKDP  ALPAYEDLKRMGIYKRKDPNGHFVAYRDFRRDPEAHPLKTPSGKIEIYSSRLAEIAARWQ  LEKDEVISPLPVYASTFEGWDDPLRSRYPLQLFGFHYKARTHSSYGNVDVLQAACRQEVW  INPLDAEKRGIKNGDMVRVFNQRGEVRLPAKVTPRIMPGVSAMGQAPGTTPI |
| 186. | Protein_ID\|573.43724.peg.7523_3-oxo-tetronate kinase | MKLVSGGSGLAIGLARDWAQRHGARGESAQAGMPLAGPAVVLSGSCSVMTNSQVAAYRQQ  APARAVDLSACFTDLESYVRTLTDWVDAQRDAPLAPMIYATTEPQTLQRIQAQYGDKASS  ERVEQLFAALAAALKAKGFTRFIVAGGETSSIVAQTRG |
| 187. | Protein_ID\|573.43724.peg.7524_3-oxo-tetronate kinase | MQLGVIADDFTGATDIASFLVRNGMPTVQLNGVPTRDLPLTSEAVVISLKTRSCPAEMAV  SQSLAALRWLQAQGCQQFYFKYCSTFDSTAQGNIGPVLDALLAELGETRTVISPALPVNG  RTVYQGYLFVGEQLLNESGMRHHPVTPMEDAHLGRLIERQGRGKAALIAWPIVARGRRRS  PPRWRQSTIRRCTMWCSTPSVNRICSPRAWRCGR |
| 188. | Protein_ID\|573.43724.peg.7644_type 1 fimbriae anchoring protein FimD | MRHGPAHTVTVVSLSILLGGQSALLHAQATFNMDLLEKNDHLPAVDLQRFNQQAGQPPGA  YPVSWQVNGVTLDARKTVTFRQNDRGQLTPCLKPEDLLQAGVNPAVLSQATGATSRSCPE  LNALLPGSTVNFDFAHQRLVMTIPQALMTHRARDNVPSALWDEGISAFQSNYRYSGASQR  TREGSTERDNYLMLKSGVNVGAWRLRASNNLTANSDDKPQWTTSGAWLERDLTRWQSELT  LGDTFTSGDVFDAVQFQGISLASSDAMLPDSQKGFAPTIRGIARTNAQVTVRQNGYVLYQ  TYVTPGAFVIDDLYPTASSGNLEVAVKESDGEIRRFTQPYASVTSMQREGSLKYNLVAGR  YHSDDASQRPLMMQLSLMRGFAHNLTLFGGLQSAAQYHNLSLGAGQGLGEAGALSLQLLN  ARDRHQQDPIDGRAWQLQYSKGFDRLGTQLTFTGWRYSHQRYATLSEAFSSPGSDDDLQD  SDNKKATLQITASQSLPYDITLYLSLDQDSYWSGGATQRTANMGISSQVHGIAWSLSYSD  SRSSHGDEEDDEPHSDKVVTLSLSVPLSHLLPGSYAGYTLTSSRHSVGSQMVSLNGTLLD  NHALSYAVSQTRDRQNGSSGSLTAGYSSGRGDLNLGYSHDSQAARLNYGASGAF |
| 189. | Protein_ID\|573.43724.peg.7654_hypothetical protein | MAGAVLKTLQKEEMVARFGGDEFVAVKPFSDEGEVDAFAARLWHCFSGKQTFAATEVVLS  ASIGISVYPEDGTDINTILSNSDLAMYRAKSSLDHKICWYEREMDDKTRQRNMMAADIRR  GIHAGEFSLHYQAIRNIKDRSITGYEALLRWQHPQLGTIPPDVFIPIAEESGAIVPLGYW  VLEQVCNESLENGLNRKVSVNISPVQLRHRSFIEKVREILMRTAYPVSLLEFEVTETAFV  INKQLAFSVLHHLQKMGISIALDDFGTGYSSLSMLRDFHFDVIKLDRSFMTDVESNPQVR  SFVRAIISLGNSINTPLIAEGVETAGQLQILEEEGCDEMQGFLFGEPVDIKHLPDRR |
| 190. | Protein_ID\|573.43724.peg.7743_Uncharacterized LysM domain protein YgaU | MGLLNFVKEAGEKIWDAVSGDSKEDRADKLKKHIDGLNLPGAEKVNIDVAEDGTATVTGD  VASQEDKEKILVAVGNVTGVGQVSDGVKVTQSGAESRFYTVKSGDTLSAISKAMYGSAND  YQRIFEANKPMLTHPDKIYPGQVLIIPAK |
| 191. | Protein_ID\|573.43724.peg.7892_Catalase-peroxidase KatG (EC 1.11.1.21) | MSTSNDPSNNASAGKCPFHAETPKQSAGSGTANRDWWPNQLRVDLLNQHSNRSNPLGENF  NYREEFKKLDYSALKADLRALLTDSQEWWPADWGSYIGLFIRMAWHGAGTYRTVDGRGGA  GRGQQRFAP |
| 192. | Protein_ID\|573.43724.peg.7893Catalase-peroxidase KatG (EC 1.11.1.21) | MKQKYGQKISWADLYMLAGNVALENAGFRTFGFGAGREDVWEPDLDVDWGDEKEWLAHRH  PESLAKQAIGATEMGLIYVNPEGPNASGEPLSAAAAIRATFGNMAMDDEEIVALIAGGHT  LGKTHGAAETSHVGAEPEAAPLEAQGLGWHSSYGSGAGADAITSGLEVVWTQTPTQWSNY  FFENLFKYEWVQTRSPAGAIQFEAKDAPEIIPDPFNPEKKRKPTMLVTDLTLRFDPEFEK  ISRRFLNDPQAFNEAFARAWFKLTHRDMGPKSRYLGPEVPKEDLIWQDPLPAATHQPSAE  DIASLKSAIAGAGLSVSELVSVAWASASTFRGGDKRGGANGARLALAPQKDWPVNAIASR  VLPTLQAIQRASGKASLADIIVLAGVVGVEQAAAAAGVSVNVPFTPGRVDALPEQTDVES  FDLLQPLADGFRNYRRIEGGVSTETLLIDKAQQLTLTAPEMTVLVGGLRVLGANYDGSKH  GVFTDRVGVLSNDFFVNLLDMATVWKAADDNAELFTGSDRKTGEAKYSATRVDLVFGSNS  VLRALAEVYACADGQQKLVHDFVAAWTKVMNLDRFDL |
| 193. | Protein_ID\|573.43724.peg.7985_Benzoate 1,2-dioxygenase alpha subunit (EC 1.14.12.10) | MQKTLSTLKDKINNALVVDRENHIYRCHRSIFTDPQLFEFEMKHIFEGNWVFLAHESQIP  QPGDYYTLTLGRQPVIITRDKKNELHALINSCAHRGAMLCRRKTGNKNSFTCPFHGWTFS  NNGKLLKAKDESTGAYPETFKHEGSHDLQKLPRFQSYRGFLFGSLNADVQPLEAYLGETC  KIIDLIVDQAPEGLEVLKGSSSYVYEGNWKLGAENGADGYHVSVVHWNYASTMSRRNYEA  EGTHTVDANGWSKSLGGGYGFDNGHMLLWTRALNPEVRPVYAHRERLQAEFGERRADQMV  NETRNLCLYPNVYLMDQFSTQIRVIRPIAVDKTEVTIWCFAPKGESDQARALRIRQYEDF  FNVSGMGTPDDLEEFSACQRGYLGENLPWSDLSRGALRWVDGADEHAQHAGFSPRLSGVK  SEDEALYIAHHHHWQTLMLAAIEQEQQRYDQSITQRVEVA |
| 194. | Protein_ID\|573.43724.peg.8005_Respiratory nitrate reductase delta chain (EC 1.7.99.4) | MRILKVIGLLLEYPDELLWENRDDALALVRADVPSLTPFVSELLTAPLLDRQAEWCEVFE  RGRATSLLLFEHVHAESRDRGQAMVDLMNQYQQAGLQIDCRELPDHLPLYLEYLSILPPA  EAREGLQNIAPILALIGGRLKQRACPYYQLFDALLALAKSPLTSDSVTKQVAGEKRDDTR  QALDAVWEEEQVKFIEDNATACDSSSMQAYQRRFSQDVAPQYVDIRAGGPK |
| 195. | Protein_ID\|573.43724.peg.8050_Uncharacterized protein YncE | MQDDGKAHFYLNLSLDTAGHRAFITDSKQPEVLVVDTRDGKVLEKIAAPESLAVLFNPAR  NEAYVTHRKAGEVSVIDGKSYKVVKTFKTPTHPNSLALSEDGKTLYVSVKQASSREKEAT  APDDVIRIAL |
| 196. | Protein_ID\|573.43724.peg.8164_Transposase | MLMCDATGLSQRRACRLTGLSLSTCRYEAHRPAADAHLSGRITELALERRRFGYHRIWQL  LRREGLHVNHKRVYRLYHLSGLGVKRRRRRKGLATERLPLLRPAAPNLTWSMDFVMDALS  TGRRIKCLTCVDDFTKECLTVTVAFGISGVQVSRILDSIALFRGYPATIRTDQGRSSLAV  HWINGPLSMVLSCA |
| 197. | Protein_ID\|573.43724.peg.8856_hypothetical protein | MAVGKKIKTIDGLGHAVTGIALNKNGDNIYLVNGDGEIINLDASSYEIKKRFVVEPEKKH  FFLNVSLDEKNGRAFITDPDIPGVLVVDVNSGKILHRIDVINSLAILFNPLRNEIYITHR  NAKQVSIVDSNSYYVKTTIKTRLMPNSLSLSHDGENLYVSVKQEKKR |

Supplementary Table 4. Statistics for pan-genomics and ST of studied *K. pneumoniae* genomes.

| Genome no. | Organism name | No. of core genes | No. of accessory genes | No. of unique genes | No. of exclusively absent genes | ST according to Pasteur institute scheme |
| --- | --- | --- | --- | --- | --- | --- |
| 1. | 3189STDY5864735 | 1303 | 3941 | 1 | 0 | 69e0273f2bf9ad1162108683eae6b29c6e2a76a4 (Novel ST) |
| 2. | 3189STDY5864736 | 1303 | 3934 | 8 | 1 | 711 |
| 3. | 3189STDY5864738 | 1303 | 3944 | 5 | 1 | 711 |
| 4. | 3189STDY5864739 | 1303 | 4426 | 24 | 0 | 59d9e867964a1a508a6f94d4b05f15995c241ff5 (Novel ST) |
| 5. | 3189STDY5864742 | 1303 | 4014 | 3 | 2 | 79d0aa5063336acb237d29921fdbfa7dfde9b3f7 (Novel ST) |
| 6. | 3189STDY5864746 | 1303 | 4414 | 3 | 1 | 15 |
| 7. | 3189STDY5864747 | 1303 | 3933 | 4 | 1 | 711 |
| 8. | 3189STDY5864748 | 1303 | 4366 | 1 | 0 | b4008278f0c9c2eda18cd211e7105684ca78d6d3 (Novel ST) |
| 9. | 3189STDY5864754 | 1303 | 4200 | 7 | 2 | 336 |
| 10. | 3189STDY5864755 | 1303 | 4173 | 3 | 0 | 336 |
| 11. | 3189STDY5864759 | 1303 | 4198 | 0 | 0 | 336 |
| 12. | 3189STDY5864760 | 1303 | 3894 | 77 | 0 | 13 |
| 13. | 3189STDY5864763 | 1303 | 4186 | 2 | 0 | 336 |
| 14. | 3189STDY5864774 | 1303 | 4158 | 8 | 1 | 22 |
| 15. | 3189STDY5864776 | 1303 | 4101 | 57 | 0 | 437 |
| 16. | 3189STDY5864778 | 1303 | 4354 | 4 | 0 | 15 |
| 17. | 3189STDY5864780 | 1303 | 4347 | 10 | 2 | 15 |
| 18. | 3189STDY5864781 | 1303 | 4144 | 7 | 2 | 22 |
| 19. | 3189STDY5864784 | 1303 | 4505 | 5 | 0 | 336 |
| 20. | 3189STDY5864786 | 1303 | 4011 | 10 | 0 | 711 |
| 21. | 3189STDY5864788 | 1303 | 4130 | 8 | 0 | 278 |
| 22. | 3189STDY5864790 | 1303 | 4535 | 13 | 1 | 336 |
| 23. | 3189STDY5864792 | 1303 | 4314 | 164 | 3 | 292 |
| 24. | 3189STDY5864793 | 1303 | 4306 | 9 | 1 | 3805 |
| 25. | 3189STDY5864795 | 1303 | 4303 | 6 | 1 | 3805 |
| 26. | 3189STDY5864796 | 1303 | 4115 | 10 | 1 | 278 |
| 27. | 3189STDY5864799 | 1303 | 4300 | 5 | 1 | 3805 |
| 28. | 3189STDY5864800 | 1303 | 3932 | 4 | 0 | 29 |
| 29. | 3189STDY5864801 | 1303 | 3922 | 12 | 0 | 29 |
| 30. | 3189STDY5864809 | 1303 | 3837 | 120 | 1 | 442 |
| 31. | 3189STDY5864813 | 1303 | 4191 | 5 | 0 | 15 |
| 32. | 3189STDY5864814 | 1303 | 4318 | 47 | 0 | 15 |
| 33. | 3189STDY5864815 | 1303 | 4397 | 14 | 0 | 15 |
| 34. | 3189STDY5864816 | 1303 | 4231 | 41 | 0 | 336 |
| 35. | 3189STDY5864822 | 1303 | 4061 | 6 | 0 | 15 |
| 36. | 3189STDY5864827 | 1303 | 4292 | 77 | 0 | 474 |
| 37. | 3189STDY5864831 | 1303 | 3930 | 92 | 0 | 29 |
| 38. | 3189STDY5864837 | 1303 | 4322 | 1 | 0 | 15 |
| 39. | 3189STDY5864838 | 1303 | 4327 | 3 | 0 | 15 |
| 40. | 3189STDY5864842 | 1303 | 4071 | 8 | 0 | 17 |
| 41. | 3189STDY5864845 | 1303 | 4338 | 5 | 1 | 15 |
| 42. | 3189STDY5864846 | 1303 | 4338 | 132 | 0 | 3278 |
| 43. | 3189STDY5864848 | 1303 | 4314 | 2 | 0 | 15 |
| 44. | 3189STDY5864853 | 1303 | 3796 | 4 | 1 | 48 |
| 45. | 3189STDY5864855 | 1303 | 4098 | 7 | 0 | b4008278f0c9c2eda18cd211e7105684ca78d6d3(novel ST) |
| 46. | 3189STDY5864858 | 1303 | 3954 | 27 | 0 | 15 |
| 47. | 3189STDY5864859 | 1303 | 4098 | 3 | 0 | 15 |
| 48. | 3189STDY5864862 | 1303 | 4092 | 11 | 0 | 15 |
| 49. | 3189STDY5864864 | 1303 | 4148 | 20 | 0 | 45 |
| 50. | 3189STDY5864869 | 1303 | 4439 | 11 | 0 | 48 |
| 51. | 3189STDY5864872 | 1303 | 4405 | 7 | 1 | 15 |
| 52. | 3189STDY5864873 | 1303 | 4142 | 1 | 0 | 15 |
| 53. | 3189STDY5864874 | 1303 | 4731 | 184 | 5 | 48 |
| 54. | 3189STDY5864884 | 1303 | 4346 | 8 | 0 | 48 |
| 55. | 3189STDY5864895 | 1303 | 4455 | 26 | 0 | 48 |
| 56. | 3189STDY5864927 | 1303 | 4103 | 8 | 0 | 48 |
| 57. | 3189STDY5864934 | 1303 | 3989 | 53 | 0 | 48 |
| 58. | 3189STDY5864942 | 1303 | 4081 | 5 | 0 | 48 |
| 59. | 3189STDY5864948 | 1303 | 4439 | 7 | 0 | 37 |
| 60. | 3189STDY5864949 | 1303 | 4422 | 14 | 1 | 37 |
| 61. | 3189STDY6864256 | 1303 | 4258 | 7 | 0 | 3805 |
| 62. | 3189STDY6864257 | 1303 | 4391 | 3 | 0 | 38 |
| 63. | 3189STDY6864258 | 1303 | 4100 | 5 | 0 | 278 |
| 64. | 3189STDY6864259 | 1303 | 4082 | 3 | 0 | 15 |
| 65. | 4771 | 1303 | 4229 | 98 | 0 | 147 |
| 66. | 10718 | 1303 | 4301 | 39 | 0 | 2096 |
| 67. | CFSAN044566 | 1303 | 4237 | 23 | 0 | 14 |
| 68. | CFSAN059603 | 1303 | 3931 | 75 | 6 | 097a98f594e7282c6a6b500268e98f4c26410fef (novel ST) |
| 69. | CFSAN059608 | 1303 | 3894 | 38 | 9 | 48 |
| 70. | CFSAN059612 | 1303 | 4141 | 95 | 2 | 48 |
| 71. | CFSAN059616 | 1303 | 3957 | 82 | 0 | 37 |
| 72. | CFSAN059617 | 1303 | 3865 | 22 | 0 | 29 |
| 73. | CFSAN059622 | 1303 | 4140 | 1 | 0 | 231 |
| 74. | CFSAN059623 | 1303 | 4142 | 111 | 1 | 101 |
| 75. | CFSAN059624 | 1303 | 4452 | 28 | 0 | 437 |
| 76. | CFSAN059625 | 1303 | 4233 | 120 | 0 | 15 |
| 77. | CFSAN059626 | 1303 | 4415 | 23 | 1 | 11 |
| 78. | CFSAN059627 | 1303 | 4442 | 1 | 0 | 11 |
| 79. | CFSAN059628 | 1303 | 4097 | 4 | 0 | 11 |
| 80. | CFSAN059629 | 1303 | 4357 | 36 | 0 | 340 |
| 81. | CFSAN059630 | 1303 | 4354 | 1 | 0 | 147 |
| 82. | CFSAN059631 | 1303 | 4342 | 1 | 0 | 147 |
| 83. | CFSAN059632 | 1303 | 4596 | 18 | 0 | 147 |
| 84. | CFSAN059634 | 1303 | 4496 | 3 | 0 | 147 |
| 85. | CFSAN059635 | 1303 | 3887 | 28 | 0 | 39 |
| 86. | CFSAN059637 | 1303 | 4350 | 3 | 0 | ee9134e03602927beb248ae12d509c9a86de052f (novel ST) |
| 87. | CFSAN059638 | 1303 | 4332 | 4 | 0 | 14 |
| 88. | CFSAN059639 | 1303 | 4511 | 4 | 0 | 147 |
| 89. | CFSAN059640 | 1303 | 4419 | 63 | 0 | 15 |
| 90. | CFSAN059641 | 1303 | 4338 | 6 | 0 | ee9134e03602927beb248ae12d509c9a86de052f (novel ST) |
| 91. | CFSAN059642 | 1303 | 4327 | 32 | 0 | 17 |
| 92. | CFSAN059643 | 1303 | 4435 | 0 | 0 | 11 |
| 93. | CFSAN059644 | 1303 | 4253 | 77 | 0 | 20 |
| 94. | CFSAN059645 | 1303 | 4197 | 45 | 1 | 9cbbf4ffbf4e01295f1d57b0206129d7e50c3dd1 (novel ST) |
| 95. | CFSAN059646 | 1303 | 4449 | 6 | 6 | 14 |
| 96. | DA48896 | 1303 | 4283 | 71 | 14 | 147 |
| 97. | JRCGR1 | 1303 | 4390 | 1095 | 31 | 2096 |
| 98. | K184 | 1303 | 4271 | 1607 | 10 | 2096 |
| 99. | KP_008 | 1303 | 4100 | 10 | 0 | 337 |
| 100. | KP_017 | 1303 | 4063 | 4 | 0 | 337 |
| 101. | KP_030 | 1303 | 4340 | 7 | 0 | 3736 |
| 102. | KP_040 | 1303 | 4175 | 1 | 0 | 231 |
| 103. | KP_044 | 1303 | 4202 | 2 | 0 | 231 |
| 104. | KP_046 | 1303 | 4358 | 5 | 0 | 3737 |
| 105. | KP_063 | 1303 | 4294 | 3 | 0 | 147 |
| 106. | KP_084 | 1303 | 4049 | 1 | 0 | 337 |
| 107. | KP_089 | 1303 | 4339 | 185 | 1 | 410 |
| 108. | KP_093 | 1303 | 4125 | 0 | 0 | 231 |
| 109. | KP_109 | 1303 | 4118 | 1 | 0 | 231 |
| 110. | KP_116 | 1303 | 4202 | 0 | 0 | 231 |
| 111. | KP_121 | 1303 | 4374 | 1 | 0 | 147 |
| 112. | KP_123 | 1303 | 4372 | 2 | 0 | 147 |
| 113. | KP_135 | 1303 | 4204 | 1 | 0 | 617 |
| 114. | KP_136 | 1303 | 4194 | 0 | 0 | 617 |
| 115. | KP_144 | 1303 | 4068 | 4 | 0 | 337 |
| 116. | KP_158 | 1303 | 4248 | 12 | 0 | 147 |
| 117. | KP_160 | 1303 | 4459 | 11 | 0 | 14 |
| 118. | KP_184 | 1303 | 4266 | 4 | 0 | 391 |
| 119. | KP_195 | 1303 | 4206 | 1 | 0 | 617 |
| 120. | KP_208 | 1303 | 4062 | 3 | 0 | 337 |
| 121. | KP_215 | 1303 | 4228 | 37 | 0 | 231 |
| 122. | KP_241 | 1303 | 4194 | 1 | 0 | 231 |
| 123. | KP_280 | 1303 | 4262 | 3 | 0 | 391 |
| 124. | KP_301 | 1303 | 4391 | 6 | 0 | 3739 |
| 125. | KP_307 | 1303 | 4373 | 6 | 0 | 3740 |
| 126. | KP_310 | 1303 | 4387 | 4 | 1 | 3741 |
| 127. | KP_317 | 1303 | 4386 | 5 | 0 | 147 |
| 128. | KP_320 | 1303 | 3833 | 52 | 0 | 584 |
| 129. | KP_328 | 1303 | 4222 | 7 | 0 | 147 |
| 130. | KP_343 | 1303 | 4332 | 1 | 0 | 147 |
| 131. | KP_346 | 1303 | 4333 | 3 | 1 | 3742 |
| 132. | KP1 | 1303 | 4292 | 20 | 0 | 147 |
| 133. | KP35W | 1303 | 3945 | 137 | 0 | 29 |
| 134. | KP36J | 1303 | 4135 | 167 | 0 | 15 |
| 135. | KP59J | 1303 | 4100 | 40 | 0 | 231 |
| 136. | KP75W | 1303 | 4191 | 43 | 0 | 11 |
| 137. | MMGA109 | 1303 | 4214 | 137 | 18 | 2096 |
| 138. | MMGK6 | 1303 | 4029 | 120 | 0 | 3663 |
| 139. | MMGK9 | 1303 | 4161 | 16 | 0 | 11 |
| 140. | MMGN70 | 1303 | 3989 | 66 | 0 | 231 |
| 141. | MMGN122 | 1303 | 4134 | 516 | 0 | 231 |
| 142. | MMGX21 | 1303 | 4309 | 211 | 0 | 340 |
| 143. | MMGX130 | 1303 | 4207 | 32 | 3 | 11 |
| 144. | PH9 | 1303 | 3614 | 25 | 13 | 11 |
| 145. | PH10 | 1303 | 638 | 14 | 1841 | 47b617df7a84892f319ee5d455d5894df59c33f4 |
| 146. | PH12 | 1303 | 3692 | 3 | 0 | 48 |
| 147. | PH24-1 | 1303 | 3993 | 3 | 0 | 11 |
| 148. | PH25 | 1303 | 3865 | 19 | 0 | 11 |
| 149. | PH28-1 | 1303 | 3704 | 6 | 4 | 925b9839ef92be0fa7074962c7fdd5d54b80f0a8 |
| 150. | PH38-1 | 1303 | 3851 | 6 | 0 | 925b9839ef92be0fa7074962c7fdd5d54b80f0a8 |
| 151. | PH40 | 1303 | 3825 | 9 | 0 | 925b9839ef92be0fa7074962c7fdd5d54b80f0a8 |
| 152. | PH44 | 1303 | 4460 | 8 | 0 | 15 |
| 153. | PH49-2 | 1303 | 3917 | 40 | 0 | 925b9839ef92be0fa7074962c7fdd5d54b80f0a8 (novel ST) |
| 154. | PH72 | 1303 | 4053 | 127 | 1 | 268 |
| 155. | PH73 | 1303 | 3961 | 67 | 3 | 873 |
| 156. | PH102 | 1303 | 3964 | 2 | 0 | 11 |
| 157. | PH124 | 1303 | 3926 | 6 | 0 | 11 |
| 158. | PH139 | 1303 | 3923 | 10 | 0 | 11 |
| 159. | PH150-2 | 1303 | 3903 | 10 | 0 | 6d3f160d7ff40d70aef0630d180a21a809c325b5 (novel ST) |
| 160. | PH152 | 1303 | 4173 | 25 | 0 | 437 |
| 161. | WU2 | 1303 | 4110 | 232 | 1 | 5fcc9cf6ccd867241d47109a1b4b7c75ff401e3d (novel ST) |
| 162. | WU3 | 1303 | 3926 | 25 | 0 | bc336f40a74612d68331bce57c788a68ebfaba54 (novel ST) |
| 163. | WU6 | 1303 | 3779 | 103 | 0 | 253 |
| 164. | WU7 | 1303 | 3671 | 62 | 0 | 882 |
| 165. | WU8 | 1303 | 3686 | 51 | 0 | 111 |
| 166. | WU9 | 1303 | 3363 | 24 | 2 | 200 |
| 167. | WU10 | 1303 | 3606 | 39 | 0 | 5064 |
| 168. | WU12 | 1303 | 4179 | 104 | 2 | 258 |

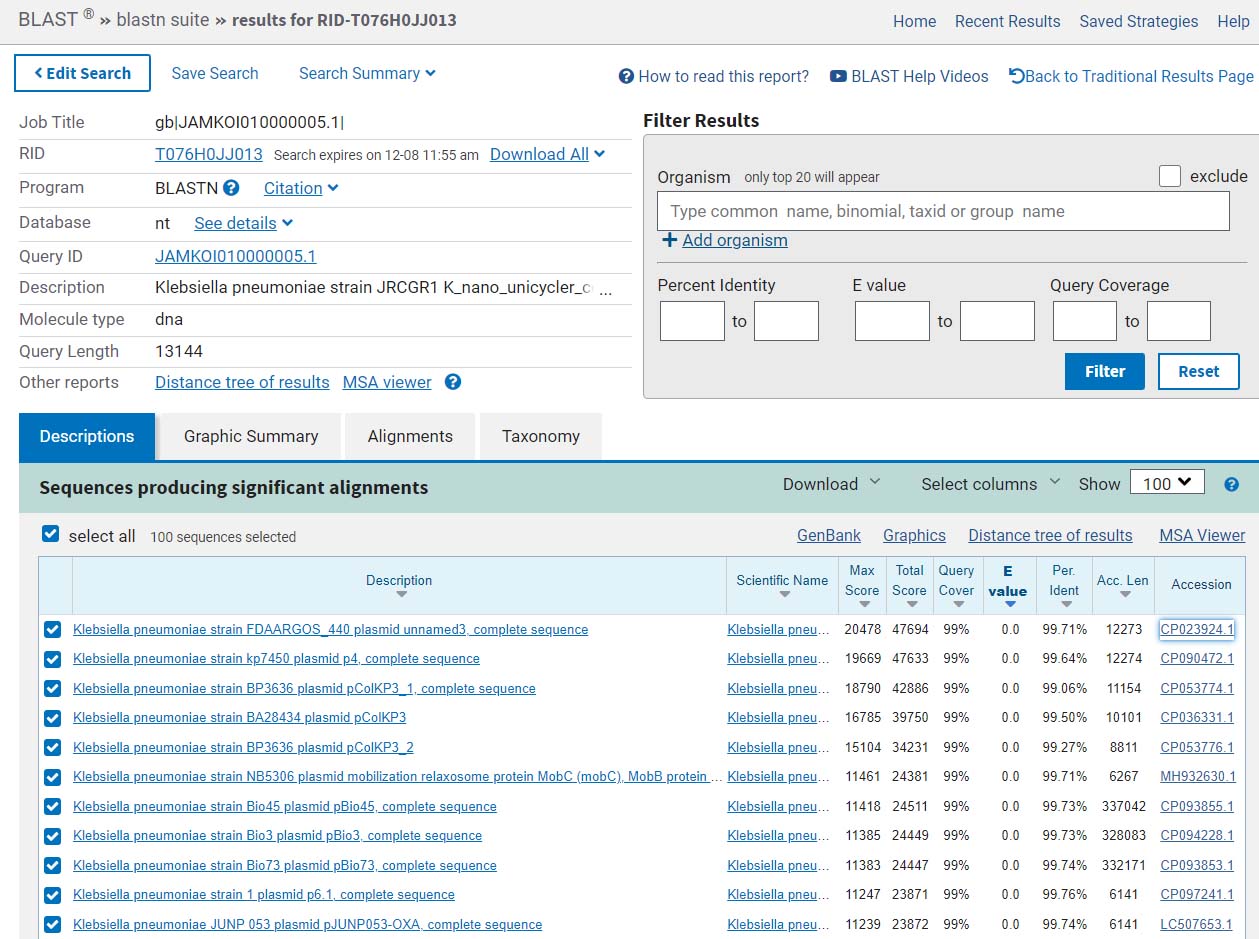

### Supplementary Fig. 1A. BLAST results confirming contig 5 as plasmid sequence.

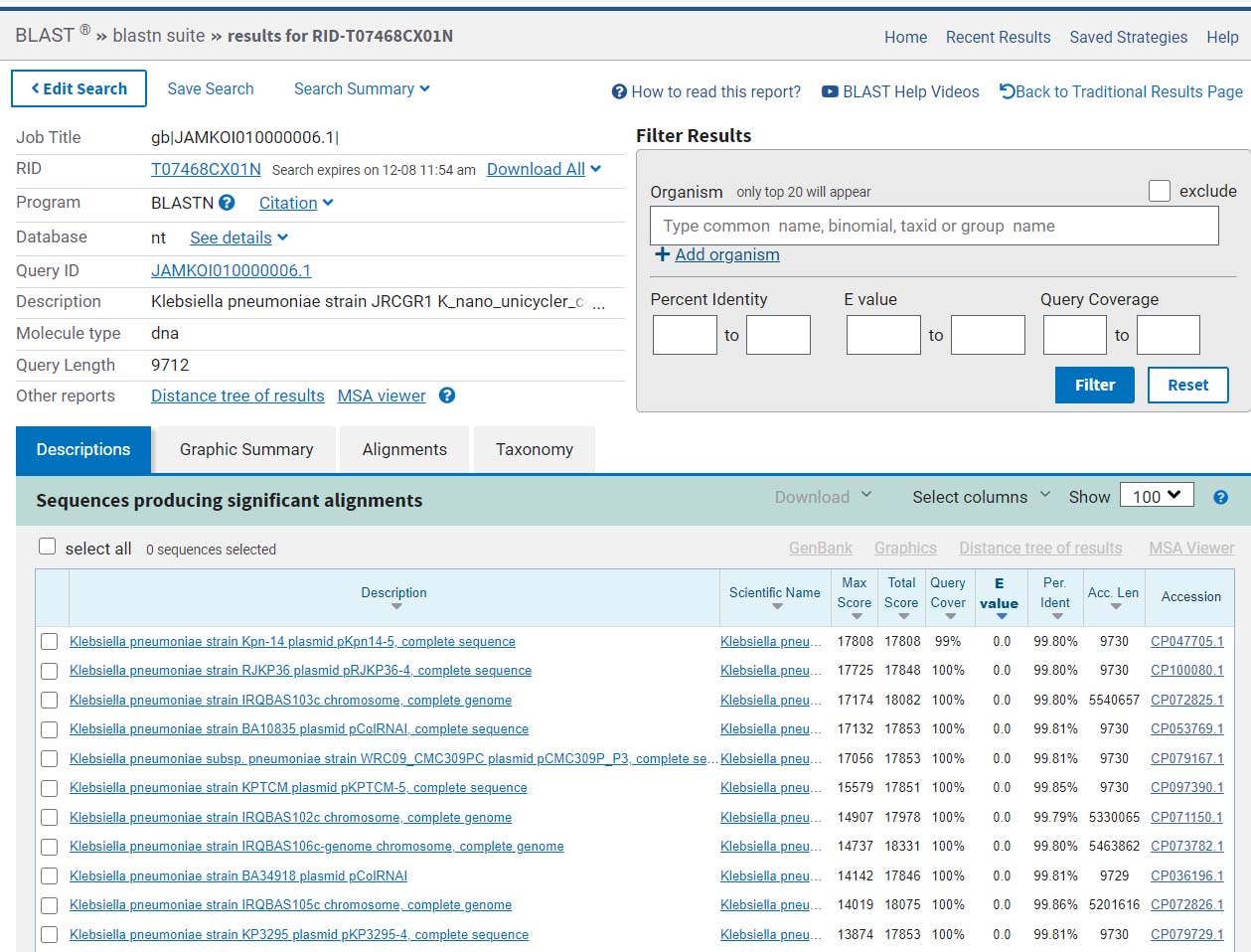

### Supplementary Fig. 1V. BLAST results confirming contig 6 as plasmid sequence.

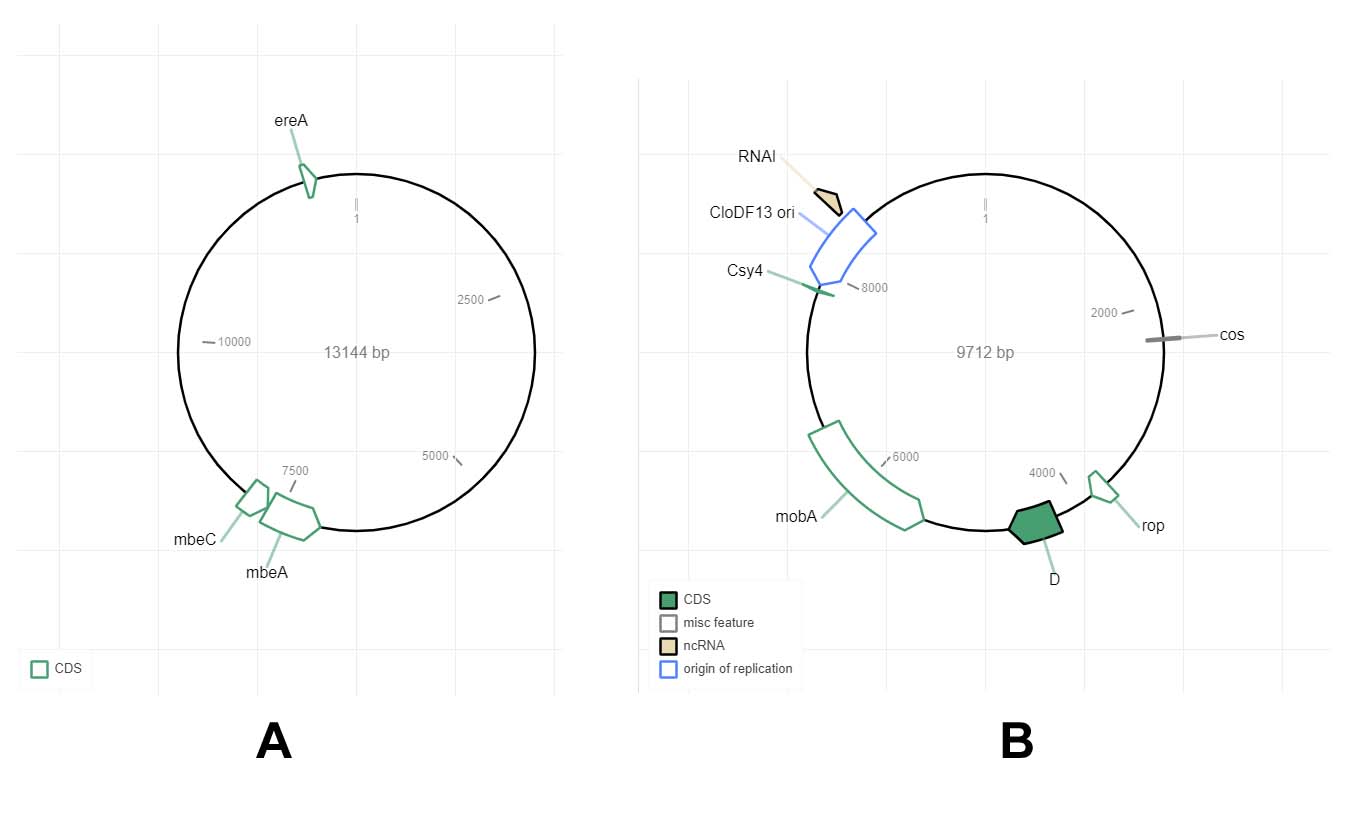

### Supplementary Fig. 2 (A) Circular plasmid map of contig 5 and (B) Circular plasmid map of contig 6.

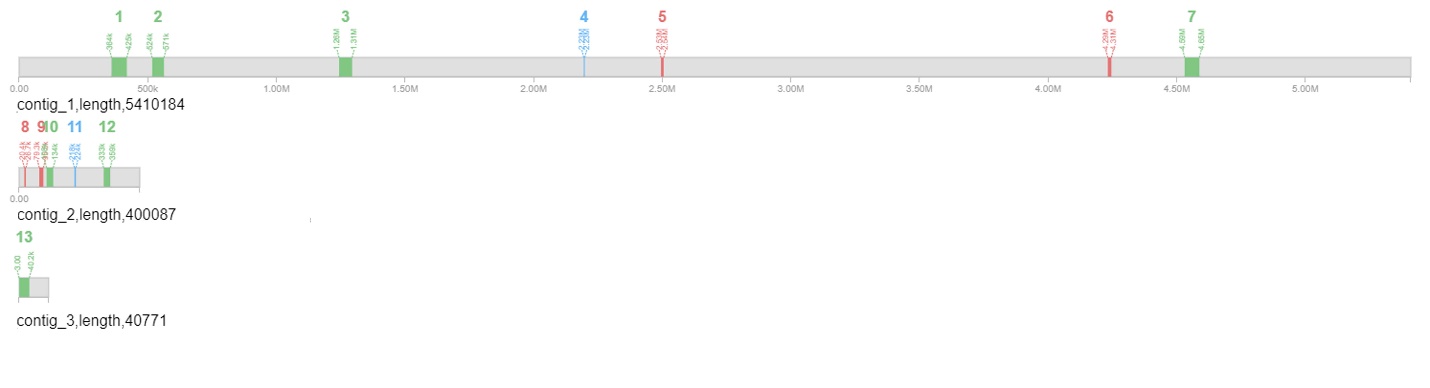

### Supplementary Fig. 3. Prophage regions identified using PHAST.
